# Pharmacological Characterisation of Novel Adenosine Receptor A_3_R Antagonists

**DOI:** 10.1101/693796

**Authors:** Kerry Barkan, Panagiotis Lagarias, Margarita Stampelou, Dimitrios Stamatis, Sam Hoare, Karl-Norbert Klotz, Eleni Vrontaki, Antonios Kolocouris, Graham Ladds

## Abstract

**Background and Purpose:** The adenosine A_3_ receptor (A_3_R) belongs to a family of four adenosine receptor (AR) subtypes which all play distinct roles throughout the body. A_3_R antagonists have been described as potential treatments for numerous diseases including asthma. Given the similarity between ARs orthosteric binding sites, obtaining highly selective antagonists is a challenging but critical task.

**Experimental approach:** 39 potential A_3_R, antagonists were screened using agonist-induced inhibition of cAMP. Positive hits were assessed for AR subtype selectivity through cAMP accumulation assays. The antagonist affinity was determined using Schild analysis (pA_2_ values) and fluorescent ligand binding. Further, a likely binding pose of the most potent antagonist (K18) was determined through molecular dynamic (MD) simulations and consistent calculated binding free energy differences between K18 and congeners, using a homology model of A_3_R, combined with mutagenesis studies.

**Key Results:** We demonstrate that K18, which contains a 3-(dichlorophenyl)-isoxazole group connected through carbonyloxycarboximidamide fragment with a 1,3-thiazole ring, is a specific A_3_R (<1 µM) competitive antagonist. Structure-activity relationship investigations revealed that loss of the 3-(dichlorophenyl)-isoxazole group significantly attenuated K18 antagonistic potency. Mutagenic studies supported by MD simulations identified the residues important for binding in the A_3_R orthosteric site. Finally, we introduce a model that enables estimates of the equilibrium binding affinity for rapidly disassociating compounds from real-time fluorescent ligand-binding studies.

**Conclusions and Implications:** These results demonstrate the pharmacological characterisation of a selective competitive A_3_R antagonist and the description of its orthosteric binding mode. Our findings may provide new insight for drug discovery.

**What is already known:**

- The search for AR subtype specific compounds often leads to ones with multiple subtype binding

**What this study adds:**

- This study demonstrates the pharmacological characterisation of a selective competitive A_3_R antagonist
- MD simulations identified the residues important for binding in the A_3_R orthosteric site

**Clinical significance:**

- This study offers insight into A_3_R antagonists that may provide new opportunities for drug discovery

## INTRODUCTION

The <u>adenosine A_3_ receptor</u> (A_3_R), belongs to a family of four <u>adenosine receptor</u> (AR) subtypes (<u>A_1_R</u>, <u>A_2A_R</u>, <u>A_2B_R</u> and A_3_R), and is involved in a range of pathologies including cardiovascular, neurological and tumour-related diseases. In particular, mast cell regulation and myocardial preconditioning are key physiological processes regulated by the A_3_R (Fredholm *et al.*, 2011). Unsurprisingly therefore, A_3_R is a pharmaceutical target. Interestingly, the A_3_R has been described as enigmatic, whereby many of the effects attributed to A_3_Rs are contradictory (Gessi *et al.*, 2008). Despite this, A_3_R antagonists having been described as potential treatments of asthma, chronic obstructive pulmonary disease (COPD) and glaucoma (Miwatashi *et al.*, 2008, Okamura *et al.*, 2004, Haeusler *et al*., 2015), to name a few, and continuous research into both agonists and antagonists at the A_3_R are warranted. While a number of novel potent and selective A_3_R antagonists have been previously described (Yaziji *et al.*, 2011, Yaziji *et al.*, 2013, Areias *et al.*, 2019), one of the challenges associated with the druggability of the AR family has been the targeting of individual subtypes with sufficient specificity to limit off-target side effects (Chen *et al.*, 2013).

Although all AR members are activated by the endogenous agonist <u>adenosine</u>, the A_2A_R and A_2B_R are predominantly G_s_-coupled whereas A_1_R and A_3_R generally couple to G_i/o_. This classical pathway following A_3_R activation and G_i/o_ coupling is the inhibition of adenylate cyclase (AC) resulting in a decrease in cAMP levels. Extracellular signal-regulated kinase 1/2 (ERK1/2) activation has also been described as downstream of A_3_R and is reported to be dependent on βγ-subunits released from pertussis toxin (PTX)-sensitive G_i/o_ proteins, phosphatidylinostitol-3-kinase (PI3K), the small GTP binding protein Ras, and MAP/ERK kinase (MEK) (Schulte and Fredholm, 2002).

The A_2A_R is one of the best structurally characterised G protein-coupled receptors (GPCRs), with multiple crystal structures available (Carpenter *et al.*, 2016, Lebon *et al.*, 2011, Lebon *et al.*, 2015, Xu *et al.*, 2011, Liu *et al.*, 2012, Doré *et al.*, 2011, Jaakola *et al.*, 2008, Cheng et al., 2017). Although the remaining AR subtypes have proven more difficult to crystallise with the A_3_R structure yet to be resolved, there are a number of A_1_R structures published (Draper-Joyce *et al.*, 2018, Glukhova *et al.*, 2017, Cheng *et al.*, 2017). Thus, *in silico* screening of vast compound libraries against receptor structures, known as structural-based drug design (SBDD), offers huge potential in the development of highly selective ligands (Carlsson *et al*., 2010, Katritch *et al*., 2010, Lenselink *et al.*, 2016, Lagarias *et al.*, 2018). The limited availability of diverse high-resolution structures of the remaining AR subtypes bound to pharmacologically distinct ligands has meant there is a discrepancy between the capability to predict compound binding versus pharmacological behaviour; partial agonism, inverse agonism, biased agonist, antagonism, allosteric modulation etc (Sexton and Christopoulos, 2018). With this in mind, the potential antagonists (K1-K25, K28 and K35) identified in our previously published virtual screening investigation and binding experiments (Lagarias *et al.*, 2018) and some newly identified potential antagonists (K26, K27, K29-K34 and K36-K39) were pharmacologically characterised using cAMP accumulation. We were able to identify a potent and selective A_3_R antagonist, K18 (O4-{[3-(2,6-dichlorophenyl)-5-methylisoxazol-4-yl]carbonyl}-2-methyl-1,3-thiazole-4-carbohydroximamide) and, using molecular dynamic (MD) simulations combined with site-directed mutagenesis, elude to the potential binding site. Binding free energy calculations of similar in structure analogs of K18 were consistent with the proposed A_3_R orthosteric binding area. Kinetic binding experiments of K5, K17 and K18 using a bioluminescence resonance energy transfer (BRET) method combined with functional assays led to the identification of important structural features of K18 for binding and activity. Further evaluation of this compound (and structurally related analogues) may afford a novel therapeutic benefit in pathologies such as inflammation and asthma.

## RESULTS

### Identification of A_3_R selective antagonists

We initially conducted a blinded screen of 39 compounds (K1-39) to identify selective A_3_R antagonists some of which have previously been identified to bind A_1_R, A_3_R or A_2A_R (Lagarias *et al.*, 2018). Our screen was carried out using A_3_R expressing Flp-In™-Chinese hamster ovary (CHO) cells where cAMP accumulation was detected following a combined stimulation of 10 μM forskolin (to allow A_3_R mediated G_i/o_ response to be observed), 1 μM tested compound and the predetermined IC_80_ concentration of NECA (3.16 nM). Compound K1-39 were identified by unblinding (Table 1 and Supplementary Table 1) but are hereinafter referred to as their denoted ‘K’ number. For the purpose of structure-activity relationships studies, the previously uncharacterised compounds (K26, K27, K29-K34 and K36-K39), were assayed both functionally and through radioligand binding (Supplementary Table 1).

**Table 1.**
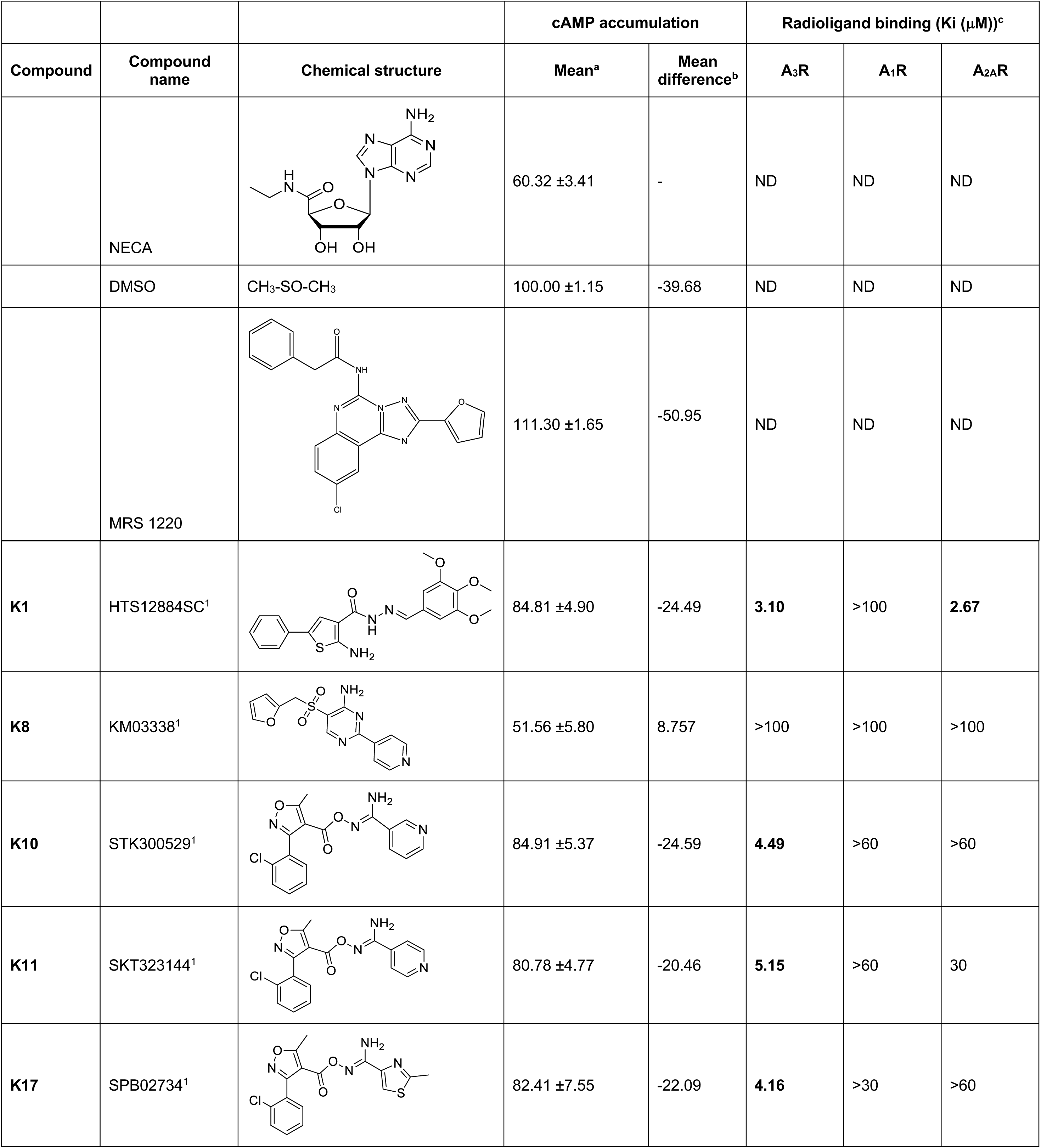
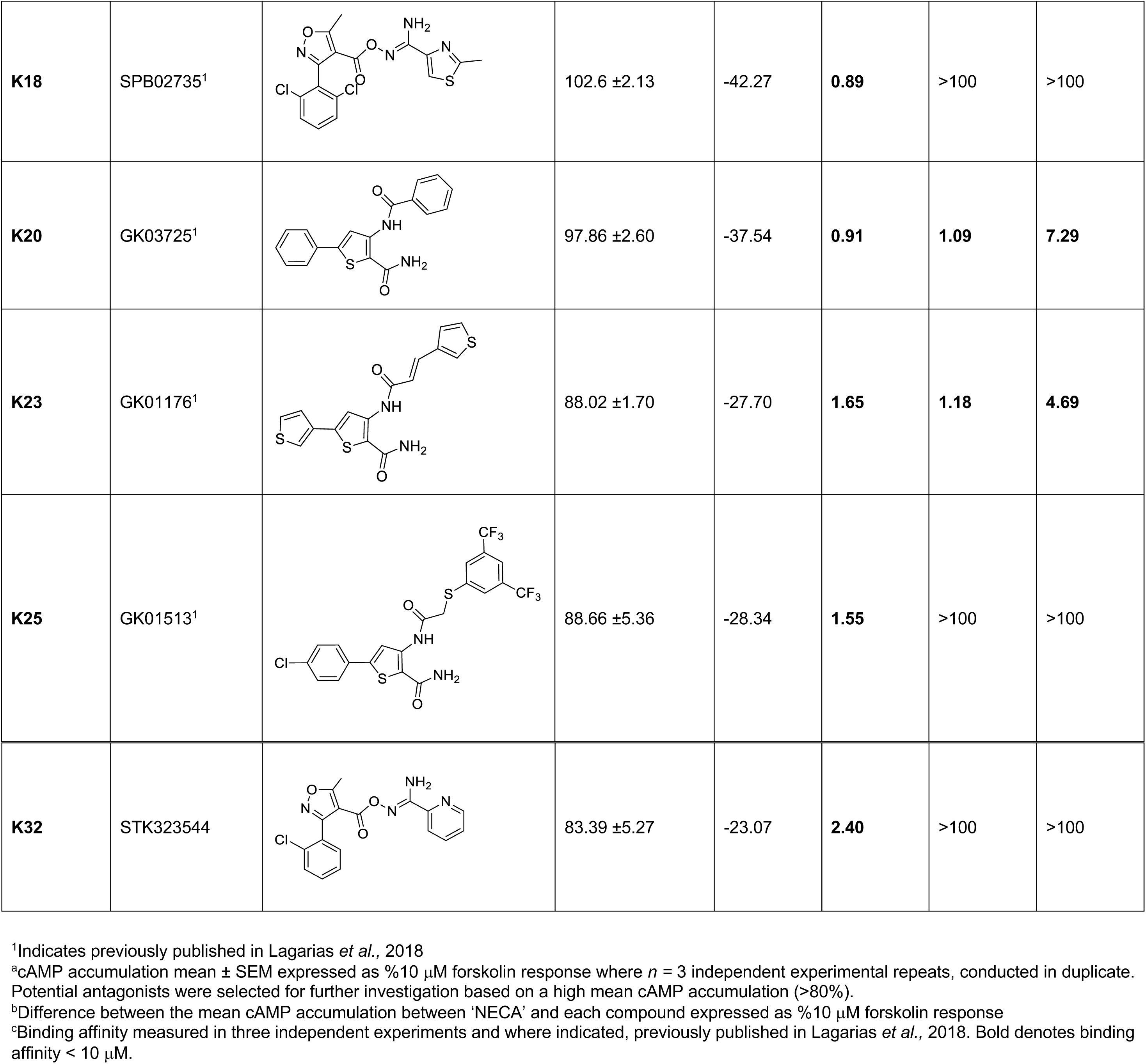
Mean cAMP accumulation as measured in Flp-In CHO cells stably expressing A_3_R following stimulation with 10 μM forskolin only (DMSO) or 10 μM forskolin, NECA at the predetermined IC_80_ concentration and 1 μM test compound/MRS 1220/DMSO control. Binding affinities were obtained through radioligand binding assays against the A_1_R, A_2A_R and A_3_R.

| Compound | Compound name | Chemical structure | cAMP accumulation |  | Radioligand binding (K<sub>i</sub> (μM))<sup>c</sup> |  |  |
| --- | --- | --- | --- | --- | --- | --- | --- |
|  |  |  | Mean<sup>a</sup> | Mean difference<sup>b</sup> | A<sub>3</sub>R | A<sub>1</sub>R | A<sub>2A</sub>R |
|  | NECA |  | 60.32 ±3.41 | - | ND | ND | ND |
|  | DMSO | CH<sub>3</sub>-SO-CH<sub>3</sub> | 100.00 ±1.15 | -39.68 | ND | ND | ND |
|  | MRS 1220 |  | 111.30 ±1.65 | -50.95 | ND | ND | ND |
| **K1** | HTS12884SC<sup>1</sup> |  | 84.81 ±4.90 | -24.49 | **3.10** | >100 | **2.67** |
| **K8** | KM03338<sup>1</sup> |  | 51.56 ±5.80 | 8.757 | >100 | >100 | >100 |
| **K10** | STK300529<sup>1</sup> |  | 84.91 ±5.37 | -24.59 | **4.49** | >60 | >60 |
| **K11** | SKT323144<sup>1</sup> |  | 80.78 ±4.77 | -20.46 | **5.15** | >60 | 30 |
| **K17** | SPB02734<sup>1</sup> |  | 82.41 ±7.55 | -22.09 | **4.16** | >30 | >60 |
| K18 | SPB02735<sup>1</sup> |  | 102.6 ±2.13 | -42.27 | **0.89** | >100 | >100 |
| K20 | GK03725<sup>1</sup> |  | 97.86 ±2.60 | -37.54 | **0.91** | **1.09** | **7.29** |
| K23 | GK01176<sup>1</sup> |  | 88.02 ±1.70 | -27.70 | **1.65** | **1.18** | **4.69** |
| K25 | GK01513<sup>1</sup> |  | 88.66 ±5.36 | -28.34 | **1.55** | >100 | >100 |
| **K32** | STK323544 |  | 83.39 ±5.27 | -23.07 | **2.40** | >100 | >100 |

<sup>1</sup>Indicates previously published in Lagarias *et al.*, 2018
<sup>a</sup>cAMP accumulation mean ± SEM expressed as %10 µM forskolin response where *n* = 3 independent experimental repeats, conducted in duplicate. Potential antagonists were selected for further investigation based on a high mean cAMP accumulation (>80%).
<sup>b</sup>Difference between the mean cAMP accumulation between 'NECA' and each compound expressed as %10 µM forskolin response
<sup>c</sup>Binding affinity measured in three independent experiments and where indicated, previously published in Lagarias *et al.*, 2018. Bold denotes binding affinity < 10 µM.

Co-stimulation with 10 μM of both forskolin and NECA reduced the cAMP accumulation when compared to 10 μM forskolin alone and this was reversed with the known A_3_R antagonist <u>MRS 1220</u> (Table 1 and Supplementary Fig 1). Compounds K1, K10, K11, K17, K18, K20, K23, K25 and K32 were identified as potential antagonists at the A_3_R through their ability to elevate cAMP accumulation when compared to forskolin and NECA co-stimulation. Of the nine potential A_3_R antagonists, eight (excluding K11) appeared to be antagonists at the tested concentration of 1 μM (Supplementary Fig. 2 and Supplementary Table 2).

A number of compounds previously documented (K5, K9, K21, K22 and K24; Lagarias *et al.*, 2018) or determined in this study (K26, K27 and K34) to have sub-micromolar binding affinities for A_3_R showed no activity in our cAMP-based screen (Table 1, Supplementary Table 1). To ensure robustness of our functional screen, full inhibition curves of NECA in the presence or absence of tested compounds (1 μM or 10 μM) were constructed in A_3_R Flp-In CHO cells (Supplementary Fig. 3, Supplementary Table. 3). In this preliminary data all nine compounds (K5, K9, K11, K21, K22, K24, K26, K27 and K34) appeared to reduce the NECA potency at the highest tested concentration (10 μM) but showed no effect at 1 μM and thus appear to be low potency antagonists at the A_3_R.

### AR subtype selectivity and specificity

The similarity of the different ARs has meant many compounds display reduced selectivity. Using the A_3_R Flp-In CHO or CHO-K1 cells expressing A_1_R, A_2A_R or A_2B_R incubated with a single high concentration of antagonist (10 μM) and increasing concentrations of NECA identified K10, K17, K18 and K25 as A_3_R selective antagonists (Fig. 1). K20 and K23 were antagonists at both the A_1_R and A_3_R (Fig. 1 and Table 2). K1, K20 and K23 showed weak antagonism at the A_2A_R and none of the tested antagonist showed any antagonism of the NECA stimulated response at the A_2B_R. These selectivity findings agree with our previously published radioligand binding data (Lagarias *et al.*, 2018) and are summarised in Table 2.

**Table 2.** Potency of NECA stimulated cAMP inhibition or accumulation as determined in Flp-In CHO or CHO-K1 cells expressing one of four ARs subtype (A_3_R, A_1_R, A_2A_R or A_2B_R). Cells stably expressing A_3_R, A_1_R, A_2A_R or A_2B_R were stimulated with 10 μM forskolin (in the case of A_3_R and A_1_R), 10 μM tested compound/DMSO and increasing concentrations of NECA.

|  | pIC<sub>50</sub>/pEC<sub>50</sub><sup>a</sup> |  |  |  |
| --- | --- | --- | --- | --- |
|  | A<sub>3</sub>R | A<sub>1</sub>R | A<sub>2A</sub>R | A<sub>2B</sub>R |
| **NECA only** | 9.24 ± 0.1 | 8.98 ± 0.1 | 7.88 ± 0.1 | 7.24 ± 0.2 |
| **K1** | 8.01 ± 0.2**** | 8.97 ± 0.1 | 7.12 ± 0.1**** | 7.23 ± 0.2 |
| **K10** | 7.74 ± 0.2**** | 8.82 ± 0.1 | 7.84 ± 0.1 | 7.19 ± 0.2 |
| **K17** | 7.59 ± 0.1**** | 8.68 ± 0.1 | 7.76 ± 0.1 | 7.15 ± 0.2 |
| **K18** | 6.70 ± 0.1**** | 8.85 ± 0.1 | 7.75 ± 0.1 | 7.10 ± 0.2 |
| **K20** | 7.12 ± 0.2**** | 7.43 ± 0.1 **** | 7.12 ± 0.1**** | 7.08 ± 0.1 |
| **K23** | 7.72 ± 0.1**** | 7.38 ± 0.1 **** | 7.26 ± 0.1** | 7.04 ± 0.2 |
| **K25** | 7.64 ± 0.1**** | 9.00 ± 0.1 | 7.98 ± 0.1 | 7.22 ± 0.2 |
| **K32** | 7.56 ± 0.1**** | 8.85 ± 0.1 | 7.80 ± 0.1 | 7.14 ± 0.2 |

<sup>a</sup>Negative logarithm of NECA concentration required to produce a half-maximal response in the absence (NECA only) or presence of 10 μM compound at each AR subtype
Data are expressed as mean ± SEM obtained in $n = 5$ independent experimental repeats, conducted in duplicate. Statistical significance (\*, $p < 0.05$ ; \*\*, $p < 0.01$ ; \*\*\*, $p < 0.001$ ; \*\*\*\*, $p < 0.0001$ ) compared to 'NECA only' was determined by one-way ANOVA with Dunnett's post-test.

**Figure 1.**
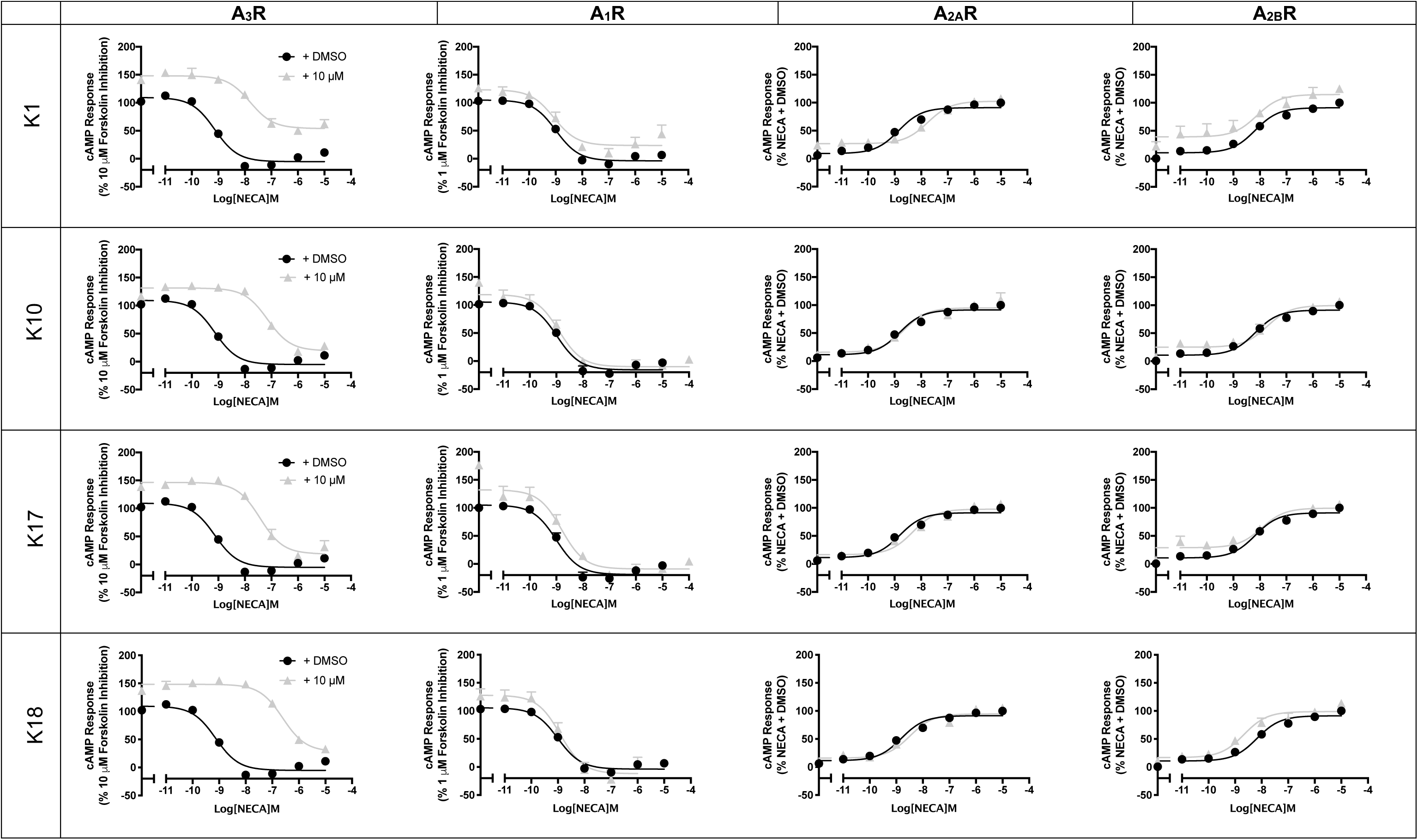
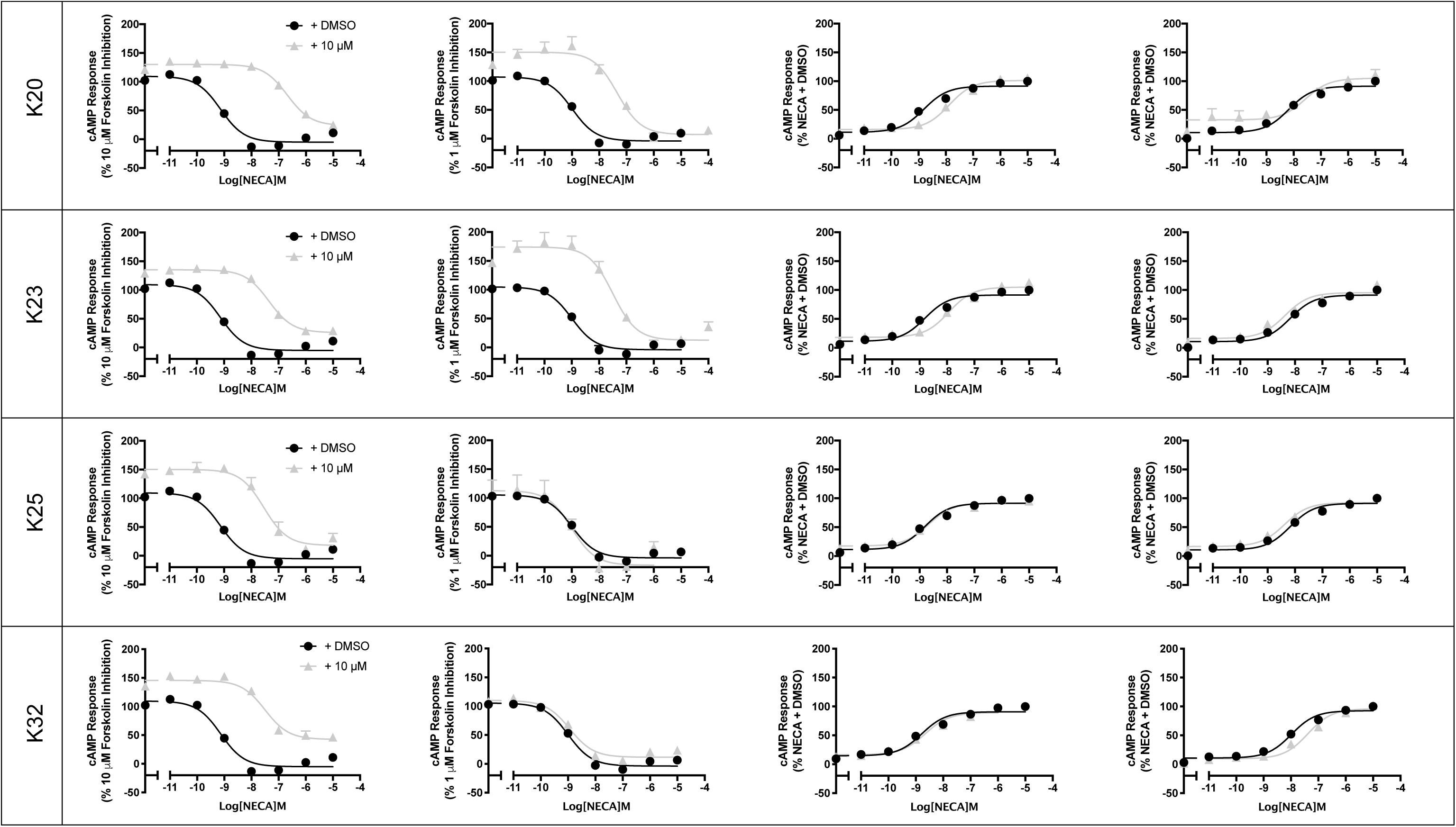
Characterisation of A_3_R antagonist at all AR subtypes. A_3_R Flp-In CHO cells or CHO-K1 cells (2000 cells/well) stably expressing one of the remaining AR subtypes were exposed to forskolin in the case of G_i_-coupled A_1_R and A_3_R (1 μM or 10 μM, respectively) or DMSO control in the case of G_s_-coupled A_2A_R and A_2B_R, NECA and test compound (10 μM) for 30 min and cAMP accumulation detected. All values are mean ± SEM expressed as percentage forskolin inhibition (A_1_R and A_3_R) or stimulation (A_2A_R and A_2B_R), relative to NECA. *n* ≥ 3 independent experimental repeats, conducted in duplicate.

### Characterisation of competitive antagonists at the A_3_R

All eight A_3_R antagonists were confirmed to antagonise <u>IB-MECA</u> (Fig. 2 and Table 3) and preliminary data suggests this extends to NECA antagonism (Supplementary Fig. 4 and Supplementary Table 4) in a concentration-dependent manner. Schild analysis characterised K10, K17, K18, K20, K23, K25 and K32 as competitive antagonists at the A_3_R (Schild slope not significantly different from unity, Fig. 2). Interestingly, the Schild slope deviated from unity for K1 (in competition experiments with NECA, but not IB-MECA) suggesting a more complicated mechanism of antagonism at the A_3_R. K20 and K23 were also characterised as competitive antagonists at the A_1_R (Supplementary Fig. 5 and Supplementary Table 5).

**Table 3.**
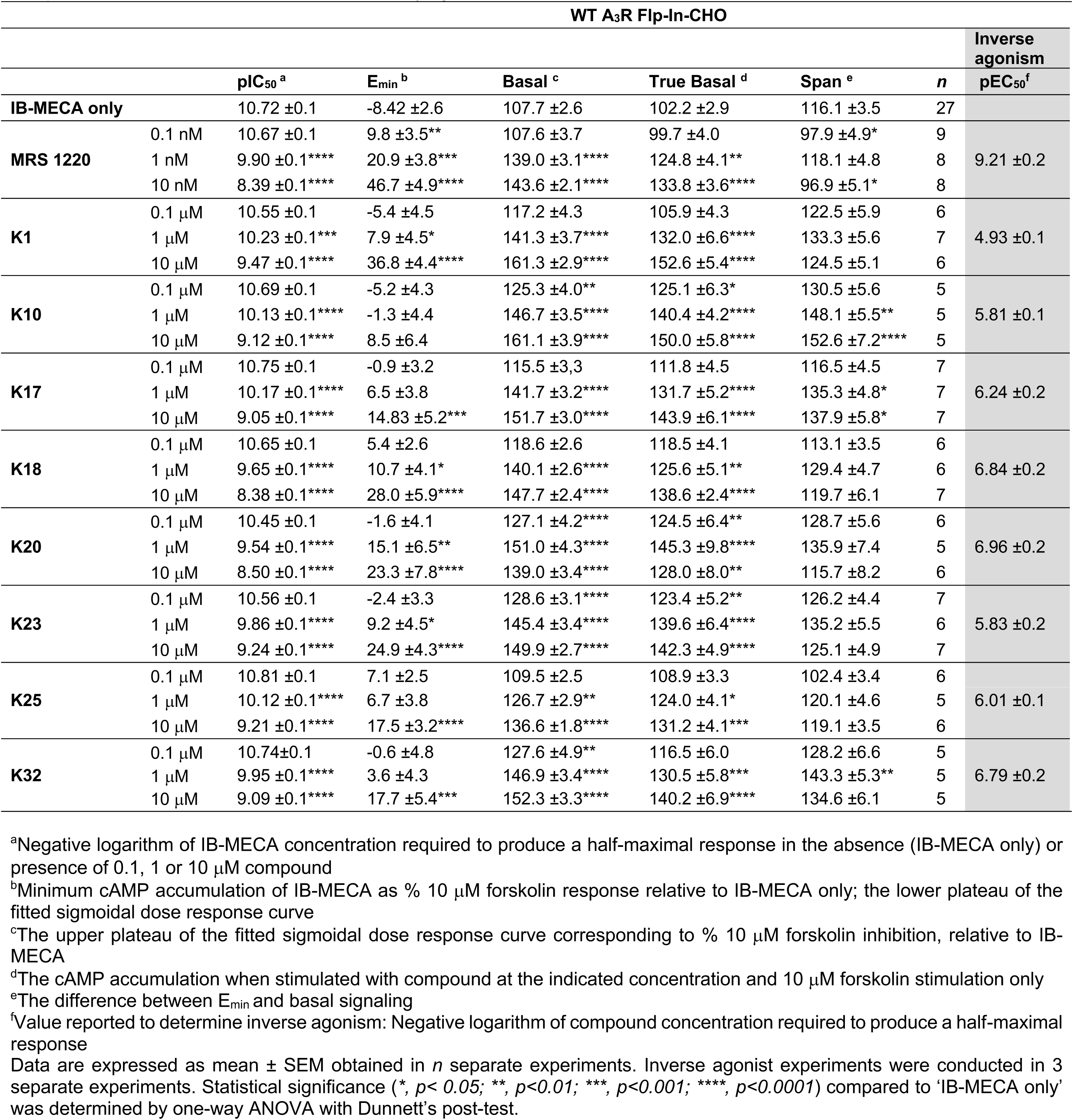
IB-MECA stimulated cAMP inhibition at WT A_3_R: activity of MRS 1220 and potential antagonists. Forskolin stimulated cAMP inhibition was measured in Flp-In-CHO stably expressing A_3_R following stimulation with 10 μM forskolin, compound at the indicated concentration and varying concentrations of IB-MECA.

**Figure 2.**
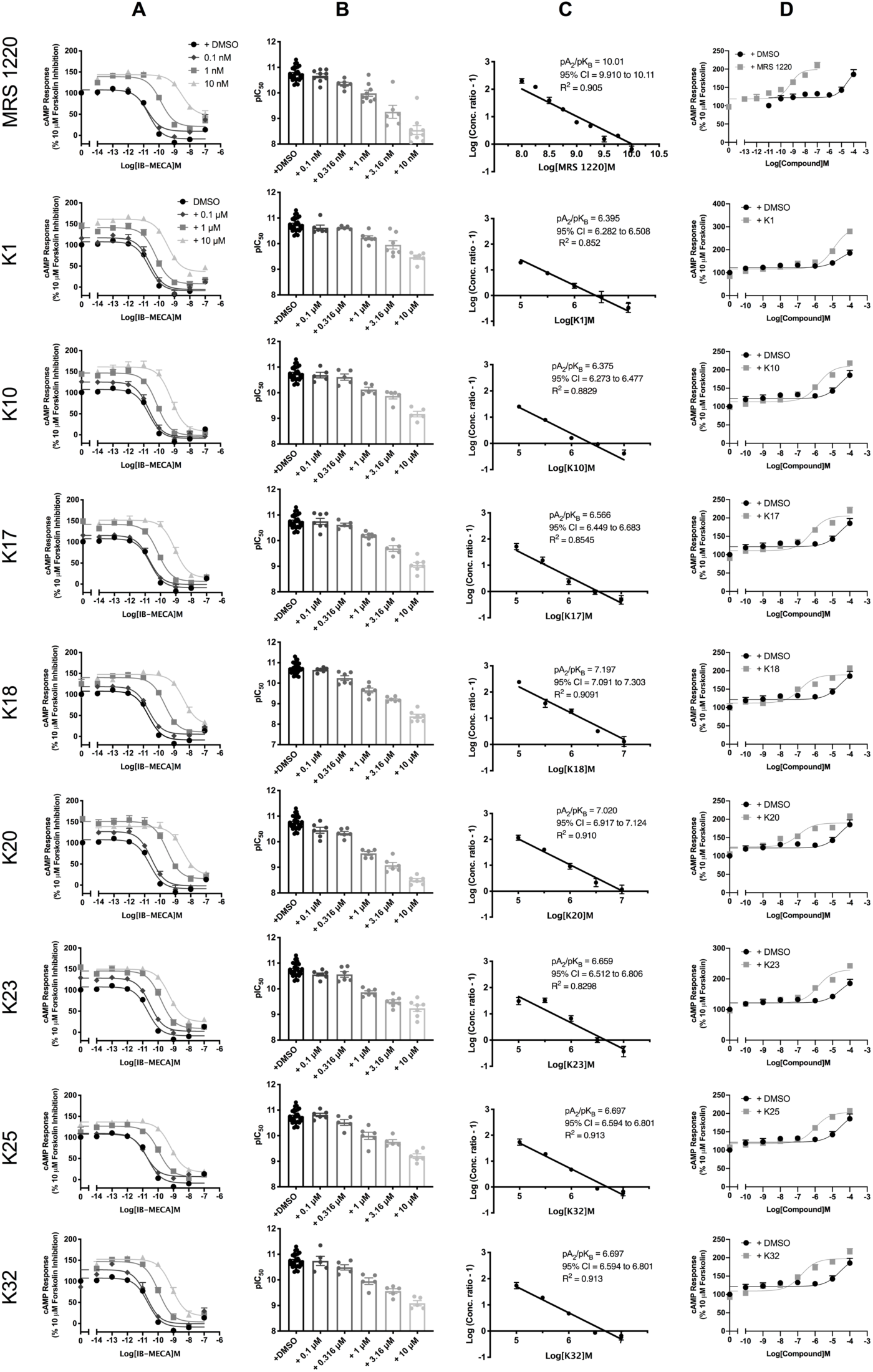
IB-MECA stimulated cAMP inhibition at WT A_3_R: activity of MRS 1220 and potential antagonists. Flp-In-CHO cells (2000 cells/well) stably expressing WT A_3_R were exposed to forskolin 10 μM, IB-MECA and test compound/MRS 1220/DMSO control for 30 min and cAMP accumulation detected. **A**) Representative dose response curves are shown as mean ± SEM expressed as percentage forskolin inhibition (10 μM) relative to IB-MECA. Key indicated in K1 is identical for all ‘K’ test compounds shown. **B**) pIC_50_ values for individual repeats including half-log concentration are shown as mean ± SEM **C**) Schild analysis of data represented in **A/B**. A slope of 1 indicates a competitive antagonist. The x-axis is expressed as -log (molar concentration of antagonist) giving a negative Schild slope. **D**) Inverse agonism at the A_3_R. cAMP accumulation following a 30-minute stimulation with forskolin (10 μM) and increasing concentrations of antagonist/DMSO control was determined in WT A_3_R expressing Flp-In-CHO cells. Representative dose response curves are shown as mean ± SEM expressed as percentage forskolin (10 μM), relative to IB-MECA.

When comparing the activity of A_3_R selective antagonists (K10, K17, K18 and K25), K18 was the most potent, showed A_3_R specificity and greater A_3_R binding affinity (Table 2) and we propose it as our lead compound. The original competition binding experiments that identified the panel of antagonist was performed using [^3^H]HEMADO (Lagarias *et al.*, 2018). To ensure that the different ligands used in our studies was not influencing our characterisation of the compounds, we assessed the ability of K18 to antagonise cAMP inhibition by HEMADO at the A_3_R and compared its potency to K17 (Supplementary Fig. 6 and Table 6). In this exploratory data K18 again displayed higher potency than K17 at the A_3_R.

In addition, we wanted to determine if K18 could also antagonise the activity of the A_3_R when an alternative downstream signalling component was measured; ERK1/2 phosphorylation (Fig. 3). In line with previously reported findings (Graham *et al.*, 2001; Schulte and Fredholm, 2002), agonists at the A_3_R increased ERK1/2 phosphorylation after 5 minutes, with IB-MECA 10-fold more potent than NECA (Supplementary Fig. 7) and preliminary data suggests this was entirely G_i/o_-mediated (pERK1/2 levels were abolished upon addition of PTX). K18 was able to antagonise A_3_R-mediated phosphorylation of ERK1/2 with the antagonist potency (pA_2_ values) not significantly different compared to the cAMP-inhibition assay (Fig. 3C).

**Figure 3.**
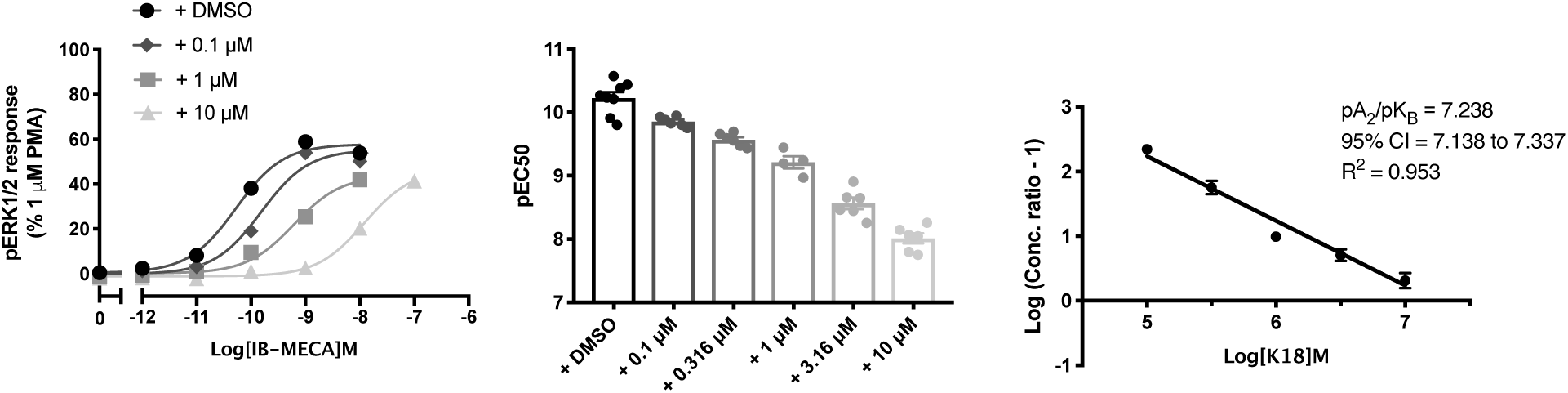
K18 also reduced levels of agonist stimulated ERK1/2 phosphorylation. pERK1/2 was detected in Flp-In-CHO cells stably expressing A_3_R (2000 cells/well) stimulated for 5 minutes with IB-MECA, with or without K18. **A)** Representative dose response curves for IB-MECA with K18 at the indicated concentration or DMSO control shown as mean ± SEM expressed as % 1μM PMA response. **B**) pEC_50_ values for individual repeats are shown as mean ± SEM**. C**) Schild analysis of data represented in **A/B**.

### A_3_R constitutive activity and inverse agonism

We next determined if any of our competitive antagonist could function as inverse agonists of the A_3_R. In our hands, the A_3_R, when expressed in Flp-In™-CHO cells, displays constitutive activity (Supplementary Fig. 8). All eight characterised A_3_R antagonists showed a concentration-dependent inverse agonism of the A_3_R when compared to DMSO control (Fig. 2). This was also found to be the case for <u>DPCPX</u>, K20 and K23 at the A_1_R (Supplementary Fig. 9). Notably, DMSO showed a concentration-dependent elevation in cAMP accumulation above that of forskolin alone.

### MD simulation of the binding mode of K18 at A_3_R

We next sought to investigate the potential binding pose of K18 within the A_3_R orthosteric site. Building upon our previous studies where we have generated a homology model of the A_3_R, K18 was docked into the orthosteric site of the A_3_R using the GoldScore scoring function and the highest scoring pose was inserted in a hydrated POPE bilayer. The complex was subjected to MD simulations in the orthosteric binding site of A_3_R with Amber14ff for 100 ns and the trajectory analysed for protein-ligand interactions. We identified a potential binding pose of K18 within the established orthosteric A_3_R binding pocket (Fig. 4). The MD simulations suggest that K18 forms hydrogen bonds, van der Waals (vdW) and π-π interactions inside the orthosteric binding site of A_3_R (Fig. 4A). More specifically, MD simulations showed that the 3- (dichlorophenyl) group is positioned close to V169^5.30^, M177^5.40^, I249^6.54^ and L264^7.34^ of the A_3_R orthosteric binding site forming attractive vdW interactions. The isoxazole ring is engaged in an aromatic *π-π* stacking interaction with the phenyl group of F168^5.29^ (Fig. 4A). The thiazole ring is oriented deeper into the receptor favouring interactions with L246^6.51^, L90^3.32^ and I268^7.39^. Hydrogen bonding interactions can be formed between: (a) the amino group of the carbonyloxycarboximidamide molecular segment and the amide side chain of N250^6.55^; (b) the nitrogen or the sulphur atom of the thiazole ring and N250^6.55^ side chain (Fig. 4A). For structural comparison and insight, we also modelled K5 and K17 binding at the A_3_R given the structural similarity: K17 and K18 possess one and two chlorine atoms attached to the phenyl ring of the phenyl-isoxazole system, respectively, whereas K5 has none (Fig. 4B and C).

**Figure 4.**
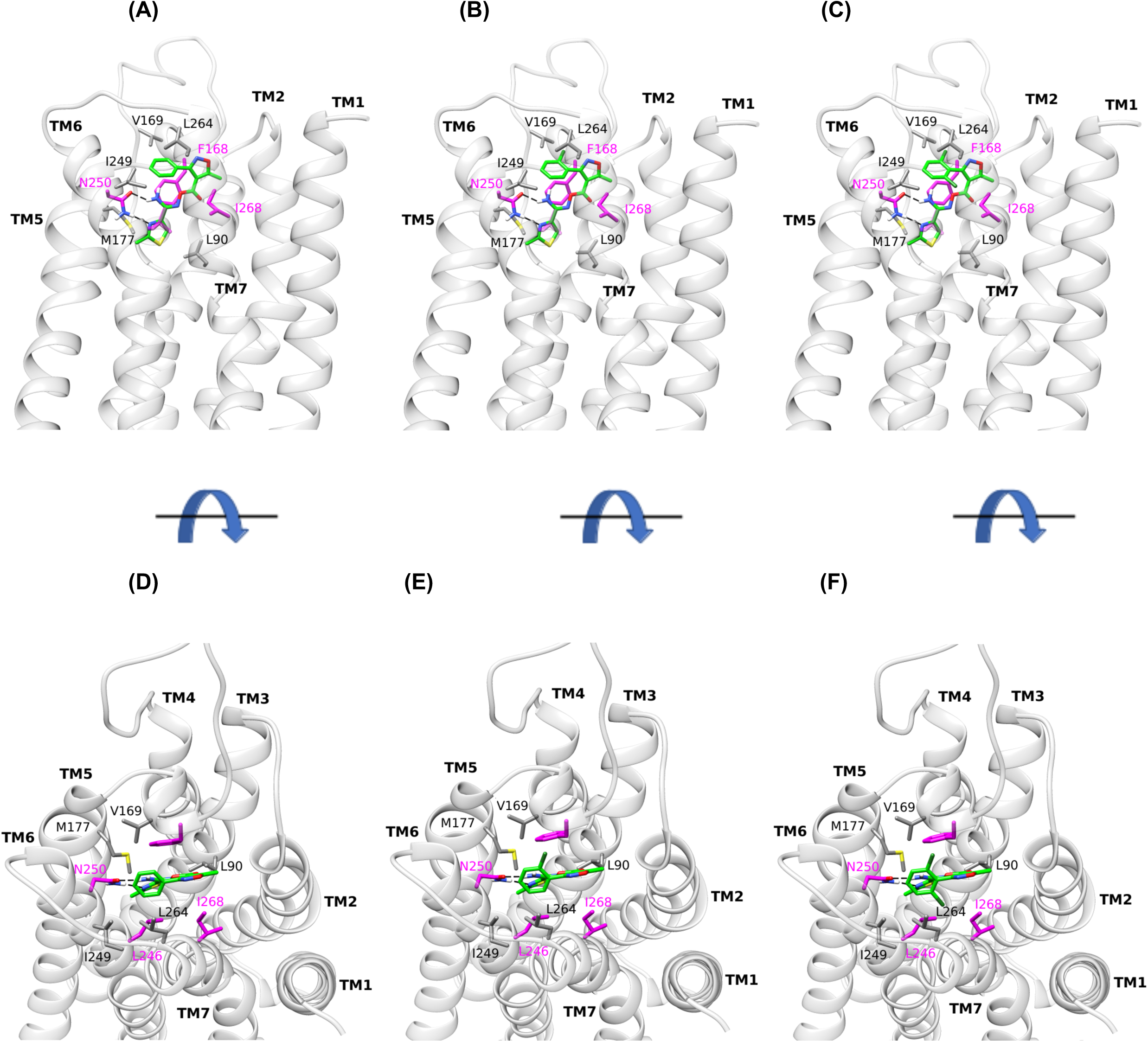
Orthosteric binding area average structure of WT A_3_R in complex with K5, K17 and K18 from MD simulations with Amber14ff. Side **(A)**, top **(D)** view of K5 complex; side **(B)**, top **(E)** view of K17 complex; side **(C)**, top **(F)** view of K18 complex. Side chains of critical residues for binding indicated from the MD simulations are shown in sticks. Residues L90^3.32^, V169^5.30^, M177^5.40^, I249^6.54^ and L264^7.34^, in which carbon atoms are shown in grey, were confirmed experimentally; in residues F168^5.29^, L246^6.51^, I268^7.39^ and N250^6.55^ carbon atoms are shown in magenta; nitrogen, oxygen and sulfur atoms are shown in blue, red and yellow respectively.

### Molecular Mechanics-Poisson Boltzmann Surface Area (MM-PBSA) calculations validate binding pose of K18

We next applied the MM-PBSA method in the MD simulation trajectories of the compounds to calculate their binding free energies (Δ*G*_eff_ -see methods for derivation (Stamatis *et al.*, 2019)) and evaluated the energetic contributions for their binding (Table 4). The calculated ranking in the binding free energies were in agreement with experimental differences in potencies (K5 < K17 < K18 < MRS 1220). The known A_3_R antagonist MRS 1220 displayed the lowest Δ*G*_eff_ value, which is also in agreement with the experimental potencies. The calculations suggested that the major difference between the energetic components of Δ*G*_eff_ values for K5, K17 and K18 is on the solvation energies (Δ*G*_solv_). The two chlorine atoms make K18 more lipophilic and reduce the energy required to transfer the compound from solution to the binding area, increasing the free energy of binding and activity compared to K17 and K5.

**Table 4.** Binding of K5, K17, K18 and MRS 1220 to the A_3_R orthosteric binding area. Effective binding energies (Δ*G*_eff_) and energy components (*E*_vdW_, *E*_EL_, Δ*G*_solv_) in kcal mol^-1^ calculated using the MM-PBSA method.

| | $E_{\text{vdW}}^a$ | $E_{\text{EL}}^b$ | $\Delta G_{\text{solv}}^c$ | $\Delta G_{\text{eff}}^d$ | $pK_B/pK_i^e$ | | |
| --- | --- | --- | --- | --- | --- | --- | --- |
|  |  |  |  |  | Schild analysis<sup>f</sup> | NanoBRET<sup>g</sup> | Radioligand binding<sup>h</sup> |
| MRS 1220 | -64.6 ± 2.6 | -11.5 ± 2.5 | 39.2 ± 2.4 | -36.9 ± 3.6 | 10.07 | 9.99 ± 0.04 | 8.2 - 9.2 |
| K5 | -42.0 ± 2.7 | -9.6 ± 5.2 | 30.8 ± 4.3 | -20.8 ± 4.3 | ND | 6.06 ± 0.09 | 5.02 |
| K17 | -47.0 ± 2.4 | -8.8 ± 2.7 | 29.8 ± 2.9 | -25.9 ± 3.6 | 6.35 | 6.33 ± 0.03 | 5.38 |
| K18 | -46.3 ± 2.9 | -7.5 ± 2.4 | 26.9 ± 3.1 | -26.9 ± 2.7 | 7.20 | 6.92 ± 0.10 | 6.05 |

<sup>a</sup>vdW energy of binding calculated using molecular mechanics
<sup>b</sup>Electrostatic energy of binding calculated using molecular mechanics
<sup>c</sup>Difference in solvation energy between the complex, the protein and the ligand, i.e. $G_{\text{complex, solv}} - (G_{\text{protein, solv}} + G_{\text{ligand, solv}})$
<sup>d</sup>Effective binding free energy calculated as $\Delta G_{\text{eff}} = \Delta E_{\text{MM}} + \Delta G_{\text{solv}}$ ; in Table 4, $\Delta E_{\text{MM}} = E_{\text{vdW}} + E_{\text{EL}}$ (see Materials and Methods)
<sup>e</sup>Equilibrium dissociation constant of MRS 1220, K5, K17 and K18 as determined through three independent experimental approaches: Schild analysis ( $pK_B$ ), NanoBRET ( $pK_i$ ) or radioligand binding ( $pK_i$ )
<sup>f</sup> $pK_B$ obtained through Schild analysis in A<sub>3</sub>R stably expressing Flp-In CHO cells
<sup>g</sup> $pK_i$ (mean ± SEM) obtained in NanoBRET binding assays using Nluc-A<sub>3</sub>R stably expressing HEK 293 cells and determined through fitting our “Kinetics of competitive binding, rapid competitor dissociation” model or in the case of MRS 1220 through fitting with the ‘Kinetics of competitive binding’ model with a determined $K_{\text{on}}$ ( $k_3$ ) and $K_{\text{off}}$ ( $k_4$ ) rate of $3.25 \pm 0.28 \times 10^8 \text{ M}^{-1} \text{ min}^{-1}$ and $0.0248 \pm 0.005 \text{ min}^{-1}$ , respectively
<sup>h</sup> $pK_i$ values previously published for K5, K17 and K18 (Lagarias *et al.*, 2018) or MRS 1220 (Stoddart *et al.*, 2015) through radioligand binding assays.

### MD simulation of the binding mode of analogs of K18 at A_3_R

Significantly, when the 4-thiazolyl in K17 was changed to 2-,3- or 4-pyridinyl in compounds K32, K10, K11 we observed antagonistic activity only for compounds K32 and K10 (not K11) (Table 1) since the pyridine nitrogen in compounds K32 and K10 can interact with N250^6.55^ due to their proximate positions in binding conformation (see Fig. 4B for K17). This interaction is preserved with both the 2-,3-pyridinyl groups but lost when the nitrogen of the pyridinyl group is in the 4-position. In addition, following MD simulations of compounds K26, K27, K29-K34 and K36-K39, we observed that K26 (Supplementary Fig. 10) displayed a similar binding pose to that of K18 (Fig. 4). However, K26 (and K34 for that matter) which containing a o-diphenylcarbonyl (also present in K18) functionally showed weak antagonistic potency (> 1 μM, Supplementary Fig. 2) suggesting a more complex binding mode is present. We also observed that compared to K26 and K34, the p-substitution in compounds K29 and K36-38 was not favourable for binding at all since this led to a loss of the vdW interaction with the hydrophobic area of the A_3_R towards TM5 and TM6; as was demonstrated in MD simulations for K36 (Supplementary Fig. 10).

### Experimental evaluation of the binding mode of K18 at A_3_R

Of the identified residues predicted to mediate an interaction between K18 and the A_3_R, the ones which showed (according to the MD simulations) the most frequent and the most important contacts were chosen for investigation and included amino acids L90^3.32^, F168^5.29^, V169^5.30^, M177^5.40^, L246^6.51^, I249^6.54^, N250^6.55^, L264^7.34^ and I268^7.39^ (Fig. 4). Site-directed mutagenesis was performed replacing each residue with an alanine in the A_3_R and expressed in the Flp-In-CHO™ cells lines. Each mutant was then screened for their ability to suppress forskolin-induced cAMP accumulation in response to NECA/IB MECA stimulation in the presence and absence of K18.

Mutation of residues F168^5.29^, L246^6.51^, N250^6.55^ and I268^7.39^ abolished agonist induced suppression of forskolin-induced cAMP accumulation and were discontinued in this study (Stamatis *et al.*, 2019). Both L90A^3.32^ and M177A^5.40^ showed a significantly decreased NECA and IB-MECA potency. L264A^7.34^ showed a slight decrease in IB-MECA potency whereas the potency of NECA was similar to WT. Whereas the NECA stimulated cAMP inhibition in V169A^5.30^ or I249A^6.54^ expressing Flp-In CHOs was comparable to WT, the IB-MECA stimulated cAMP inhibition was enhanced in potency (Table 5). Mutation of V169^5.30^ to glutamate, the amino acid present in other AR subtypes, enhanced both NECA and IB-MECA potency.

**Table 5.** Antagonistic potency of K18 at A_3_R mutants. cAMP accumulation as measured in Flp-In-CHO cells stably expressing WT or mutant A_3_R following stimulation with 10 μM forskolin, varying concentrations of IB-MECA and +/-K18 at the indicated concentration.

| **+ DMSO** |  |  |  |  |  |  |
| --- | --- | --- | --- | --- | --- | --- |
|  | **pIC<sub>50</sub><sup>a</sup>** | **E<sub>min</sub><sup>b</sup>** | **Basal<sup>c</sup>** | **True Basal<sup>d</sup>** | **Span<sup>e</sup>** | **n** |
| **WT** | 10.73 ± 0.1 | 35.0 ± 1.6 | 60.1 ± 0.9 | 57.7 ± 1.3 | 25.1 ± 2.0 | 11 |
| **L90A** | 9.03 ± 0.1**** | 43.3 ± 3.2 | 73.2 ± 2.7*** | 71.5 ± 2.9*** | 29.9 ± 1.8 | 8 |
| **V169A** | 11.33 ± 0.1**** | 30.9 ± 1.7 | 55.3 ± 2.4 | 54.1 ± 2.5 | 24.3 ± 2.1 | 10 |
| **M177A** | 7.65 ± 0.1**** | 38.6 ± 2.7 | 70.2 ± 2.0* | 66.7 ± 1.8 | 31.6 ± 2.0 | 7 |
| **I249A** | 10.76 ± 0.1 | 34.9 ± 2.2 | 62.6 ± 2.7 | 59.9 ± 2.6 | 27.7 ± 1.3 | 11 |
| **L264A** | 10.53 ± 0.1 | 41.1 ± 2.2 | 72.0 ± 2.3** | 70.8 ± 2.6** | 30.9 ± 2.2 | 9 |
| **+ 0.1 μM K18** |  |  |  |  |  |  |
|  | **pIC<sub>50</sub><sup>a</sup>** | **E<sub>min</sub><sup>b</sup>** | **Basal<sup>c</sup>** | **True Basal<sup>d</sup>** | **Span<sup>e</sup>** | **n** |
| **WT** | 10.64 ± 0.1 | 37.3 ± 1.8 | 63.0 ± 2.2 | 61.8 ± 2.6 | 25.8 ± 0.9 | 5 |
| **L90A** | 7.88 ± 0.1**** | 50.2 ± 3.4* | 77.2 ± 2.6** | 74.9 ± 2.8* | 27.0 ± 3.1 | 7 |
| **V169A** | 11.11 ± 0.1 * | 31.9 ± 1.8 | 62.6 ± 2.2 | 60.6 ± 3.1 | 30.6 ± 2.1 | 7 |
| **M177A** | 7.69 ± 0.1**** | 38.8 ± 2.5 | 70.9 ± 2.4 | 68.7 ± 2.3 | 32.1 ± 1.9 | 5 |
| **I249A** | 10.65 ± 0.1 | 35.6 ± 3.1 | 68.5 ± 3.3 | 67.0 ± 3.4 | 32.9 ± 1.3 | 8 |
| **L264A** | 9.86 ± 0.1*** | 45.7 ± 2.0 | 79.7 ± 2.7** | 77.7 ± 3.0** | 34.0 ± 2.8 | 7 |
| **+ 1 μM K18** |  |  |  |  |  |  |
|  | **pIC<sub>50</sub><sup>a</sup>** | **E<sub>min</sub><sup>b</sup>** | **Basal<sup>c</sup>** | **True Basal<sup>d</sup>** | **Span<sup>e</sup>** | **n** |
| **WT** | 9.65 ± 0.1 | 38.3 ± 2.4 | 67.4 ± 1.5 | 63.0 ± 1.8 | 29.1 ± 2.0 | 6 |
| **L90A** | 6.61 ± 0.1**** | 54.3 ± 3.6** | 76.7 ± 3.2 | 73.5 ± 3.1 | 22.4 ± 2.6 | 8 |
| **V169A** | 10.40 ± 0.1**** | 31.9 ± 2.3 | 68.8 ± 1.7 | 66.5 ± 2.0 | 36.9 ± 2.5 | 7 |
| **M177A** | 7.27 ± 0.1**** | 40.0 ± 3.5 | 71.4 ± 2.4 | 66.3 ± 2.5 | 31.4 ± 2.0 | 5 |
| **I249A** | 9.78 ± 0.1 | 36.9 ± 3.2 | 76.3 ± 3.7 | 73.1 ± 3.8 | 39.3 ± 2.1* | 8 |
| **L264A** | 8.80 ± 0.1**** | 47.9 ± 2.7 | 83.6 ± 2.1*** | 79.8 ± 2.4** | 35.7 ± 3.0 | 8 |
| **+ 10 μM K18** |  |  |  |  |  |  |
|  | **pIC<sub>50</sub><sup>a</sup>** | **E<sub>min</sub><sup>b</sup>** | **Basal<sup>c</sup>** | **True Basal<sup>d</sup>** | **Span<sup>e</sup>** | **n** |
| **WT** | 8.38 ± 0.2 | 45.1 ± 1.7 | 72.0 ± 1.5 | 68.9 ± 1.6 | 26.9 ± 1.3 | 7 |
| **L90A** | ND | 59.9 ± 2.9** | 81.7 ± 2.3 | 78.1 ± 2.4 | 22.8 ± 2.0 | 5 |
| **V169A** | 9.44 ± 0.1 **** | 33.5 ± 1.8** | 71.8 ± 1.6 | 69.2 ± 1.5 | 38.3 ± 2.0* | 8 |
| **M177A** | 6.12 ± 0.2**** | 45.7 ± 3.2 | 72.1 ± 2.3 | 67.6 ± 2.4 | 26.6 ± 1.6 | 5 |
| **I249A** | 8.55 ± 0.2 | 36.6 ± 2.1* | 78.0 ± 4.2 | 74.7 ± 4.2 | 38.7 ± 3.6* | 8 |
| **L264A** | 7.98 ± 0.1 | 49.1 ± 3.1 | 85.4 ± 2.7* | 82.8 ± 2.6** | 33.7 ± 5.0 | 5 |

<sup>a</sup>Negative logarithm of IB-MECA concentration required to produce a half-maximal response
<sup>b</sup>Minimum cAMP accumulation of IB-MECA as %100 μM forskolin. The lower plateau of the fitted sigmoidal dose response curve
<sup>c</sup>The upper plateau of the fitted sigmoidal dose response curve corresponding %100 μM forskolin
<sup>d</sup>The cAMP accumulation when stimulated with 10 μM forskolin only + DMSO/K18 at the indicated concentration
<sup>e</sup>The difference between E<sub>min</sub> and basal signalling
ND indicates an incomplete dose response curve due to the increased potency of K18 at this mutant.
Data are expressed as mean ± SEM obtained in *n* separate experiments. All individual experiments were conducted in duplicate. Statistical significance (\*, $p < 0.05$ ; \*\*, $p < 0.01$ ; \*\*\*, $p < 0.001$ ; \*\*\*\*, $p < 0.0001$ ) compared to WT IB-MECA stimulation +/- K18 at each indicated concentration was determined by one-way ANOVA with Dunnett's post-test.

Further, the affinity (pA_2_) of K18 at the WT and mutant A_3_R were compared in order to determine the potential antagonist binding site (Fig. 5, Table 5). K18 displayed no difference in affinity at I249A^6.54^ (compared to WT), whereas M177A^5.40^ and V169A^5.30^ were significantly smaller. Interestingly, we found an increase in the affinity for L90A^3.32^ and L264A^7.34^ when compared to WT. As would be expected, the K18 affinity at the A_3_R mutants was not different between agonists, confirming agonist independence (Supplementary Fig. 11). These experimental findings are reflected in our predicted binding pose of K18 at the WT A_3_R (Fig. 4).

**Figure 5.**
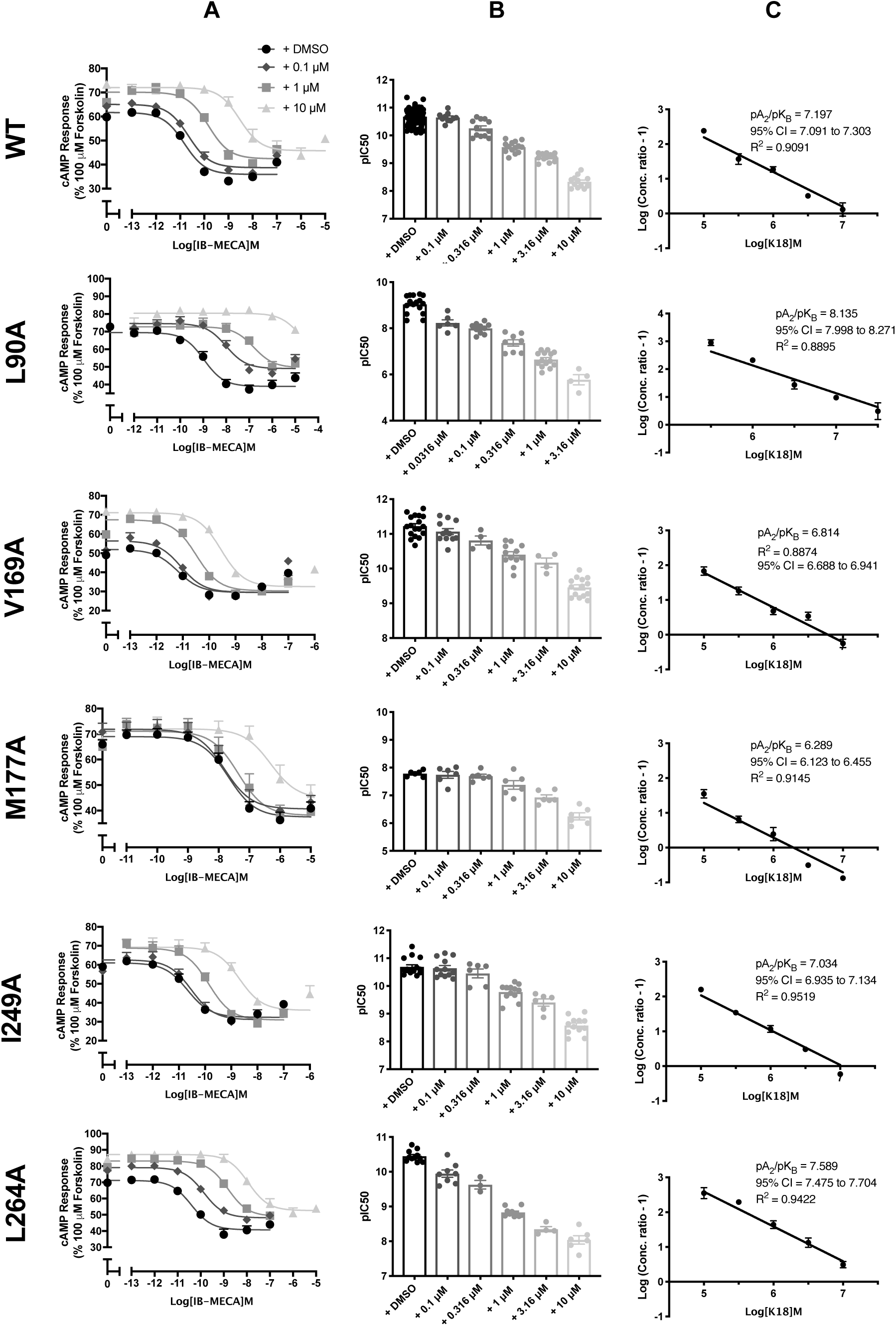
IB-MECA stimulated cAMP inhibition at WT or mutant A_3_R with increasing concentrations of K18. Flp-In-CHO cells (2000 cells/well) stably expressing WT or mutant A_3_R were exposed to forskolin 10 μM, IB-MECA and K18 at varying concentrations for 30 min and cAMP accumulation detected. **A**) Representative dose response curves are shown as mean ± SEM expressed as percentage maximum forskolin response (100 μM). **B**) pIC_50_ values for individual repeats including half-log concentration are shown as mean ± SEM **C**) Schild analysis of data represented in **A/B**.

### Kinetics of A_3_R antagonists determined through BRET

NanoBRET techniques have been successfully used to determine the real-time kinetics of ligand binding to GPCRs (Stoddart *et al.*, 2018, Sykes *et al.*, 2019, reviewed in Soave *et al.*, 2019). Here, we investigated the ability of the selective A_3_R antagonists MRS 1220, K17 or K18 to inhibit specific binding of the fluorescent A_3_R antagonist CA200645 to Nluc-A_3_R (Stoddart *et al.*, 2015, Bouzo-Lorenzo *et al.*, 2019). The kinetic parameters for CA200645 at Nluc-A_3_R were initially determined as K_on_ (*k*_1_) = 2.86 ± 0.89 × 10^7^ M^-1^, K_off_ (*k*_2_) = 0.4397 ± 0.014 min^-1^ with a K_D_ of 17.92 ± 4.45 nM. (Supplementary Fig 12). Our MRS 1220 kinetic data was fitted with the original ‘kinetic of competitive binding’ model (Motulsky and Mahan, 1984; built into GraphPad Prism 8.0) with a determined K_on_ (*k*_3_) and K_off_ (*k*_4_) rate of 3.25 ± 0.28 × 10^8^ M^-1^ min^-1^ and 0.0248 ± 0.005 min^-1^, respectively. This gave a residence time (RT) (RT = 1/K_off_) of 40.32 min. It was noticed in the analysis for K5, K17 and K18 that the fit in some cases was ambiguous (Regression with Prism 8: “Ambiguous”, 2019) and/or the fitted value of the compound dissociation rate constant was high (*k*_4_ > 1 min^-1^, corresponding to a dissociation *t*_1/2_ of < 42 sec). In order to determine the reliability of the fitted *k*_4_ value, data were also analysed using an equation that assumes compound dissociation is too rapid for the dissociation rate constant to be determined reliably and the fits to the two equations were compared (“Kinetics of competitive binding, rapid competitor dissociation”, derived in the Appendix I). This model allowed estimate of the equilibrium binding affinity of the compound (*K*_i_) but not the binding kinetics of K5, K17 and K18 (Supplementary Fig. 13 and Table 4). These affinity values were in agreement with those obtained via Schild analysis (except for K5) and were approximately 10-fold higher than those determined through radioligand binding (Table 4). Notably, the order of affinity (K5 < K17 < K18) was consistent.

### Assessing the species selectivity of A_3_R antagonists

There is a lack of selective antagonists for the rat A_3_R with only a small number reported including the 6-phenyl-1,4-dihydropyridine MRS 1191 and the trazoloquinazoline MRS 1220 which both bind the human A_3_R (Jacobson *et al.*, 1997). Comparison of residues of the binding area between human and rat A_3_R show that they differ in residues 167^5.28^, 169^5.30^, 176^5.37^, 253^6.58^, 264^7.34^ (Supplementary Fig. 14). The scarcity of rat A_3_R antagonists prompted us to investigate if our characterised compounds were also potential antagonists at the rat A_3_R. Using a Nluc-tagged rat A_3_R expressing HEK 293 cell line, we conducted NanoBRET ligand-binding experiments whereby we determined the ability of our compounds to inhibit specific binding of the fluorescent antagonist AV039 to Nluc-rat A_3_R. As expected, AV039 was displaced by increasing concentrations of MRS 1220 (pK_i_ 6.788 ±0.1) (Supplementary Fig. 15 and Supplementary Table 7). We found very weak binding of K17, K18, K10 and K32, with no binding detected below the concentration of 10 μM, whereas K1, K20, K23 and K25 were determined as potential rat A_3_R antagonists (pK_i_ 6.07 ±0.04, 5.71 ±0.03, 5.93 ±0.04 and 6.37 ±0.06, respectively) (Supplementary Fig. 15 and Supplementary Table 7). K25 had a higher binding affinity for the rat A_3_R when compared to the human A_3_R (Table 1) (pKi 6.37 ±0.1 and 5.81, respectively).

MD simulations of the rat A_3_R (performed as described previously for the human A_3_R) suggested that the presence of M264^7.34^ most likely hampers K18 binding due to steric hindrance of the dichloro-phenyl group (Supplementary Fig. 16). In contrast, MD simulations of K25 against rat A_3_R showed the formation of stable complex (Supplementary Fig. 17). Here, the 2-amido group of the thiophene ring is hydrogen-bonded to the amido group of N250^6.55^. The thiophene ring forms aromatic π-π stacking interaction with F168^5.29^ and the 5-(p-chlorophenyl) is oriented deep in the binding area making contacts with L90^3.32^, L246^6.51^ and W243^6.48^. M264^7.34^ forces the large lipophilic moiety of the thiophene ring (3-NHCOCH_2_SPh(CF_3_)_2_) of K25 to locate close to TM2 favouring contacts with A69^2.61^, V72^2.64^, and I268^7.39^ (Supplementary Fig. 17).

### Pharmacokinetic assessments of K18

The metabolic *in vitro t*_*1/2*_ (human liver microsomes, 0.1 mg/mL) of K18 (0.1 μM) was determined (0 to 60 minutes) as 24 minutes and the intrinsic clearance (CL_int_) calculated as 287.2 μl/min/mg (Supplementary Figure 18). This was comparable to the reference compound verapamil and terfenadine (0.1 μM) with *t*_1/2_ determined as 35 and 12 minutes and CL_int_ as 200.1 or 581.1 μl/min/mg, respectively. Human plasma stability assessment determined the percentage of K18 (1 μM) remaining after 120 minutes as 90%, with a *t*_1/2_ of >120 minutes. This is considerably higher than the reference compound propantheline (1 μM) which was determined to have a half-life of 55 minutes. The *t*_1/2_ of K18 (1 μM) in PBS (pH 7.4) over 240 minutes was determined as >240 minutes, with 87% remaining at 240 minutes and was comparable to the reference compound propantheline (1 μM), with a determined *t*_1/2_ of >240 minutes.

## DISCUSSION

*In silico* SBDD efforts in ligand discovery, have proven to be highly successful (Meng *et al.*, 2011). However, given the broad and similar orthosteric binding site of ARs, the search for an AR subtype specific compound often leads to compounds active at more than one of the AR subtypes (Kolb *et al.*, 2012). Given that AR subtypes play distinct roles throughout the body, obtaining highly specific receptor antagonists and agonists is crucial. Here, we presented the pharmacological characterisation of eight A_3_R antagonists identified though virtual screening. Of these eight compounds, K10, K17, K18, K20, K23, K25 and K32 were determined to be competitive. Whereas K20 and K23 were antagonists at both the A_1_R and A_3_R, K10, K17, K18, K25 and K32 were A_3_R selective antagonists. Indeed, we found no functional activity, or indeed binding affinity (< 30 μM), at the other AR subtypes.

K1, K20 and K23 showed weak antagonism of the A_2A_R (Figure 1, Table 2). We also tested the compounds at the A_2B_R and determined none of the compound showed A_2B_R antagonistic potency (Figure 1, Table 2). These selectivity findings were in agreement with our radioligand binding data (presented here (Supplementary Table 1) and in Lagarias *et al.*, 2018 for K1-25, K28 and K35). However, a number of compounds previously determined to have micromolar binding affinity for A_3_R (K5, K9, K21, K22, K24, K26, K27 and K34), showed no antagonistic potency in our initial functional screen. Further testing confirmed that these compounds were low potency antagonists and, although supporting the previously published radioligand binding data, confirmed the need for functional testing: not all compounds with binding affinity showed high functional potency.

We showed the A_3_R, when expressed in Flp-In™-CHO cells, displays constitutive activity. Compounds which preferably bind to the inactive (R) state, decreasing the level of constitutive activity (Giraldo *et al.*, 2007) and in the case of a G_i/o_ -coupled GPCR leading to an elevated cAMP, are referred to as inverse agonists. All eight characterised A_3_R antagonists and both characterised A_1_R antagonists (K20 and K23) were found to act as inverse agonists. We also reported an elevation in cAMP accumulation when cells were stimulated with DMSO, which was concentration-dependent. Given that even low concentrations of DMSO has been reported to interfere with important cellular processes (Tunçer *et al.*, 2018), the interpretation of these data should be made with caution. The initial virtual screening described in Lagarias *et al.*, 2018 was carried out using a combination of a ligand-based and structure-based strategy and the experimental structure of A_2A_R in complex with the antagonist ZM241385 (PDB ID 3EML) (Jaakola *et al.*, 2008), described as A_2A_R selective antagonist and inverse agonist (Lebon *et al.*, 2011). Our high hit rate for A_3_R selective antagonist appears counter-intuitive since the ligand-based virtual screening tool Rapid Overlay of Chemical Structures (ROCS) was used to predict structures similar to ZM241385 (Lagarias *et al.*, 2018). Indeed, ZM241385 has little affinity for A_3_R and 500-to 1000-fold selectivity for A_2A_R over A_1_R. However, as has been previously reported, the search for an AR subtype specific compound often leads to compounds active at multiple AR subtypes (Kolb *et al.*, 2012), likely due to their similar binding site.

We hypothesized that the presence of a chloro substituent in the phenyl ring of 3-phenyl-isoxazole favoured A_3_R affinity and activity, as following 0Cl < 1Cl < 2Cl i.e. K5 < K17 < K18. This theory is supported by both our radioligand binding, NanoBRET ligand-binding and functional data which determine the relative potency and affinity of the three related compounds K5, K17 and K18 as K5 < K17 < K18. The MD simulations showed that these compounds adopted a similar binding mode at the A_3_R orthosteric binding site, but the free-energy MM-PBSA calculations showed that K18, having two chlorine atoms and an increased lipophilicity, leaves the solution state more efficiently and enters the highly lipophilic binding area.

For the first time, we have demonstrated the utilisation of a new model which expands on the ‘Kinetic of competitive binding’ model (Motulsky and Mahan, 1984; built into Prism) for fitting fast kinetics data obtained from NanoBRET experiments and assumes the unlabelled ligand rapidly equilibrates with the free receptor. Very rapid competitor dissociation can lead to failure of the fit, eliciting either an ambiguous fit (Regression with Prism 8: “Ambiguous”, 2019) or unrealistically large K_3_ and K_4_ values. Whereas we were able to successfully fit the MRS 1220 kinetic data with the Motulsky and Mahan model due to its slow dissociation, fitting of K5, K17 and K18 kinetic data with this model often resulted in an ambiguous fit. Our new model, assuming fast compound dissociation, successfully fit the data and allowed the determination of binding affinity. In the cases where the data was able to fit the Motulsky and Mahan model, the dissociation constant was higher (of the order of 1 min-1), indicating rapid dissociation. Although we found nearly a 10-fold differences in determined binding affinity for MRS 1220, K5, K17 and K18 between BRET ligand binding and radioligand binding assays, we demonstrated the order of affinity remains consistent. Indeed, this was seen across all three experimental approached: Schild analysis, NanoBRET ligand-binding assay and radioligand binding.

Our MD simulations describe the potential binding site of K18, our most potent and selective A_3_R antagonist, within the A_3_R orthosteric binding area (Fig. 4A). Here, K18 is stabilised through hydrogen bonding interactions between the amino group and thiazole ring of the ligand and the amide side chain of N250^6.55^. In addition, the dichloro-phenyl ring can be oriented to the unique lipophilic area of A_3_R including V169^5.30^, M177^5.40^, I249^6.54^ and L264^7.34^ stabilized in that cleft through attractive vdW interactions; K18 is further stabilized through *π-π* aromatic stacking interactions between isoxazole ring and the phenyl group of F168^5.29^ and the thiazole group is oriented deeper into the receptor favouring interactions with L246^6.51^ and L90^3.32^ and possibly with I268^7.39^. The determination of critical residues for antagonist binding becomes particularly difficult in the case of competitive antagonists whereby important amino acids are likely overlapping with those for agonist binding. In combination with our mutagenesis data, the final binding pose of K18 appears to be within the orthosteric binding site, involving residues previously described to be involved in binding of A_3_R compounds (Arruda *et al.*, 2017). We reported no detectable G_i/o_ response following co-stimulation with forskolin and NECA or IB-MECA for A_3_R mutants F168A^5.29^, L246A^6.51^, N250A^6.55^ and I268A^7.39^ (Stamatis *et al.*, 2019). These findings are in line with previous mutagenesis studies investigating residues important for agonist and antagonist binding at the human A_3_R (Gao *et al.*, 2002, May *et al.*, 2012). L90A^3.32^, V169A^5.30^, M177A^5.40^, I249A^6.54^ and L264A^7.34^ A_3_R all showed a detectable G_i/o_ response when stimulated with agonists (Stamatis *et al.*, 2019).

Through performing Schild analysis (results of which were used to inform modelling in Lagarias *et al.*, 2019) we were able to experimentally determine the effect of receptor mutation on antagonist affinity for L90A^3.32^, V169A/E^5.30^, M177A^5.40^, I249A^6.54^ and L264A^7.34^ A_3_R. The pA_2_ value for I249A^6.54^ A_3_R was similar to WT, whereas M177A^5.40^ and V169A^5.30^ were significantly smaller suggesting these residues appear to be involved in K18 binding. Interestingly we found an increase in the pA_2_ for L90A^3.32^ and L264A^7.34^ when compared to WT, suggesting an enhanced ability of K18 to act as an antagonist. Further evidence was provided by the MM-PBSA calculations which were in agreement, based on the proposed binding model, between the calculated binding free energy by congeners of K18 having one or no chlorine atoms, i.e. compounds K17 and K5, and the known A_3_R antagonist MRS 1220, and binding affinities and antagonistic potencies. Importantly, substitution of the 1,3-thiazole ring in K17 with either a 2-pyridinyl ring (K32) or a 3-pyridinyl ring (K10) but not a 4-pyridinyl ring (K11) maintained A_3_R antagonistic potency. Although we have not directly determined the effects of similar pyridinyl ring substitutions on the higher affinity antagonist K18, we suspect there would be no significant increase in the potency of K18 given the small changes we observed for K17.

The human and rat A_3_R display 72% homology (Salvatore *et al.*, 1993). Antagonists that are A_3_R-selective across species or at rat A_3_R alone are useful pharmacological tools to define the role of these receptors. The lack of rat A_3_R selective antagonists prompted us to investigate if our characterised A_3_R antagonists were potential antagonists at the rat A_3_R. We reported no binding of our lead A_3_R antagonist, K18, at the rat A_3_R and MD simulations suggest this is due to steric hinderance by M264^7.34^. This finding suggests that K18 may not only be A_3_R specific within the human ARs but may also be selective across species. Of the compounds that showed rat A_3_R binding (K1, K20, K23 and K25), K25 showed the highest binding affinity and represents an interesting candidate for further investigation. MD simulations showed K25 forms a stable complex with rat A_3_R and we reported a potential binding pose.

In conclusion, through pharmacological characterisation of a number of potential A_3_R antagonists, this study has determined K18 as a specific (<1 µM) A_3_R competitive antagonist. Our mutagenic studies, supported by MD simulations, identified the residues important for K18 binding are located within the orthosteric site of the A_3_R. Importantly, the absence of a chloro substituent, as is the case in K5, led to affinity loss. Further MD simulations have investigated the selectivity profile of K18 and have demonstrated that K18 failed to bind A_1_R and A_2A_R due to a more polar area close to TM5, TM6 when compared to A_3_R (Lagarias *et al.*, 2019). While a number of novel potent and selective A_3_R antagonists have been previously described (Yaziji *et al*., 2011, Yaziji *et al.*, 2013, Areias *et al.*, 2019) we present findings of a unique scaffold which can be used as a starting point for detailed structure-activity relationships (SARs) and represents a useful tool that warrants further assessment for its therapeutic potential.

## MATERIALS AND METHODS

### Cell culture and Transfection

Cell lines were maintained using standard subculturing routines as guided by the European Collection of Cell Culture (ECACC) and checked annually for mycoplasma infection using an EZ-PCR mycoplasma test kit from Biological Industries (Kibbutz Beit-Haemek, Israel). All procedures were performed in a sterile tissue culture hood using aseptic technique and solutions used in the propagation of each cell line were sterile and pre-warmed to 37°C. All cells were maintained at 37°C with 5% CO_2_, in a humidified atmosphere. This study used CHO cell lines as a model due to the lack of endogenous AR subtype expression (Brown *et al.*, 2008). CHO-K1-A_1_R, CHO-K1-A_2A_R, CHO-K1-A_2B_R and CHO-K1-A_3_R cells were routinely cultured in Hams F-12 nutrient mix (21765029, Thermo Fisher Scientific) supplemented with 10% Foetal bovine serum (FBS) (F9665, Sigma-Aldrich). Flp-In-CHO cells purchased from Thermo Fisher Scientific (R75807) were maintained in Hams F-12 nutrient mix supplemented with 10% FBS containing 100 μg/mL Zeocin™ Selection Antibiotic (Thermo Fisher Scientific).

Stable Flp-In-CHO cell lines were generated through co-transfection of the pcDNA5/FRT expression vector (Thermo Fisher Scientific) containing the gene of interest and the Flp recombinase expressing plasmid, pOG44 (Thermo Fisher Scientific). Transfection of cells seeded in a T25 flask at a confluency of ≥80% was performed using TransIT®-CHO Transfection Kit (MIR 2174, Mirus Bio), in accordance with the manufacturer’s instructions. Here, a total of 6 μg of DNA (receptor to pOG44 ratio of 1:9) was transfected per flask at a DNA:Mirus reagent ratio of 1:3 (w/v). 48 hours post-transfection, selection using 600 μg/mL hygromycin B (Thermo Fisher Scientific) (concentration determined through preforming a kill curve) was performed for two days prior to transferring the cells into a fresh T25 flask. Stable Flp-In-CHO cell lines expressing the receptor of interest were selected using 600 μg/mL hygromycin B whereby the media was changed every two days. Successful mutant cell line generation for non-signalling mutants were confirmed by Zeocin™ sensitivity (100 μg/mL).

The Nluc-tagged human A_3_R expressing HEK 293 cell line along with the Nluc-tagged rat A_3_R pcDNA3.1+ construct for the generation of stable Nluc-tagged rat A_3_R expressing HEK 293 cells were kindly gifted to us by Stephen Hill and Stephen Briddon (University of Nottingham). HEK 293 cells in a single well of 6-well plate (confluency ≥80%) were transfected with 2 μg of DNA using polyethyleneimine (PEI, 1 mg/ml, MW = 25,000 g/mol) (Polysciences Inc) at a DNA:PEI ratio of 1:6 (w/v). Briefly, DNA and PEI were added to seporate sterile tubes contianing 150 mM sodium chloride (NaCl) (total volume 50 μl), allowed to incubate at room termperature for 5 minutes, mixing together and incubating for a further 10 minutes prior to adding the combined mix dropwise to the cells. 48 hours post-transfection, stable Nluc-rat A_3_R expressing HEK 293 cell were selected using 600 μg/mL Geneticin (Thermo Fisher Scientific) whereby the media was changed every two days. HEK 293 cell lines were routinely cultured in DMEM/F-12 GlutaMAX™ (Thermo Fisher Scientific) supplemented with 10% FBS (F9665, Sigma-Aldrich).

### Constructs

The human A_3_R originally in pcDNA3.1+ (ADRA3000000, cdna.org) was cloned into the pcDNA5/FRT expression vector and co-transfected with pOG44 to generate a stable Flp-In-CHO cell line. Mutations within the A_3_R were made using the QuikChange Lightening Site-Directed Mutagenesis Kit (Agilent Technologies) in accordance with the manufacturer’s instructions. The Nluc-tagged rat A_3_R pcDNA3.1+ construct, used in the generation of the stable Nluc-tagged rat A_3_R expressing HEK 293 cell line was kindly gifted to us by Stephen Hill and Stephen Briddon (University of Nottingham). All oligonucleotides used for mutagenesis were designed using the online Agilent Genomics ‘QuikChange Primer Design’ tool and are detailed in Stamatis *et al.*, 2019 (Table S4) and purchased from Merck. All constructs were confirmed by in-house Sanger sequencing.

### Compounds

Adenosine, NECA ((2S,3S,4R,5R)-5-(6-aminopurin-9-yl)-N-ethyl-3,4-dihydroxyoxolane-2-carboxamide), IB-MECA ((2S,3S,4R,5R)-3,4-dihydroxy-5-[6-[(3-iodophenyl)methylamino]purin-9-yl]-N-methyloxolane-2-carboxamide), HEMADO ((2R,3R,4S,5R)-2-(2-hex-1-ynyl-6-methylaminopurin-9-yl)-5-(hydroxymethyl)oxolane-3,4-diol), DPCPX (8-cyclopentyl-1,3-dipropyl-7H-purine-2,6-dione) and MRS 1220 (N-(9-chloro-2-furan-2-yl-[1,2,4]triazolo[1,5-c]quinazolin-5-yl)-2-phenylacetamide) were purchased from Sigma-Aldrich and dissolved in dimethyl-sulphoxide (DMSO). CA200645, a high affinity AR xanthine amine congener (XAC) derivative containing a polyamide linker connected to the BY630 fluorophore, was purchased from HelloBio (Bristol, UK) and dissolved in DMSO.

AV039, a highly potent and selective fluorescent antagonist of the human A_3_R based on the 1,2,4-Triazolo[4,3-a]quinoxalin-1-one linked to BY630 (Vernall *et al.*, 2012), was kindly gifted to us by Stephen Hill and Stephen Briddon (University of Nottingham). <u>PMA</u> was purchased from Sigma-Aldrich. Compounds under investigation were purchased from e-molecules and dissolved in DMSO. The concentration of DMSO was maintained to <1.5% across all experiments (1.26% for all cAMP assays, 1% for pERK1/2 assays and 1.02% or 1.1% for NanoBRET ligand-binding experiments using CA200645 or AV039, respectively).

### cAMP accumulation assay

For cAMP accumulation (A_2A_R and A_2B_R) or inhibition (A_1_R or A_3_R) experiments, cells were harvested and re-suspended in stimulation buffer (PBS containing 0.1% BSA and 25 μM rolipram) and seeded at a density of 2,000 cells per well of a white 384-well Optiplate and stimulated for 30 minutes with a range of agonist concentrations. In order to allow the A_1_R/A_3_R mediated G_i/o_ response to be determined, co-stimulation with forskolin, an activator of AC (Zhang *et al.*, 1997), at the indicated concentration (depending on cell line) was performed. For testing of potential antagonists, cells received a co-stimulation stimulated with forskolin, agonist and compound/DMSO control. cAMP levels were then determined using a LANCE® cAMP kit as described previously (Knight *et al*, 2016). In order to reduce evaporation of small volumes, the plate was sealed with a ThermalSeal® film (EXCEL Scientific) at all stages.

### Phospho-ERK assay

ERK1/2 phosphorylation was measured using the homogeneous time resolved fluorescence (HTRF)® Phospho-ERK (T202/Y204) Cellular Assay Kit (Cisbio Bioassays, Codolet, France) two-plate format in accordance with the manufacturer’s instructions. A_3_R expressing Flp-In-CHO were seeded at a density of 2,000 cells per well of a white 384-well Optiplate and stimulated with agonist and test compounds for 5 minutes at 37°C. Plate reading was conducted using a Mithras LB 940 (Berthold technology). All results were normalised to 5 minutes stimulation with 1 μM PMA, a direct protein kinase C (PKC) activator (Jiang and Fleet, 2012). To determine if the measured pERK1/2 level was G_i_-mediated, we treated cells with Pertussis toxin (PTX) (Tocris Biosciences) for 16 hours at 100 ng/mL prior to pERK assay.

### Radioligand Binding

All pharmacological methods followed the procedures as described in the literature (Klotz *et al.*, 1998). In brief, membranes for radioligand binding were prepared from CHO cells stably transfected with hAR subtypes in a two-step procedure. In the first step, cell fragments and nuclei were removed at 1000 x g and then the crude membrane fraction was sedimented from the supernatant at 100000 x g. The membrane pellet was resuspended in the buffer used for the respective binding experiments and it was frozen in liquid nitrogen and stored at −80°C. For radioligand binding at the A_1_R, 1 nM [^3^H]CCPA was used, for A_2A_R 10 nM [^3^H]NECA and for A_3_R 1 nM [^3^H]HEMADO. Non-specific binding of [^3^H]CCPA was determined in the presence of 1 mM theophylline and in the case of [^3^H]NECA and [^3^H]HEMADO 100 μM R-PIA was used. *K*_i_ values from competition experiments were calculated using Prism (GraphPad Software, La Jolla, CA, U.S.A.) assuming competitive interaction with a single binding site. The curve fitting results (see Fig. 8 in Lagarias *et al.*, 2018) showed R^2^ values ≥ 0.99 for all compounds and receptors, indicating that the used one-site competition model assuming a Hill slope of n=1 was appropriate.

### NanoBRET ligand-binding

Through the use of NanoBRET, real-time quantitative pharmacology of ligand-receptor interactions can be investigated in living cells. CA200645, acts as a fluorescent antagonist at both A_1_R and A_3_R with a slow off-rate (Stoddart *et al.*, 2012). Using an N-terminally NanoLuc (Nluc)-tagged A_3_R expressing cell line, competition binding assays were conducted. The kinetic data was fitted with the ‘kinetic of competitive binding’ model (Motulsky and Mahan, 1984; built into Prism) to determine affinity (pK_i_) values and the association rate constant (K_on_) and dissociation rates (K_off_) for unlabelled A_3_R antagonists. In several cases this model resulted in an ambiguous fit (Regression with Prism 8: “Ambiguous”, 2019). We developed a new model which expands on the ‘kinetic of competitive binding’ model to accommodate very rapid competitor dissociation, assuming the unlabelled ligand rapidly equilibrates with the free receptor. This method allows determination of compound affinity (pK_i_) from the kinetic data.

In order to identify if the characterised compounds also bound the rat A_3_R, we conducted competition binding assays using Nluc-tagged rat A_3_R expressing HEK 293 cells and the fluorescent compound AV039 (Vernall *et al.*, 2012) rather than xanthine based CA200645, which have previously been reported as inactive at rat A_3_R (Siddiqi *et al.*, 1996). For both human and rat A_3_R experiments, filtered light emission at 450 nm and > 610 nm (640-685 nm band pass filter) was measured using a Mithras LB 940 and the raw BRET ratio calculated by dividing the 610 nm emission with the 450 nm emission. Here, Nluc on the N-terminus of A_3_R acted as the BRET donor (luciferase oxidizing its substrate) and CA200645/AV039 acted as the fluorescent acceptor. CA200645 was used at 25 nM, as previously reported (Stoddart *et al.*, 2015) and AV039 was used at 100 nM (pre-determined K_D_, 102 ± 7.59 nM). BRET was measured following the addition of the Nluc substrate, furimazine (0.1 μM). Nonspecific binding was determined using a high concentration of unlabelled antagonist, MRS 1220 at 10 nM or 10 μM, for human and rat A_3_R, respectively.

### Receptor binding kinetics data analysis

Specific binding of tracer vs time data was analysed using the Motulsky and Mahan method (Motulsky and Mahan, 1984; built into Prism) to determine the test compound association rate constant and dissociation rate constant. Data were fit to the “Kinetics of competitive binding” equation in Prism 8.0 (GraphPad Software Inc, San Diego, CA):

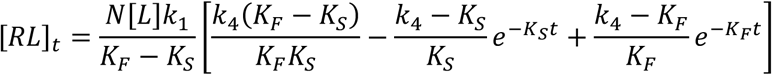

where,

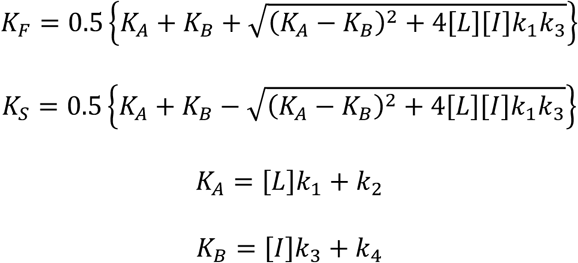

[*RL*]_*t*_ is specific binding at time *t, N* the B_max_, [*L*] the tracer concentration, [*I*] the unlabelled competitor compound concentration, *k*_1_ the tracer association rate constant, *k*_2_ the tracer dissociation rate constant, *k*_3_ the compound association rate constant and *k*_4_ the compound dissociation rate constant.

Data were also analysed using an equation that assumes compound dissociation is too rapid for the dissociation rate constant to be determined reliably and the fits to the two equations compared (“Kinetics of competitive binding, rapid competitor dissociation”, derived in the Appendix I, Supplementary material). This equation assumes rapid equilibration between compound and receptor and consequently provides an estimate of the equilibrium binding affinity of the compound (*K*_i_) but not the binding kinetics of the compound. The equation is,

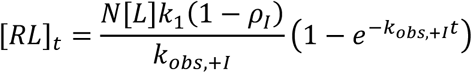

where *ρ*_*I*_ is fractional occupancy of receptors not bound by *L*:

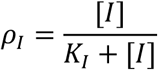

and *k*_*obs*,+ *I*_ is the observed association rate of tracer in the presence of competitor, defined as,

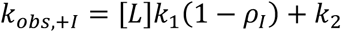

The fits to the two equations were compared statistically using a partial F-test in Prism 8 (Motulsky, 2019).

### Pharmacokinetic assessments of K18

Preliminary pharmacokinetic assessments of K18 was out-sourced to Eurofins Panlabs (Missouri, U.S.A) and including tests for intrinsic clearance (human liver microsomes), plasma (human) stability and half-life in PBS. These tests were conducted in duplicate using a single concentration of K18 (0.1 μM or 1 μM) using the substrate depletion method. Here, the percentage of K18 remaining at various incubation times was detected using high-performance liquid chromatography mass spectrometry (HPLC-MS). Reference compounds (<u>verapamil</u>, <u>terfenadine</u> and <u>propantheline</u>) were supplied and tested alongside K18. The half-life (*t*_1/2_) was estimated from the slope (k) of percentage compound remaining (In(%K18 remaining)) versus time (*t*_*1/2*_ = - In(2)/k), assuming first order kinetics. The intrinsic clearance (CL_int,_ in μl/min/mg) was calculated according to the following formula:

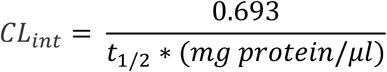

### Data and Statistical analysis

All *in vitro* assay data was analysed using Prism 8.0 (GraphPad software, San Diego, CA), with all dose-inhibition or response curves being fitted using a 3-parameter logistic equation to calculate response range or E_max_ and IC/EC_50_. Experimental design ensured random distribution of treatments across 96/384-well plates to avoid systematic bias. Agonist stimulation alone was used as an intrinsic control across all experiments. Although initial screening of the 50 compounds was blinded, due to limitations in resources, it was not possible to perform all of our experiments in a blinded manner. Normalisation was used to control for unwanted sources of variation between experimental repeats.

In the rare cases where significant outliers were identified through the ROUT method (performed in Prism with Q set to 2% (defines the chance of falsely identifying one or more outliers)) (Statistics with Prism 7: “How to: Identify outliers”, 2019), these were excluded from data analysis and presentation. Dose-inhibition/dose-response curves were normalised to forskolin, expressed as percentage forskolin inhibition for G_i_-coupled A_1_R and A_3_R (1 μM or 10 μM, respectively) or stimulation for A_2A_R and A_2B_R (100 μM, representing the maximum cAMP accumulation of the system), relative to NECA/IB-MECA (agonist allowing comparison across AR subtypes and a selective A_3_R agonist, respectively). For cAMP experiments on A_3_R mutants, data was normalised to 100 μM forskolin, representing the maximum cAMP accumulation possible for each cell line. In the case of pERK1/2 response, normalisation was performed to PMA, a direct PKC activator providing the maximum pERK1/2 level of the system.

Schild analysis was performed to obtain pA_2_ values (the negative logarithm to base 10 of the molar concentration of an antagonist that makes it necessary to double the concentration of the agonist to elicit the original submaximal response obtained by agonist alone (Schild, 1947)) for the potential antagonists. In cases where the Schild slope did not differ significantly from unity, the slope was constrained to unity giving an estimate of antagonist affinity (p*K*B). pA_2_ and p*K*B coincide when the slope is exactly unity. Through performing Schild analysis, whereby the pA_2_ is independent of agonist, we were able to experimentally determine the effect of receptor mutation on antagonist binding. The pA_2_ values obtained through conducting Schild analysis of K18 at WT and mutant A_3_R were compared in order to indicate important residues involved in K18 binding. Whereas an increase in the pA_2_ for a particular mutant when compared to WT suggested the antagonist was more potent, a decrease indicated a reduced potency.

All experiments were conducted in duplicate (technical replicates) to ensure the reliability of single values. The data and statistical analysis comply with the recommendations on experimental design and analysis in pharmacology (Curtis *et al.*, 2018). Statistical analysis, performed using Prism 8.0, was undertaken for experiments where the group size was at least n = 5 and these independent values used to calculate statistical significance (*\*, p< 0.05; **, p<0.01; ***, p<0.001; ****, p<0.0001*) using a one-way ANOVA with a Dunnett’s post-test for multiple comparisons or Students’ t-test, as appropriate. Any experiments conducted n < 5 should be considered preliminary. Compounds taken forward for further investigation after our preliminary screening (n = 3) were selected based on a high mean cAMP accumulation (>80%).

### Computational biochemistry

#### MD simulations

Preparation of the complexes between human A_3_R with K5, K17, K18 or MRS 1220 and rat A_3_R with K18 or K25 was based on a homology model of A_2A_R (see Appendix II in Supplementary material). Each ligand-protein complex was embedded in hydrated POPE bilayers. A simulation box of the protein-ligand complexes in POPE lipids, water and ions was built using the System Builder utility of Desmond (Desmond Molecular Dynamics System, version 3.0; D.E. Shaw Res. New York, 2011; Maest. Interoperability Tools, 3.1; Schrodinger Res. New York, 2012.). A buffered orthorhombic system in 10 Å distance from the solute atoms with periodic boundary conditions was constructed for all the complexes. The MD simulations were performed with Amber14 and each complex-bilayer system was processed by the LEaP module in AmberTools14 under the AMBER14 software package (Case *et al.*, 2014). Amber ff14SB force field parameters (Maier *et al*., 2015) were applied to the protein, lipid14 to the lipids (Dickson *et al.*, 2014), GAFF to the ligands (Wang *et al.*, 2004) and TIP3P (Jorgensen *et al.*, 1983) to the water molecules for the calculation of bonded, vdW parameters and electrostatic interactions. Atomic charges were computed according to the RESP procedure (Bayly *et al.*, 1993) using Gaussian03 (Frisch *et al.*, 2003) and antechamber of AmberTools14 (Case *et al.*, 2014). The temperature of 310 K was used in MD simulations in order to ensure that the membrane state is above the main phase transition temperature of 298 K for POPE bilayers (Koynova and Caffrey, 1998). In the production phase, the relaxed systems were simulated in the NPT ensemble conditions for 100 ns. The visualization of produced trajectories and structures was performed using the programs Chimera (Pettersen *et al.*, 2004) and VMD (Humphrey *et al.*, 1996). All the MD simulations were run on GTX 1060 GPUs in lab workstations or on the ARIS Supercomputer.

#### MM-PBSA calculations

Relative binding free energies of the complexes between K5, K17, K18, MRS 1220 and A_3_R was estimated by the 1-trajectory MM-PBSA approach (Massova and Kollman, 2000). Effective binding energies (Δ*G*_eff_) were computed considering the gas phase energy and solvation free energy contributions to binding. For this, structural ensembles were extracted in intervals of 50 ps from the last 50 ns of the production simulations for each complex. Prior to the calculations all water molecules, ions, and lipids were removed, and the structures were positioned such that the geometric centre of each complex was located at the coordinate origin. The polar part of the solvation free energy was determined by calculations using Poisson-Boltzmann (PB) calculations (Homeyer and Gohike, 2013). In these calculations, a dielectric constant of *ε*_solute_ = 1 was assigned to the binding area and *ε*_solute_ = 80 for water. Using an implicit solvent representation for the calculation of the effective binding energy is an approximation to reduce the computational cost of the calculations. The binding free energy for each complex was calculated using equation (1)

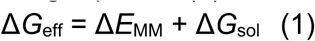

In equation (1) Δ*G*_eff_ is the binding free energy for each calculated complex neglecting the effect of entropic contributions or assuming to be similar for the complexes studied. Δ*E*_MM_ defines the interaction energy between the complex, the protein and the ligand as calculated by molecular mechanics in the gas phase. Δ*G*_sol_ is the desolvation free energy for transferring the ligand from water in the binding area calculated using the PBSA model. The terms for each complex Δ*E*_MM_ and Δ*G*_sol_ are calculated using equations (2) and (3)

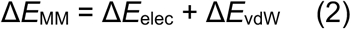

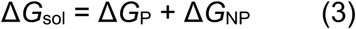

In equation (2) Δ*E*_elec_ and Δ*E*_vdW_ are the electrostatic and the vdW interaction energies, respectively. In equation (3) Δ*G*_P_ is the electrostatic or polar contribution to free energy of solvation and the term Δ*G*_NP_ is the non-polar or hydrophobic contribution to solvation free energy. Molecular mechanics energies and the non-polar contribution to the solvation free energy were calculated with the *mmpbsa.pl* module (Miller *et al.*, 2012) of Amber14 (Case *et*

### Nomenclature of Targets and Ligands

Key protein targets and ligands in this article are hyperlinked to corresponding entries in http://www.guidetopharmacology.org, the common portal for data from the IUPHAR/BPS Guide to PHARMACOLOGY (Harding *et al.*, 2018), and are permanently archived in the Concise Guide to PHARMACOLOGY 2017/18 (Alexander *et al.*, 2017).

## Abbreviations

Regression with Prism 8: ‘Ambiguous’. (2019).

https://www.graphpad.com/guides/prism/8/curve-fitting/reg_analysischeck_nonlin_ambiguous.htm

Statistics with Prism 7: “How to: Identify outliers”. (2019).

## Acknowledgments

We gratefully acknowledge the support of the Leverhulme Trust (RPG-2017-255) (KB and GL) and the BBSRC (BB/M00015X/2) (GL). This research represents part of the Ph.D work of P.L. We thank Chiesi Hellas which supported this research (SARG No 10354) and the State Scholarships Foundation (IKY) for providing a Ph.D fellowship to P.L. (MIS 5000432, NSRF 2014-2020). The work of E.V. is implemented through IKY scholarships programme and co-financed by the European Union (European Social Fund - ESF) and Greek national funds through the action entitled “Reinforcement of Postdoctoral Researchers”, in the framework of the Operational Program “Human Resources Development Program, Education and Lifelong Learning” of the National Strategic Reference Framework (NSRF) 2014 – 2020. This work was supported by computational time granted from the Greek Research & Technology Network (GRNET) in the National HPC facility - ARIS - under project IDs pr005010. We would like to thank Stephen Hill, Stephen Briddon and Mark Soave (University of Nottingham) for gifting the Nluc-A_3_R cell line, Nluc-rat A_3_R construct, the fluorescent antagonist AV039 and their technical advice. We are also grateful to Sonja Kachler for her technical assistance.

## SUPPORTING INFORMATION

**Supplementary Figure 1.**
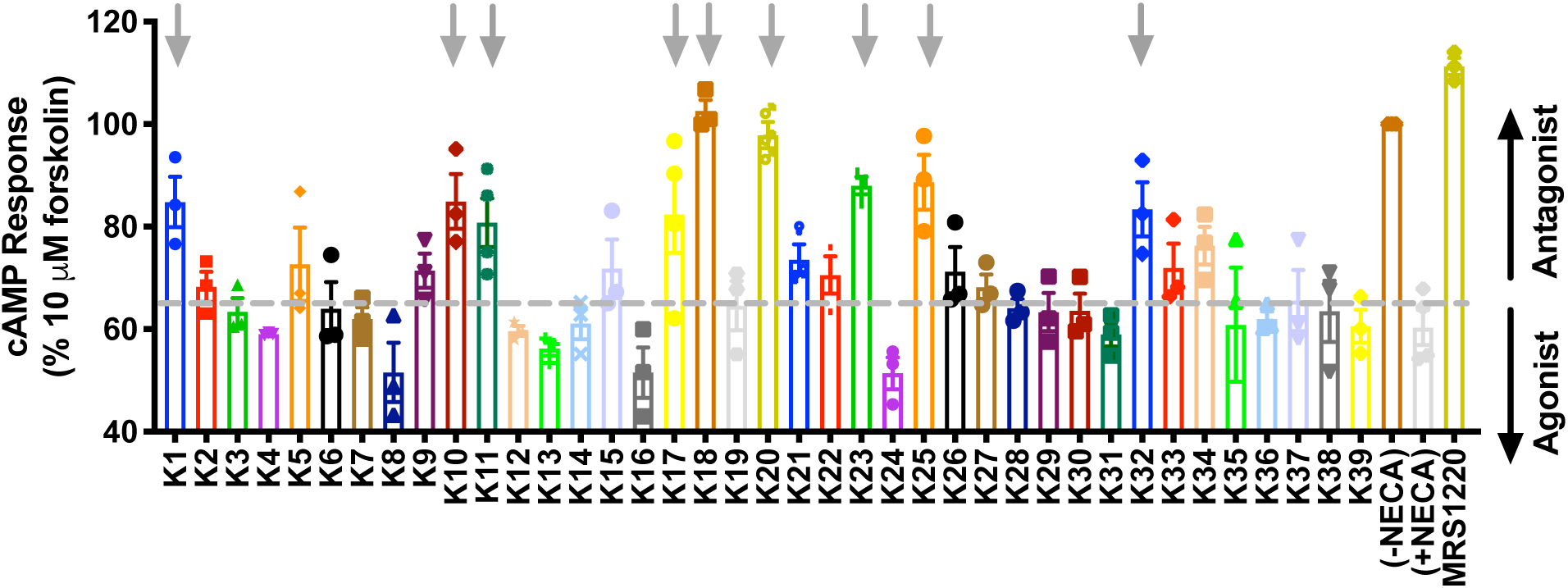
Screening for potential antagonists at the A_3_R. cAMP accumulation was determined in Flp-In CHO cells stably expressing A_3_R (2000 cells/well) co-stimulated for 30 minutes with 10 μM forskolin, NECA at the pre-determined IC_80_ concentration (3.16 nM) and 1 μM of compound/DMSO control. An elevation in cAMP accumulation above that of 10 μM forskolin and NECA, as indicated by the grey dotted line, suggesting the compound is acting as an antagonist (black upwards arrow). Included is MRS 1220 (1 μM) as a positive control for competitive antagonist of A_3_R. A reduction of cAMP accumulation (black downwards arrow) could indicate a compound is acting as an agonist. All values are mean ± SEM expressed as % 10 μM forskolin response (‘DMSO’) where *n* = 3 independent experimental repeats, conducted in duplicate. Grey downward arrow indicates potential antagonists with a cAMP level >80%.

**Supplementary Table 1.**
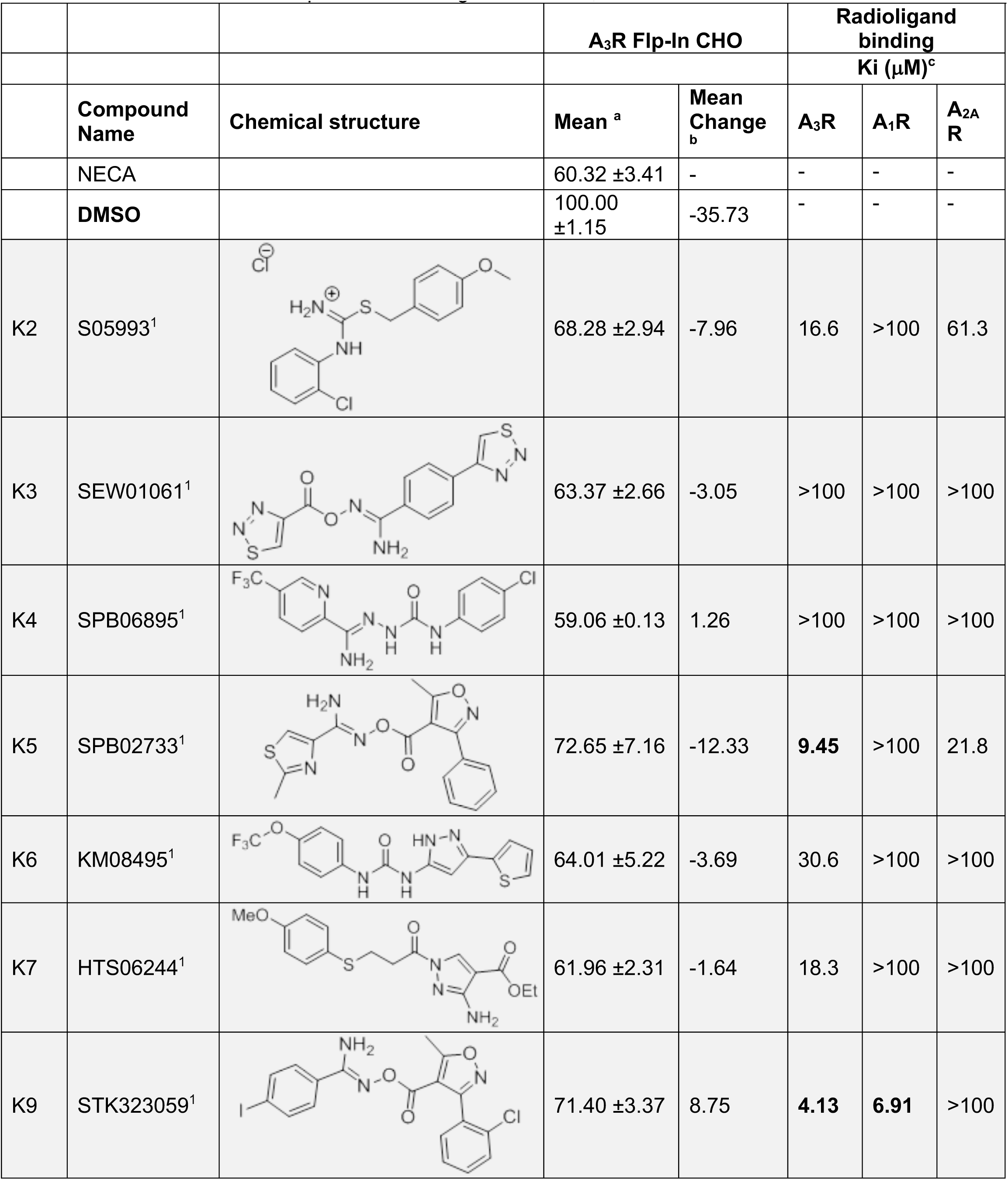
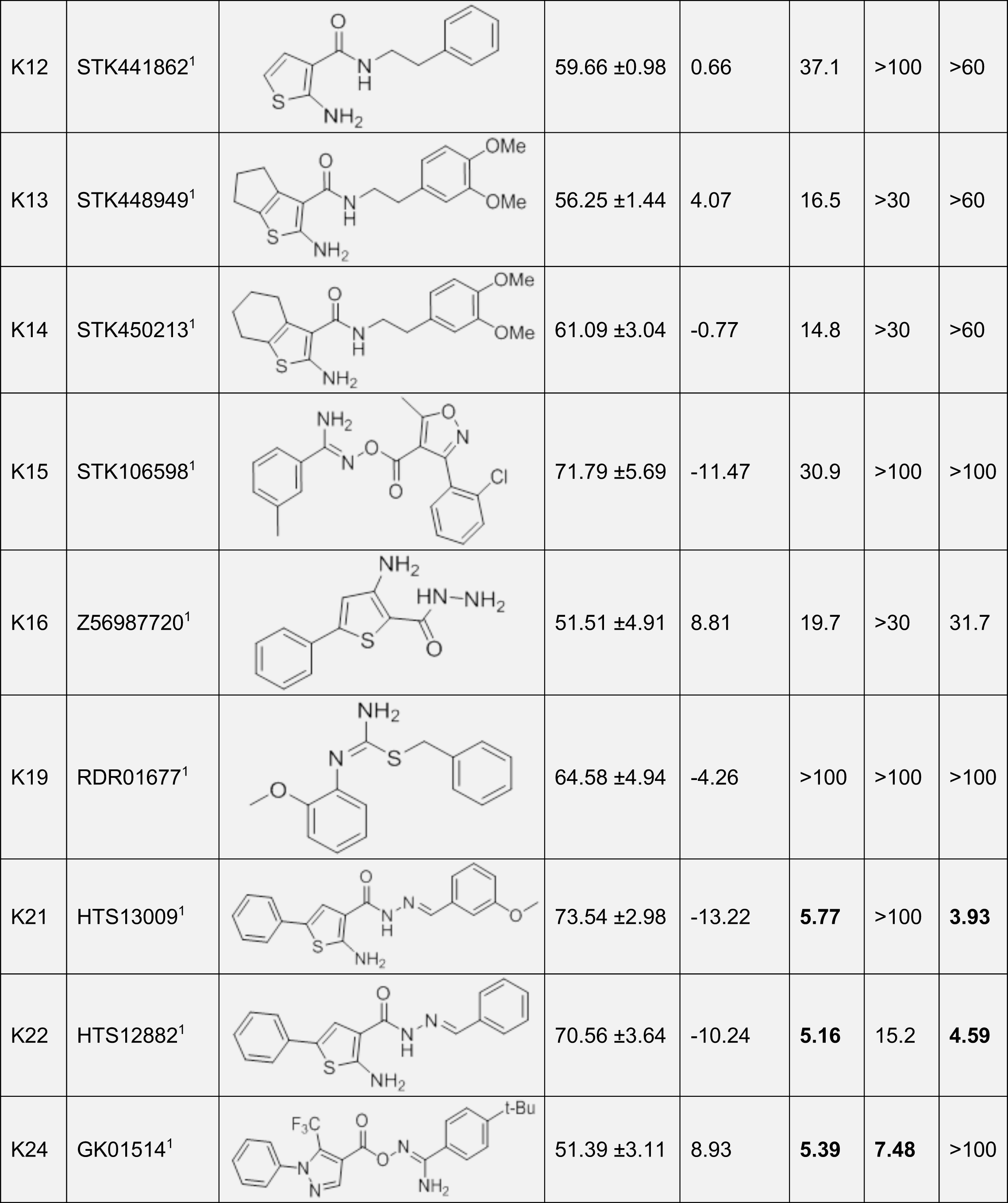
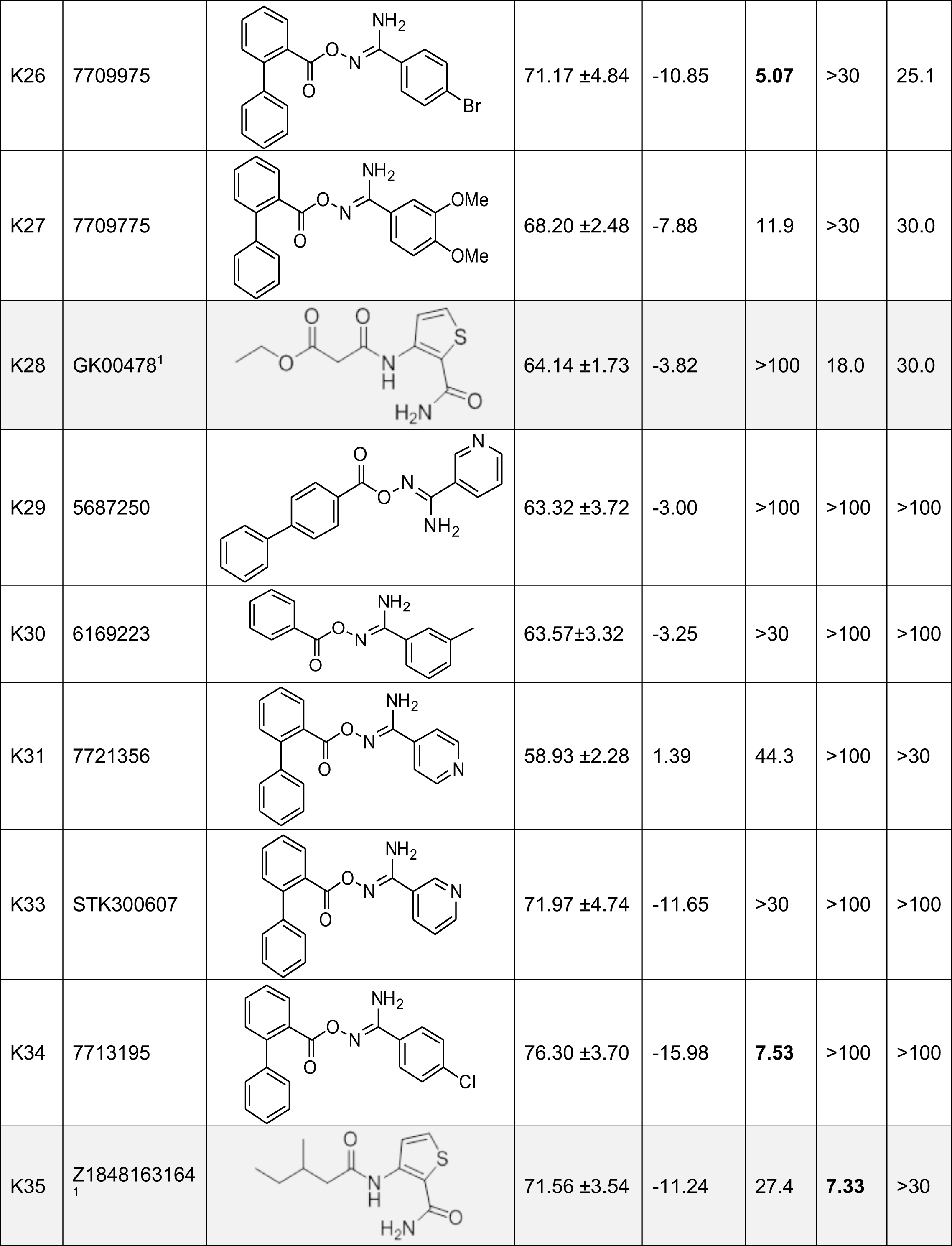
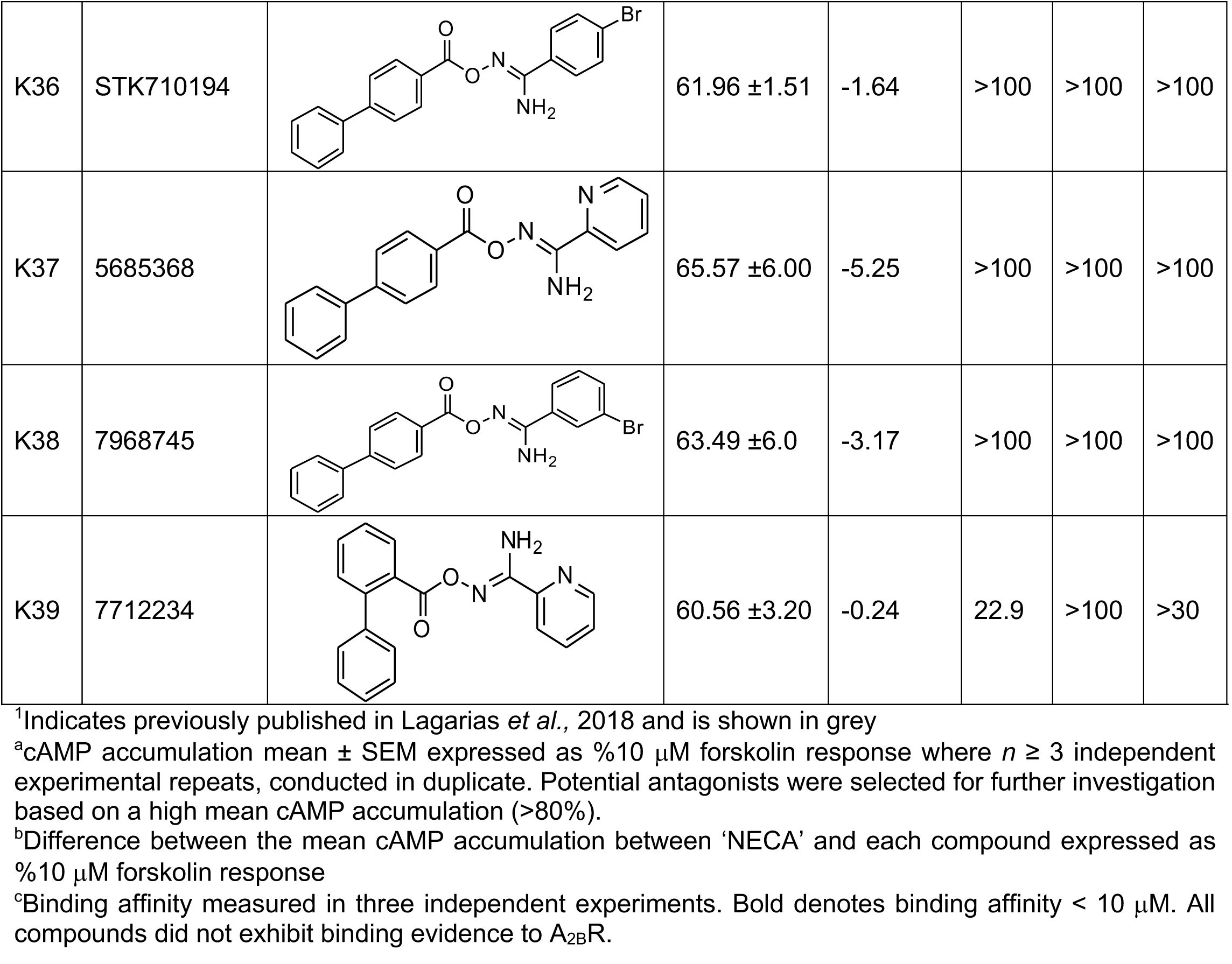
Compounds with no apparent antagonist/agonist activity at the A_3_R. Mean cAMP accumulation as measured in Flp-In CHO cells stably expressing A_3_R following stimulation with 10 μM forskolin only (DMSO) or NECA at the predetermined IC_80_ concentration and 1 μM test compound/DMSO control. Binding affinities were obtained through radioligand binding assays and chemical structures of new compounds tested against the A_1_R, A_2A_R and A_3_R are also included.

|  |  |  | A<sub>3</sub>R Flp-In CHO |  | Radioligand binding |  |  |
| --- | --- | --- | --- | --- | --- | --- | --- |
| | | | | | Ki (μM)<sup>c</sup> | | |
|  | Compound Name | Chemical structure | Mean <sup>a</sup> | Mean Change <sub>b</sub> | A<sub>3</sub>R | A<sub>1</sub>R | A<sub>2A</sub>R |
| | NECA | | 60.32 ± 3.41 | - | - | - | - |
| | DMSO | | 100.00 ± 1.15 | -35.73 | - | - | - |
| K2 | S05993<sup>1</sup> | | 68.28 ± 2.94 | -7.96 | 16.6 | >100 | 61.3 |
| K3 | SEW01061<sup>1</sup> | | 63.37 ± 2.66 | -3.05 | >100 | >100 | >100 |
| K4 | SPB06895<sup>1</sup> | | 59.06 ± 0.13 | 1.26 | >100 | >100 | >100 |
| K5 | SPB02733<sup>1</sup> | | 72.65 ± 7.16 | -12.33 | **9.45** | >100 | 21.8 |
| K6 | KM08495<sup>1</sup> | | 64.01 ± 5.22 | -3.69 | 30.6 | >100 | >100 |
| K7 | HTS06244<sup>1</sup> | | 61.96 ± 2.31 | -1.64 | 18.3 | >100 | >100 |
| K9 | STK323059<sup>1</sup> | | 71.40 ± 3.37 | 8.75 | **4.13** | **6.91** | >100 |
| K12 | STK441862<sup>1</sup> |  | 59.66 ±0.98 | 0.66 | 37.1 | >100 | >60 |
| K13 | STK448949<sup>1</sup> |  | 56.25 ±1.44 | 4.07 | 16.5 | >30 | >60 |
| K14 | STK450213<sup>1</sup> |  | 61.09 ±3.04 | -0.77 | 14.8 | >30 | >60 |
| K15 | STK106598<sup>1</sup> |  | 71.79 ±5.69 | -11.47 | 30.9 | >100 | >100 |
| K16 | Z56987720<sup>1</sup> |  | 51.51 ±4.91 | 8.81 | 19.7 | >30 | 31.7 |
| K19 | RDR01677<sup>1</sup> |  | 64.58 ±4.94 | -4.26 | >100 | >100 | >100 |
| K21 | HTS13009<sup>1</sup> |  | 73.54 ±2.98 | -13.22 | **5.77** | >100 | **3.93** |
| K22 | HTS12882<sup>1</sup> |  | 70.56 ±3.64 | -10.24 | **5.16** | 15.2 | **4.59** |
| K24 | GK01514<sup>1</sup> |  | 51.39 ±3.11 | 8.93 | **5.39** | **7.48** | >100 |
| K26 | 7709975 |  | 71.17 ±4.84 | -10.85 | **5.07** | >30 | 25.1 |
| K27 | 7709775 |  | 68.20 ±2.48 | -7.88 | 11.9 | >30 | 30.0 |
| K28 | GK00478<sup>1</sup> |  | 64.14 ±1.73 | -3.82 | >100 | 18.0 | 30.0 |
| K29 | 5687250 |  | 63.32 ±3.72 | -3.00 | >100 | >100 | >100 |
| K30 | 6169223 |  | 63.57±3.32 | -3.25 | >30 | >100 | >100 |
| K31 | 7721356 |  | 58.93 ±2.28 | 1.39 | 44.3 | >100 | >30 |
| K33 | STK300607 |  | 71.97 ±4.74 | -11.65 | >30 | >100 | >100 |
| K34 | 7713195 |  | 76.30 ±3.70 | -15.98 | **7.53** | >100 | >100 |
| K35 | Z1848163164<sub>1</sub> |  | 71.56 ±3.54 | -11.24 | 27.4 | **7.33** | >30 |
| K36 | STK710194 |  | 61.96 ±1.51 | -1.64 | >100 | >100 | >100 |
| K37 | 5685368 |  | 65.57 ±6.00 | -5.25 | >100 | >100 | >100 |
| K38 | 7968745 |  | 63.49 ±6.0 | -3.17 | >100 | >100 | >100 |
| K39 | 7712234 |  | 60.56 ±3.20 | -0.24 | 22.9 | >100 | >30 |

<sup>1</sup>Indicates previously published in Lagarias *et al.*, 2018 and is shown in grey
<sup>a</sup>cAMP accumulation mean ± SEM expressed as %10 μM forskolin response where $n \geq 3$ independent experimental repeats, conducted in duplicate. Potential antagonists were selected for further investigation based on a high mean cAMP accumulation (>80%).
<sup>b</sup>Difference between the mean cAMP accumulation between 'NECA' and each compound expressed as %10 μM forskolin response
<sup>c</sup>Binding affinity measured in three independent experiments. Bold denotes binding affinity < 10 μM. All compounds did not exhibit binding evidence to A<sub>2B</sub>R.

**Supplementary Figure 2.**
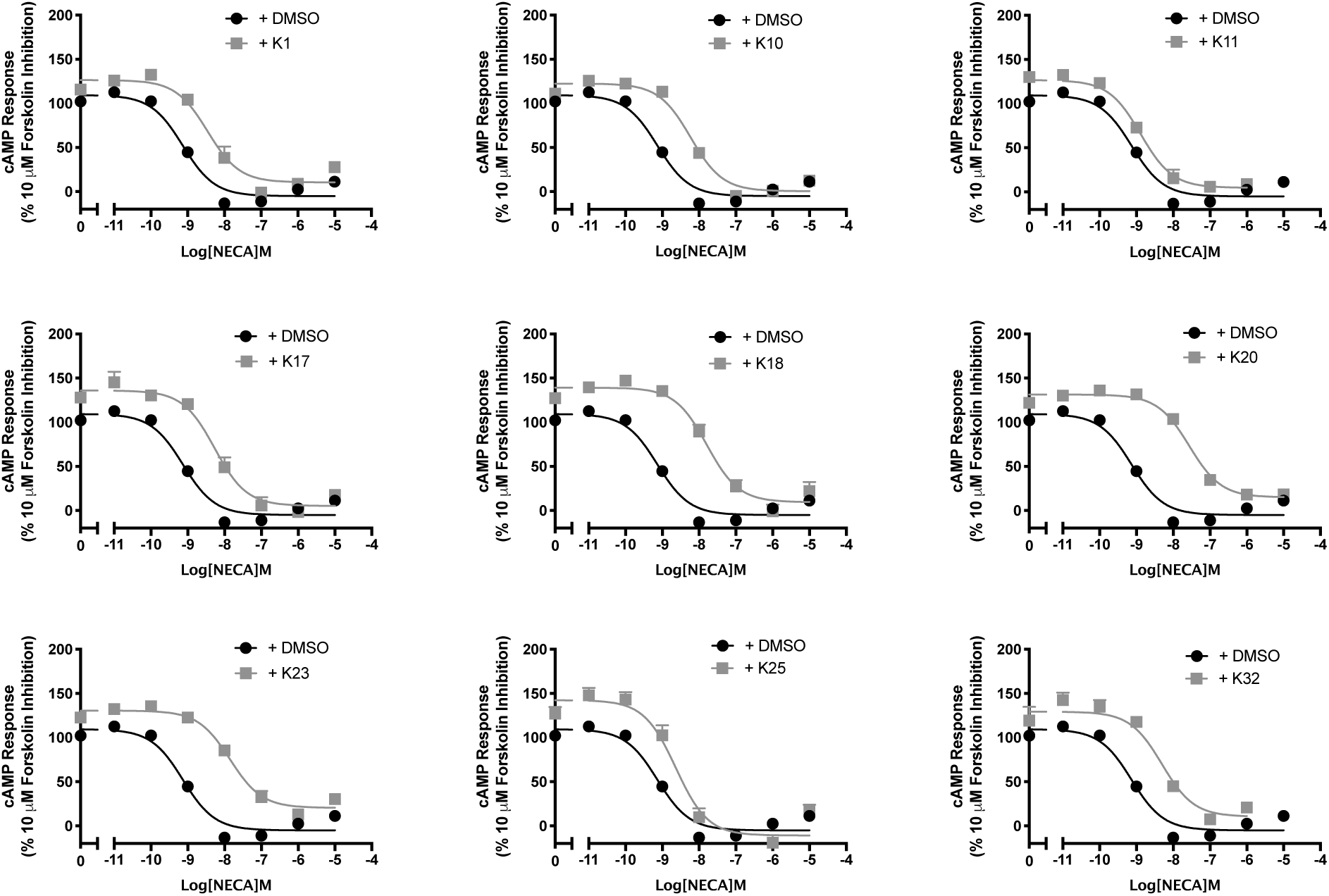
Screening of potential antagonists at the A_3_R. A_3_R Flp-In CHO cells (2000 cells/well) were exposed to forskolin 10 μM, NECA and DMSO or 1 μM test compound for 30 min and cAMP accumulation detected. All values are mean ± SEM expressed as percentage forskolin inhibition (10 μM) relative to NECA. *n* ≥ 3 independent experimental repeats, conducted in duplicate.

**Supplementary Table 2.** cAMP accumulation as measured in A_3_R Flp-In CHO cells following stimulation with 10 μM forskolin, varying concentrations of NECA and 1 μM test compound/DMSO control

| **A<sub>3</sub>R Flp-In CHO** |  |  |  |  |  |
| --- | --- | --- | --- | --- | --- |
|  | **pIC<sub>50</sub><sup>a</sup>** | **E<sub>min</sub><sup>b</sup>** | **Basal<sup>c</sup>** | **True Basal<sup>d</sup>** | **Span<sup>e</sup>** |
| **NECA only** | 8.94 ±0.1 | -5.02 ±2.4 | 109.6 ±1.9 | 100.0 ±1.7 | 110.4 ±2.2 |
| **K1** | 8.46 ±0.1 | 10.5 ±4.5 | 126.9 ±3.3 | 115.7 ±2.8 | 118.0 ±3.1 |
| **K10** | 8.23 ±0.1 | 0.5 ±3.2 | 122.6 ±0.6 | 111.0 ±2.0 | 122.1 ±1.2 |
| **K11** | 8.88 ±0.1 | 4.7 ±4.3 | 126.5 ±3.2 | 109.1 ±2.0 | 121.9 ±5.2 |
| **K17** | 8.27 ±0.1 | 5.2 ±4.6 | 135.7 ±5.4 | 128.1 ±2.1 | 131.7 ±0.8 |
| **K18** | 7.79 ±0.1 | 9.5 ±4.8 | 139.2 ±3.7 | 127.2 ±3.2 | 130.9 ±7.1 |
| **K20** | 7.56 ±0.1 | 14.9 ±3.1 | 131.4 ±2.2 | 127.0 ±3.2 | 116.5 ±5.4 |
| **K23** | 7.85 ±0.1 | 20.4 ±3.2 | 130.5 ±2.6 | 122.7 ±3.9 | 110.7 ±2.7 |
| **K25** | 8.63 ±0.1 | -10.9 ±5.7 | 136.6 ±7.1 | 138.5 ±4.9 | 134.4 ±5.8 |
| **K32** | 8.31 ±0.1 | 10.5 ±7.1 | 129.4 ±4.5 | 113.1 ±5.6 | 118.9 ±7.1 |

<sup>a</sup>Negative logarithm of NECA concentration required to produce a half-maximal response in the absence (DMSO) or presence of 1 μM compound
<sup>b</sup>Minimum cAMP accumulation of NECA as % 10 μM forskolin response, relative to NECA; the lower plateau of the fitted sigmoidal dose response curve
<sup>c</sup>The upper plateau of the fitted sigmoidal dose response curve corresponding to 100% of the 1 μM forskolin response
<sup>d</sup>The cAMP accumulation when stimulated with 1 μM compound and 10 μM forskolin stimulation only
<sup>e</sup>The difference between E<sub>min</sub> and basal signalling

**Supplementary Figure 3.**
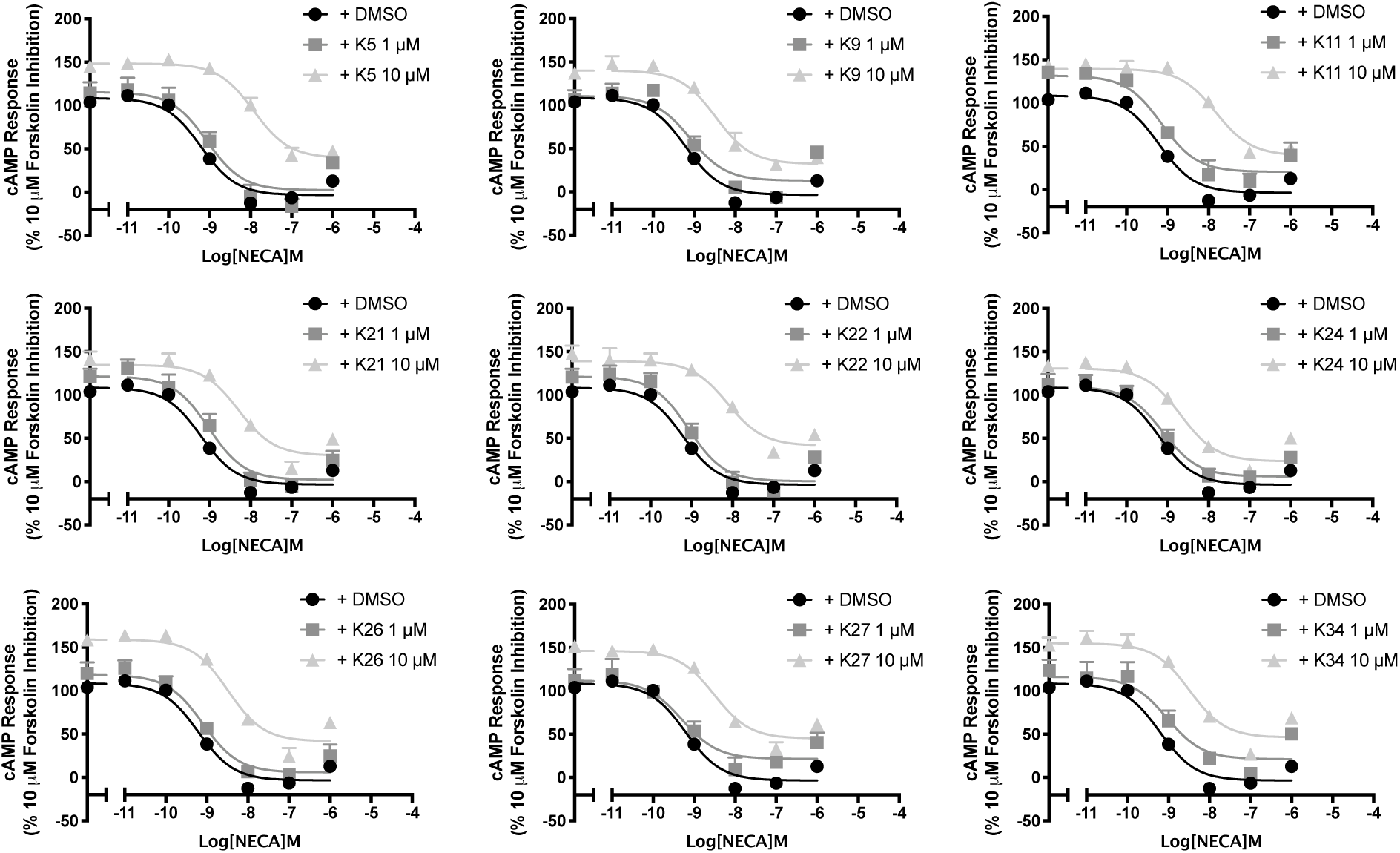
Determining functional activity of compounds with a micromolar binding affinity for A_3_R. A_3_R Flp-In CHO cells (2000 cells/well) were exposed to forskolin 10 μM, NECA and DMSO or test compound at the indicated concentration for 30 min and cAMP accumulation determined. All values are mean ± SEM expressed as percentage forskolin inhibition (10 μM) relative to NECA. *n* = 3 independent experimental repeats, conducted in duplicate.

**Supplementary Table 3.** cAMP accumulation as measured in A_3_R Flp-In CHO cells following stimulation with 10 μM forskolin, varying concentrations of NECA and 1 μM or 10 μM test compound/DMSO control

| **A<sub>3</sub>R Flp-In CHO** |  |  |  |  |  |  |
| --- | --- | --- | --- | --- | --- | --- |
|  |  | **pIC<sub>50</sub><sup>a</sup>** | **E<sub>min</sub><sup>b</sup>** | **Basal<sup>c</sup>** | **True Basal<sup>d</sup>** | **Span<sup>e</sup>** |
| **NECA only** | | 8.94 ± 0.1 | -5.02 ± 2.4 | 109.6 ± 1.9 | 100.0 ± 1.7 | 110.4 ± 2.2 |
| **K5** | 1 μM | 9.06 ± 0.2 | 2.3 ± 8.2 | 115.3 ± 7.0 | 102.5 ± 10.1 | 113.1 ± 10.5 |
| | 10 μM | 7.93 ± 0.1 | 39.3 ± 5.6 | 148.3 ± 3.1 | 142.1 ± 9.3 | 108.9 ± 6.2 |
| **K9** | 1 μM | 9.09 ± 0.2 | 13.0 ± 7.6 | 110.7 ± 6.6 | 86.2 ± 6.0 | 97.8 ± 9.7 |
| | 10 μM | 8.47 ± 0.2 | 32.3 ± 5.7 | 140.0 ± 3.9 | 123.3 ± 3.8 | 107.7 ± 6.7 |
| **K11** | 1 μM | 8.88 ± 0.1 | 4.7 ± 4.3 | 126.5 ± 3.2 | 109.1 ± 2.0 | 121.9 ± 5.2 |
| | 10 μM | 7.83 ± 0.2 | 39.4 ± 7.5 | 139.3 ± 5.0 | 119.8 ± 9.2 | 99.9 ± 8.2 |
| **K21** | 1 μM | 9.02 ± 0.2 | 2.0 ± 7.6 | 121.5 ± 6.1 | 101.6 ± 5.1 | 119.4 ± 9.4 |
| | 10 μM | 8.29 ± 0.2 | 30.1 ± 6.8 | 134.7 ± 4.3 | 118.6 ± 4.8 | 104.6 ± 7.7 |
| **K22** | 1 μM | 9.07 ± 0.2 | 0.4 ± 7.1 | 121.3 ± 5.8 | 101.9 ± 8.4 | 120.9 ± 8.8 |
| | 10 μM | 8.12 ± 0.2 | 41.6 ± 7.1 | 139.1 ± 4.2 | 124.5 ± 8.2 | 97.5 ± 8.0 |
| **K24** | 1 μM | 9.12 ± 0.2 | 5.9 ± 6.5 | 109.5 ± 4.9 | 103.6 ± 7.7 | 103.7 ± 7.9 |
| | 10 μM | 8.67 ± 0.1 | 23.4 ± 4.5 | 130.8 ± 3.3 | 117.7 ± 0.9 | 107.4 ± 5.7 |
| **K26** | 1 μM | 9.10 ± 0.2 | 5.7 ± 6.2 | 118.2 ± 4.8 | 98.9 ± 8.8 | 112.5 ± 7.6 |
| | 10 μM | 8.49 ± 0.1 | 41.5 ± 5.7 | 158.8 ± 3.9 | 143.0 ± 5.9 | 117.3 ± 6.6 |
| **K27** | 1 μM | 9.29 ± 0.2 | 21.5 ± 7.4 | 112.0 ± 6.8 | 93.1 ± 6.7 | 90.5 ± 9.7 |
| | 10 μM | 8.51 ± 0.2 | 44.9 ± 5.3 | 146.3 ± 3.7 | 133.2 ± 3.0 | 101.4 ± 6.2 |
| **K34** | 1 μM | 9.04 ± 0.2 | 21.1 ± 8.7 | 116.2 ± 6.9 | 97.1 ± 2.1 | 95.1 ± 10.8 |
| | 10 μM | 8.49 ± 0.2 | 46.2 ± 6.7 | 154.9 ± 4.7 | 143.6 ± 7.6 | 108.7 ± 7.8 |

<sup>a</sup>Negative logarithm of NECA concentration required to produce a half-maximal response in the absence (DMSO) or presence of 1 μM or 10 μM compound
<sup>b</sup>Minimum cAMP accumulation of NECA as % 10 μM forskolin response, relative to NECA; the lower plateau of the fitted sigmoidal dose response curve
<sup>c</sup>The upper plateau of the fitted sigmoidal dose response curve corresponding to 100% of the 10 μM forskolin response
<sup>d</sup>The cAMP accumulation when stimulated with 1 μM or 10 μM compound and 10 μM forskolin stimulation only
<sup>e</sup>The difference between E<sub>min</sub> and basal signalling

**Supplementary Figure 4.**
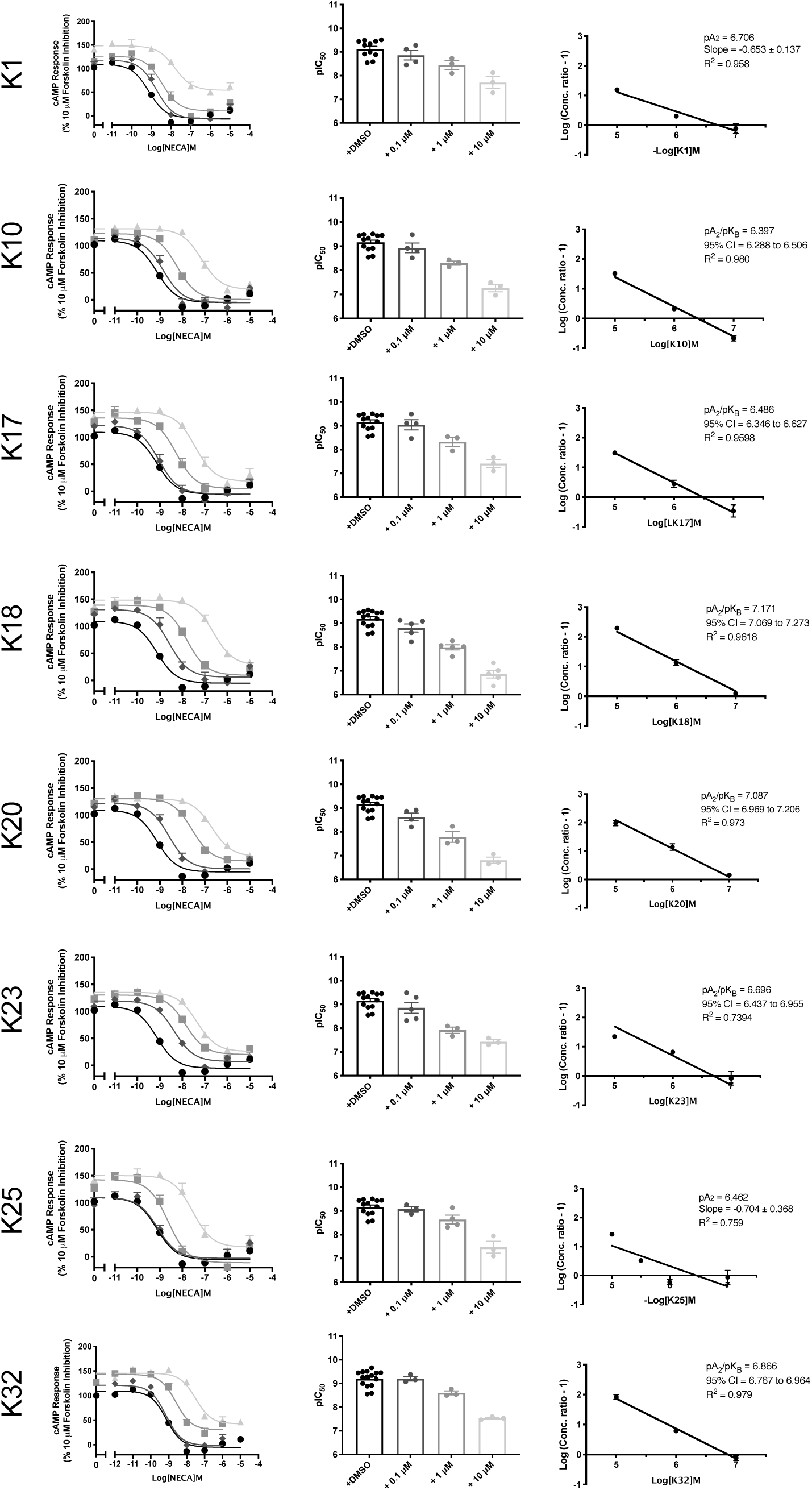
NECA stimulated cAMP inhibition at WT A_3_R: activity of potential antagonists. Flp-In-CHO cells (2000 cells/well) stably expressing WT A_3_R were exposed to forskolin 10 μM, NECA and test compound/DMSO control for 30 min and cAMP accumulation detected. **A**) Representative dose response curves are shown as mean ± SEM expressed as percentage forskolin inhibition (10 μM) relative to NECA. **B**) pIC_50_ values are shown as mean ± SEM. **C**) Schild analysis of data represented in **A/B**. A slope of 1 indicates a competitive antagonist. The x-axis is expressed as -log (molar concentration of antagonist) giving a negative Schild plot slope.

**Supplementary Table 4.** cAMP accumulation as measured in Flp-In-CHO stably expressing A_3_R following stimulation with 10 μM forskolin, compound at the indicated concentration and varying concentrations of NECA

| WT A<sub>3</sub>R Flp-In-CHO |  |  |  |  |  |  |
| --- | --- | --- | --- | --- | --- | --- |
|  |  | pIC<sub>50</sub><sup>a</sup> | E<sub>min</sub><sup>b</sup> | Basal<sup>c</sup> | True Basal<sup>d</sup> | Span<sup>e</sup> |
| **NECA only** | | 8.94 ± 0.1 | -5.02 ± 2.4 | 109.6 ± 1.9 | 100.0 ± 1.7 | 110.4 ± 2.2 |
| **K1** | 0.1 μM | 8.73 ± 0.2 | -8.38 ± 5.4 | 118.3 ± 3.3 | 107.2 ± 3.0 | 117.0 ± 2.5 |
| | 1 μM | 8.46 ± 0.1 | 10.5 ± 4.5 | 126.9 ± 3.3 | 115.7 ± 2.8 | 118.0 ± 3.1 |
| | 10 μM | 7.80 ± 0.1 | 54.1 ± 4.5 | 148.5 ± 6.6 | 145.7 ± 2.4 | 96.9 ± 3.3* |
| **K10** | 0.1 μM | 8.76 ± 0.2 | -6.0 ± 5.6 | 112.9 ± 1.9 | 98.2 ± 3.1 | 109.7 ± 2.8 |
| | 1 μM | 8.23 ± 0.1 | 0.54 ± 3.2 | 122.6 ± 0.6 | 111.0 ± 2.0 | 122.1 ± 1.2 |
| | 10 μM | 7.15 ± 0.1 | 19.2 ± 4.5 | 131.7 ± 1.8 | 121.9 ± 2.6 | 112.6 ± 2.6 |
| **K17** | 0.1 μM | 9.00 ± 0.1 | -5.5 ± 4.8 | 122.6 ± 1.9 | 115.0 ± 4.4 | 124.6 ± 2.5 |
| | 1 μM | 8.27 ± 0.1 | 5.2 ± 4.6 | 135.7 ± 5.4 | 128.1 ± 2.1 | 131.7 ± 0.8 |
| | 10 μM | 7.43 ± 0.1 | 18.7 ± 5.2 | 146.6 ± 3.3 | 138.6 ± 3.1 | 131.2 ± 4.0 |
| **K18** | 0.1 μM | 8.58 ± 0.1 | 6.1 ± 5.5 | 131.1 ± 5.4 | 122.3 ± 2.1 | 127.5 ± 4.7 |
| | 1 μM | 7.79 ± 0.1 | 9.5 ± 4.8 | 139.2 ± 3.7 | 127.2 ± 3.2 | 130.9 ± 7.1 |
| | 10 μM | 6.61 ± 0.1 | 28.3 ± 5.5 | 148.5 ± 2.6 | 143.1 ± 1.3 | 121.7 ± 1.4 |
| **K20** | 0.1 μM | 8.38 ± 0.1 | 2.2 ± 4.1 | 119.6 ± 3.9 | 122.3 ± 2.1 | 117.9 ± 3.4 |
| | 1 μM | 7.56 ± 0.1 | 14.9 ± 3.1 | 131.4 ± 2.2 | 127.0 ± 3.2 | 116.5 ± 5.4 |
| | 10 μM | 6.68 ± 0.1 | 23.6 ± 3.8 | 130.2 ± 1.9 | 143.1 ± 1.3 | 106.8 ± 5.5 |
| **K23** | 0.1 μM | 8.36 ± 0.1 | 7.5 ± 3.4 | 119.5 ± 3.2 | 117.3 ± 3.2 | 112.2 ± 1.3 |
| | 1 μM | 7.85 ± 0.1 | 20.4 ± 3.2 | 130.5 ± 2.6 | 122.7 ± 3.9 | 110.7 ± 2.7 |
| | 10 μM | 7.35 ± 0.1 | 25.9 ± 3.5 | 135.3 ± 2.3 | 129.3 ± 4.9 | 109.2 ± 7.3 |
| **K25** | 0.1 μM | 9.10 ± 0.2 | -2.7 ± 6.2 | 122.0 ± 7.0 | 126.7 ± 6.5 | 126.6 ± 4.4 |
| | 1 μM | 8.63 ± 0.1 | -10.9 ± 5.7 | 136.6 ± 7.1 | 138.5 ± 4.9 | 134.4 ± 5.8 |
| | 10 μM | 7.54 ± 0.2 | 17.9 ± 7.0 | 151.7 ± 3.5 | 148.5 ± 5.4 | 136.5 ± 2.5 |
| **K32** | 0.1 μM | 9.22 ± 0.1 | -1.3 ± 4.9 | 124.5 ± 5.4 | 107.4 ± 5.4 | 114.9 ± 8.4 |
| | 1 μM | 8.62 ± 0.1 | 30.5 ± 4.9 | 144.8 ± 4.1 | 132.3 ± 4.3 | 115.4 ± 6.8 |
| | 10 μM | 7.53 ± 0.1 | 42.6 ± 3.0 | 144.7 ± 2.9 | 126.5 ± 3.0 | 91.7 ± 6.8 |

<sup>a</sup>Negative logarithm of NECA concentration required to produce a half-maximal response in the absence (NECA only) or presence of 0.1, 1 or 10 μM compound
<sup>b</sup>Minimum cAMP accumulation of NECA as % 10 μM forskolin response relative to NECA response; The lower plateau of the fitted sigmoidal dose response curve
<sup>c</sup>The upper plateau of the fitted sigmoidal dose response curve corresponding to % 10 μM forskolin inhibition, relative to NECA
<sup>d</sup>The cAMP accumulation when stimulated with compound at the indicated concentration and 10 μM forskolin stimulation only
<sup>e</sup>The difference between E<sub>min</sub> and basal signaling

**Supplementary Figure 5.**
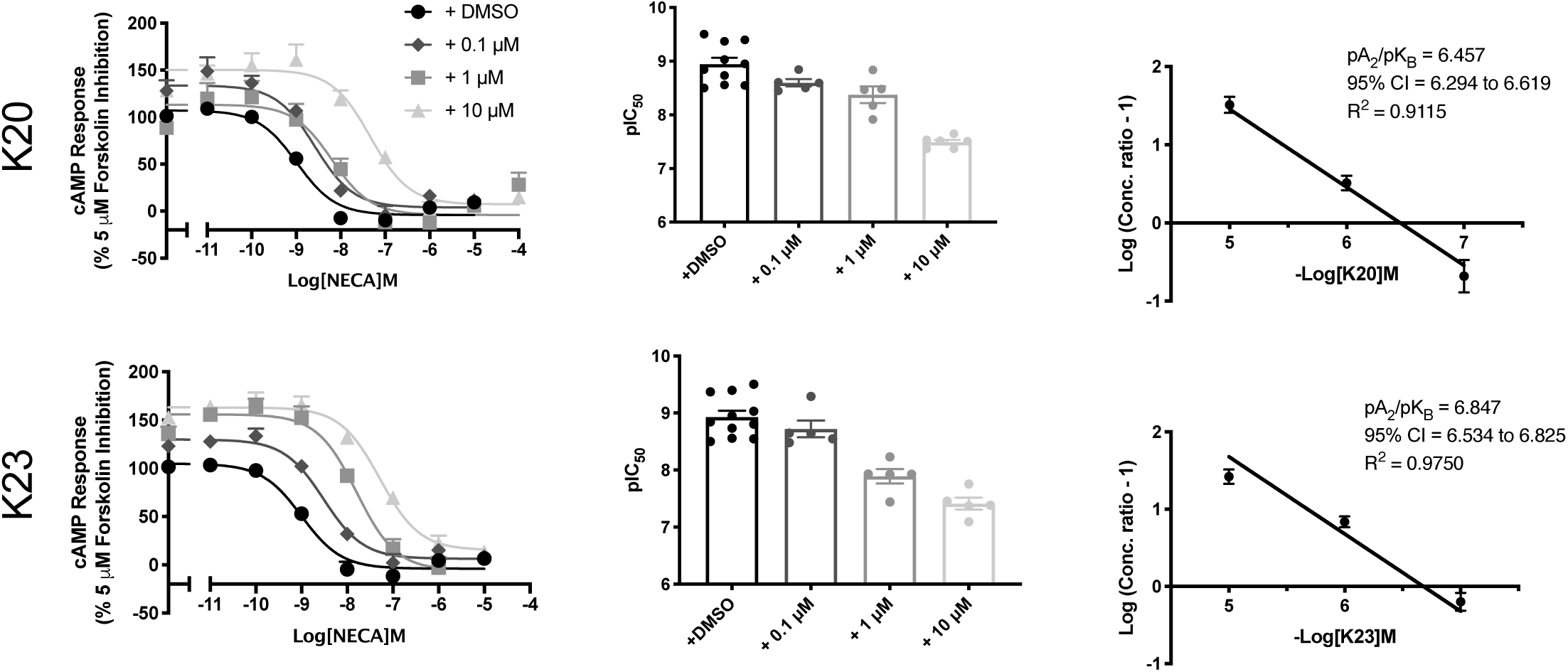
NECA stimulated cAMP inhibition at A_1_R. CHO-K1 cells (2000 cells/well) stably expressing A_1_R were exposed to forskolin 5 μM, NECA and test compound/DMSO control for 30 min and cAMP accumulation detected. **A**) Representative dose response curves are shown as mean ± SEM expressed as percentage forskolin inhibition (5 μM) relative to NECA. **B**) pIC_50_ values for individual repeats including half-log concentration are shown as mean ± SEM. **C**) Schild analysis of data represented in **A/B**. A slope of 1 indicates a competitive antagonist. The x-axis is expressed as -log (molar concentration of antagonist) giving a negative Schild plot slope.

**Supplementary Table 5.** cAMP accumulation as measured in CHO-K1 stably expressing A_1_R following stimulation with 5 μM forskolin, compound at the indicated concentration and varying concentrations of NECA

| **A<sub>1</sub>R CHO-K1** |  |  |  |  |  |  |
| --- | --- | --- | --- | --- | --- | --- |
|  |  | **pIC<sub>50</sub><sup>a</sup>** | **E<sub>min</sub><sup>b</sup>** | **Basal<sup>c</sup>** | **True Basal<sup>d</sup>** | **Span<sup>e</sup>** |
| **NECA only** | | 8.94 ± 0.1 | -5.1 ± 1.0 | 107.5 ± 3.4 | 100.0 ± 1.16 | 111.4 ± 4.0 |
| **K20** | 0.1 μM | 8.68 ± 0.1 * | -5.3 ± 9.3 | 130.1 ± 2.6 | 110.9 ± 4.4 | 135.4 ± 6.7 |
| | 1 μM | 8.37 ± 0.1 **** | -3.7 ± 3.8 | 106.6 ± 8.5 | 113.6 ± 18.1 | 98.6 ± 5.6 |
| | 10 μM | 7.38 ± 0.1 **** | 8.1 ± 2.0** | 136.6 ± 2.3** | 140.7 ± 6.4* | 153.5 ± 12.1*** |
| **K23** | 0.1 μM | 8.52 ± 0.1 **** | 3.3 ± 5.6 | 131.6 ± 5.1 | 117.6 ± 3.2 | 128.3 ± 7.2 |
| | 1 μM | 8.00 ± 0.1 **** | 0.1 ± 6.3 | 132.8 ± 3.8* | 119.8 ± 3.2* | 149.5 ± 15.5* |
| | 10 μM | 7.49 ± 0.1 **** | 12.2 ± 4.7* | 178.3 ± 16.2**** | 170.2 ± 13.6**** | 166.0 ± 15.0** |

<sup>a</sup>Negative logarithm of NECA concentration required to produce a half-maximal response in the absence (NECA only) or presence of compound
<sup>b</sup>Minimum cAMP accumulation of NECA as % 5 μM forskolin response; the lower plateau of the fitted sigmoidal dose response curve
<sup>c</sup>The upper plateau of the fitted sigmoidal dose response curve corresponding to 100% of the 5 μM forskolin response
<sup>d</sup>The cAMP accumulation when stimulated with compound at the indicated concentration and 5 μM forskolin stimulation only
<sup>e</sup>The difference between E<sub>min</sub> and basal signalling
Statistical significance (\*, $p < 0.05$ ; \*\*, $p < 0.01$ ; \*\*\*, $p < 0.001$ ; \*\*\*\*, $p < 0.0001$ ) compared to NECA only stimulation was determined by one-way ANOVA with Dunnett's post-test.

**Supplementary Figure 6.**
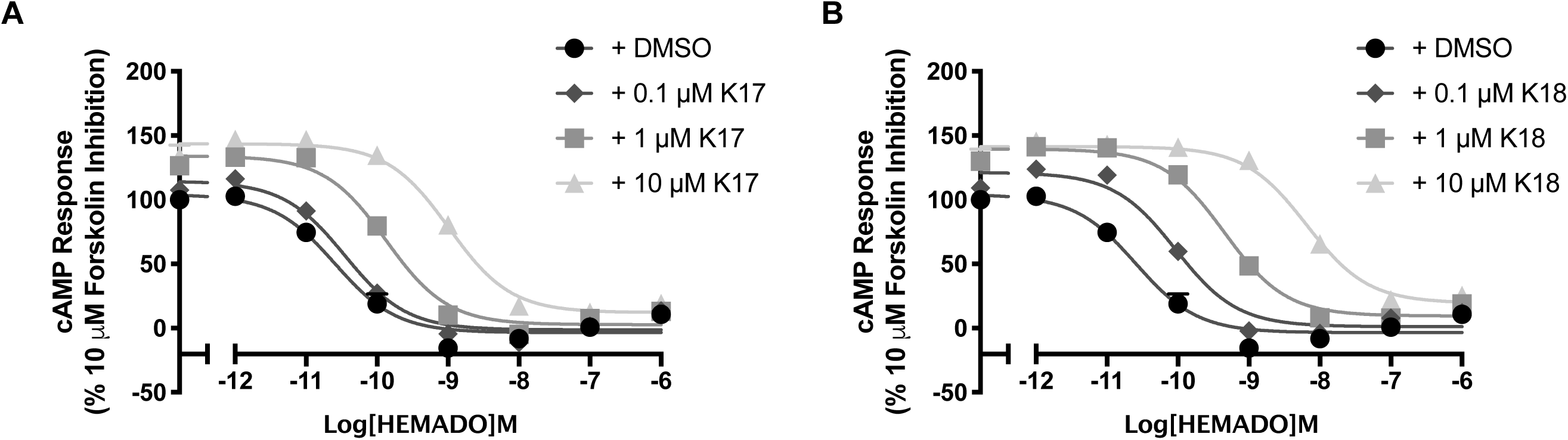
Antagonism of the HEMADO induced cAMP inhibition at the A_3_R of A) K17 and B) K18. A_3_R stably expressing Flp-In CHO cells (2000 cells/well) were exposed to forskolin 10 μM, HEMADO and DMSO/test compound at the indicated concentration for 30 min and cAMP accumulation detected. All values are mean ± SEM expressed as percentage forskolin inhibition (10 μM) relative to HEMADO. *n* = 3 independent experimental repeats, conducted in duplicate.

**Supplementary Table 6.** cAMP accumulation as measured in Flp-In-CHO cells stably expressing A_3_R following stimulation with 10 μM forskolin, HEMADO and varying concentrations of test compound

|  |  | **A<sub>3</sub>R CHO-K1** |  |  |  |  |
| --- | --- | --- | --- | --- | --- | --- |
|  |  | **pIC<sub>50</sub><sup>a</sup>** | **E<sub>min</sub><sup>b</sup>** | **Basal<sup>c</sup>** | **True Basal<sup>d</sup>** | **Span<sup>e</sup>** |
| **HEMADO only** | | 10.54 ± 0.2 | -3.48 ± 2.7 | 103.8 ± 4.1 | 100.1 ± 1.9 | 107.3 ± 4.7 |
| **K17** | 0.1 μM | 10.45 ± 0.1 | -1.31 ± 2.8 | 114.3 ± 4.0 | 107.6 ± 3.6 | 115.6 ± 4.6 |
| | 1 μM | 9.89 ± 0.1 | 2.71 ± 2.9 | 134.0 ± 3.3 | 126.4 ± 4.1 | 131.3 ± 4.3 |
| | 10 μM | 8.99 ± 0.1 | 12.16 ± 4.1 | 143.6 ± 3.4 | 133.3 ± 9.3 | 131.5 ± 5.2 |
| **K18** | 0.1 μM | 10.04 ± 0.1 | 1.20 ± 2.9 | 121.2 ± 3.5 | 109.1 ± 2.2 | 120.0 ± 4.4 |
| | 1 μM | 9.34 ± 0.1 | 9.52 ± 3.9 | 139.5 ± 2.8 | 130.1 ± 5.3 | 130.0 ± 3.9 |
| | 10 μM | 8.12 ± 0.1 | 19.6 ± 4.6 | 141.4 ± 2.8 | 130.2 ± 9.3 | 121.8 ± 5.3 |

<sup>a</sup>Negative logarithm of HEMADO concentration required to produce a half-maximal response in the absence (HEMADO only) or presence of compound
<sup>b</sup>Minimum cAMP accumulation of HEMADO as % 10 μM forskolin response; the lower plateau of the fitted sigmoidal dose response curve
<sup>c</sup>The upper plateau of the fitted sigmoidal dose response curve corresponding to 100% of the 10 μM forskolin response
<sup>d</sup>The cAMP accumulation when stimulated with compound at the indicated concentration and 10 μM forskolin stimulation only
<sup>e</sup>The difference between E<sub>min</sub> and basal signalling

**Supplementary Figure 7.**
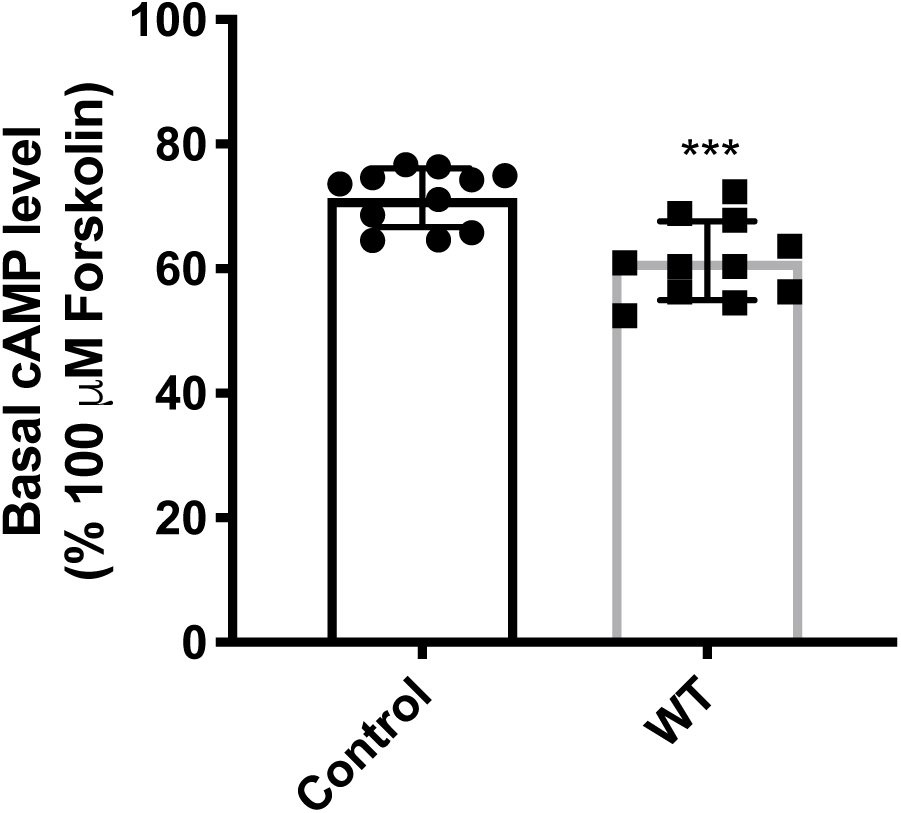
A_3_R shows constitutive activity. cAMP accumulation following a 30-minute stimulation with forskolin (5 μM and 10 μM) in WT A_3_R expressing Flp-In-CHO cells was reduced compared to control (Flp-In-CHO cells). Statistical significance (*\*\*\*, p<0.001*) compared to control was determined by Student’s t-test.

**Supplementary Figure 8.**
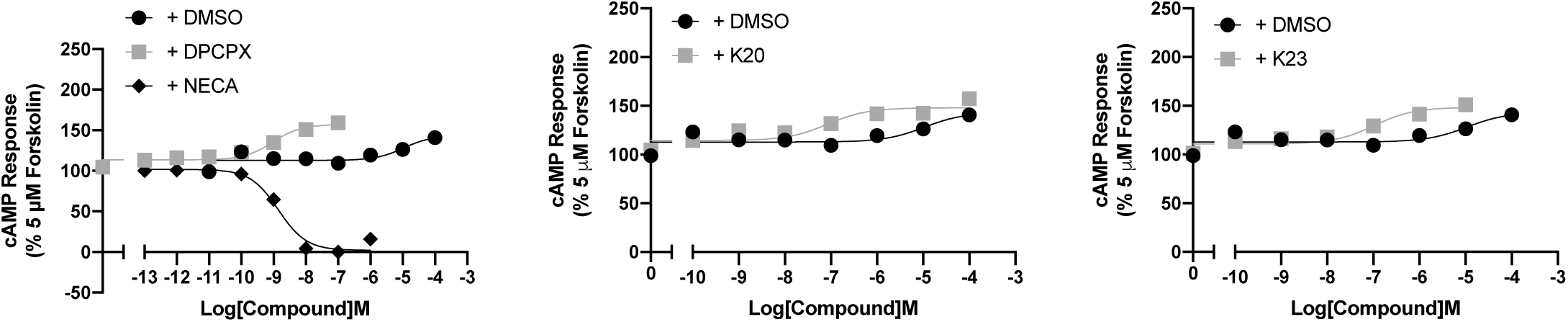
Inverse agonism at the A_1_R. cAMP accumulation following a 30-minute stimulation with forskolin (5 μM) and increasing concentrations of antagonist, NECA or DMSO control was determined in A_1_R expressing CHO cells. Representitive dose response curves are shown as mean ± SEM expressed as percentage forskolin (5 μM), relative to NECA.

**Supplementary Figure 9.**
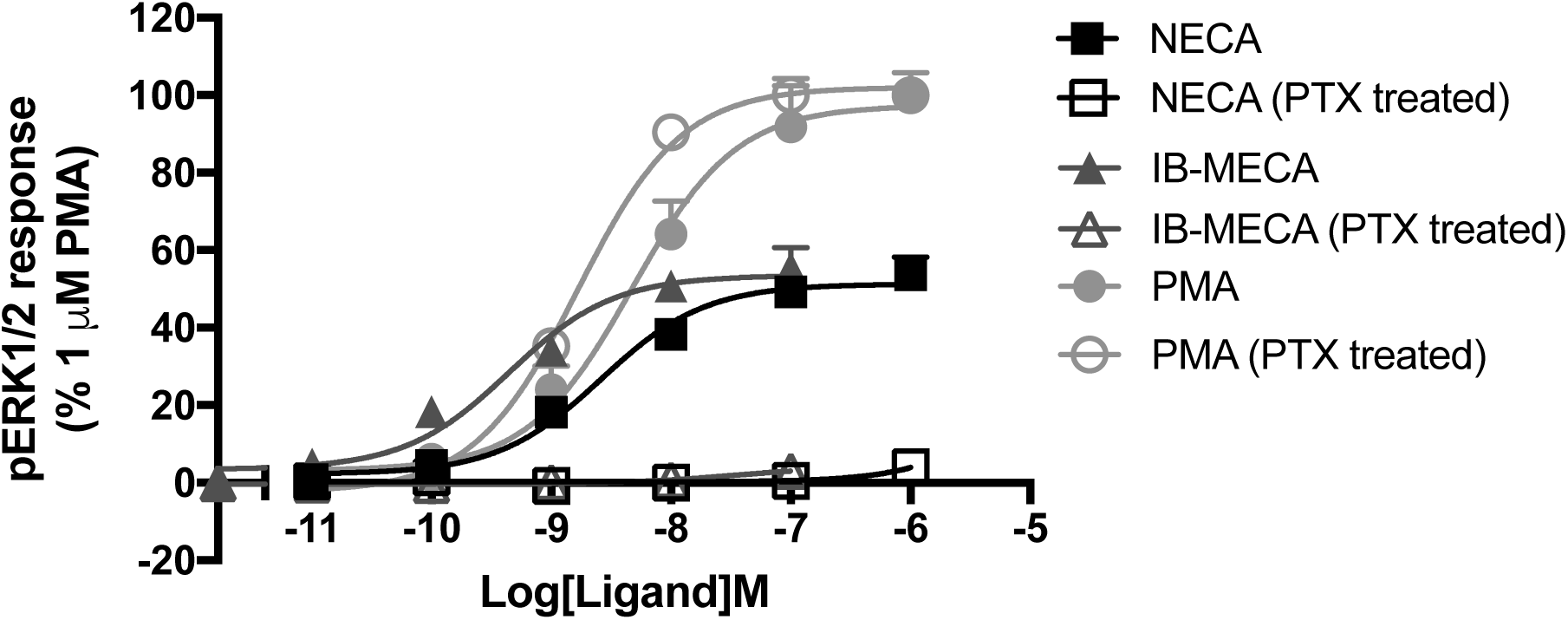
A_3_R stimulated pERK activity is entirely G_i/o_ mediated. pERK was detected in Flp-In-CHO cells stably expressing A_3_R (2000 cells/well) stimulated for 5 minutes with NECA or IB-MECA with or without Pertussis toxin (PTX) treatment (16 hours at 100 ng/mL). All values are mean ± SEM expressed as % 1μM PMA response where *n* =3 for none-PTX treated and *n* = 1 for PTX treated, conducted in duplicate.

**Supplementary Figure 10.**
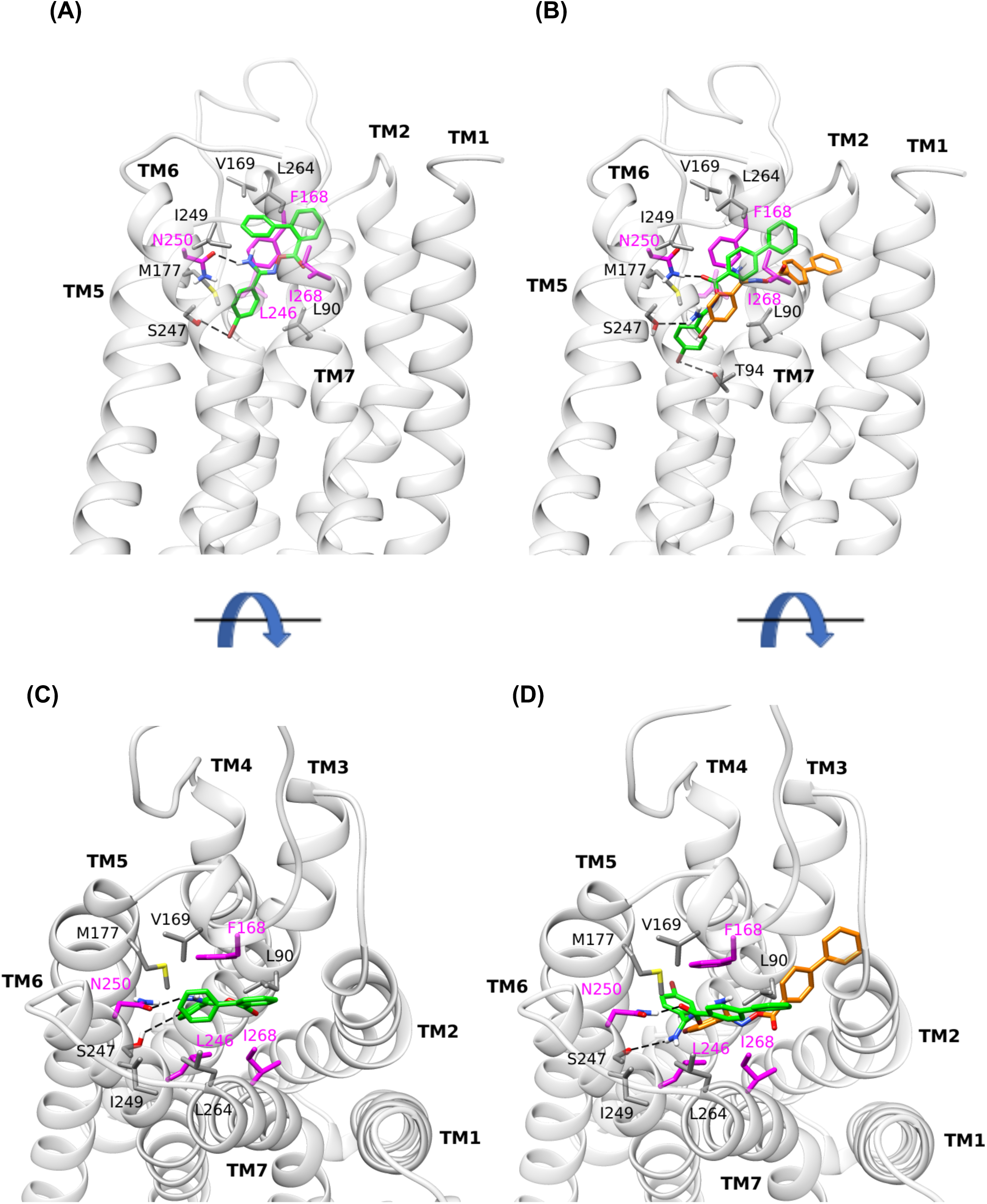
Average structure of WT A_3_R in complex with K26 and K36 from MD simulations with Amber14ff. Side **(A)**, top **(C)** view of the average structure of K26 which binds A_3_R (*K*_d_ =5.07 μM) but has not antagonistic activity and, side **(B)**, top **(D**) view of the average structure of K36 which did not bind A_3_R inside the orthosteric binding area from 100 MD simulations. Side chains of critical residues for binding resulted from the MD simulations are shown in sticks. Residues L90^3.32^, V169^5.30^, M177^5.40^, I249^6.54^, L264^7.34^, in which carbon atoms are shown in grey, were confirmed experimentally; in residues F168^5.29^, L246^6.51^, I268^7.39^ and N250^6.55^ carbon atoms are shown in magenta; nitrogen, oxygen and sulfur atoms are shown in blue, red and yellow respectively. For K36 the conformation adopted after 100 ns, which loses binding interactions with the receptor area, is indicated with orange colour for carbons.

**Supplementary Figure 11.**
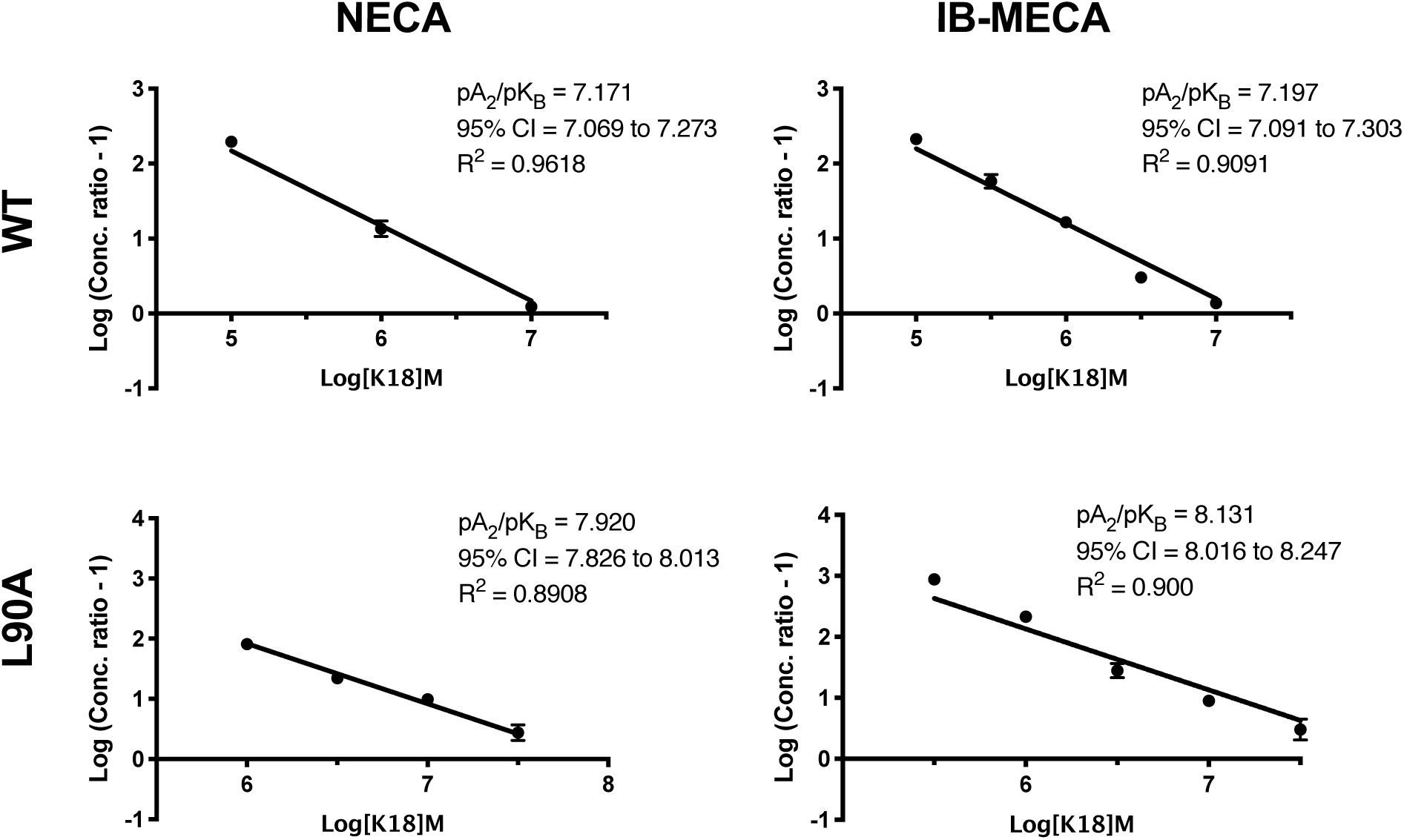
pA_2_ values obtained through Schild analysis are agonist independent. Flp-In-CHO cells (2000 cells/well) stably expressing WT or L90A^3.32^ A_3_R were exposed to forskolin 10 μM, agonist (NECA or IB-MECA) and K18 at varying concentrations for 30 min and cAMP accumulation detected. IC_50_ values determined through fitting three-parameter logistic equation to concentration response data were used to conduct Schild analysis.

**Supplementary Figure 12.**
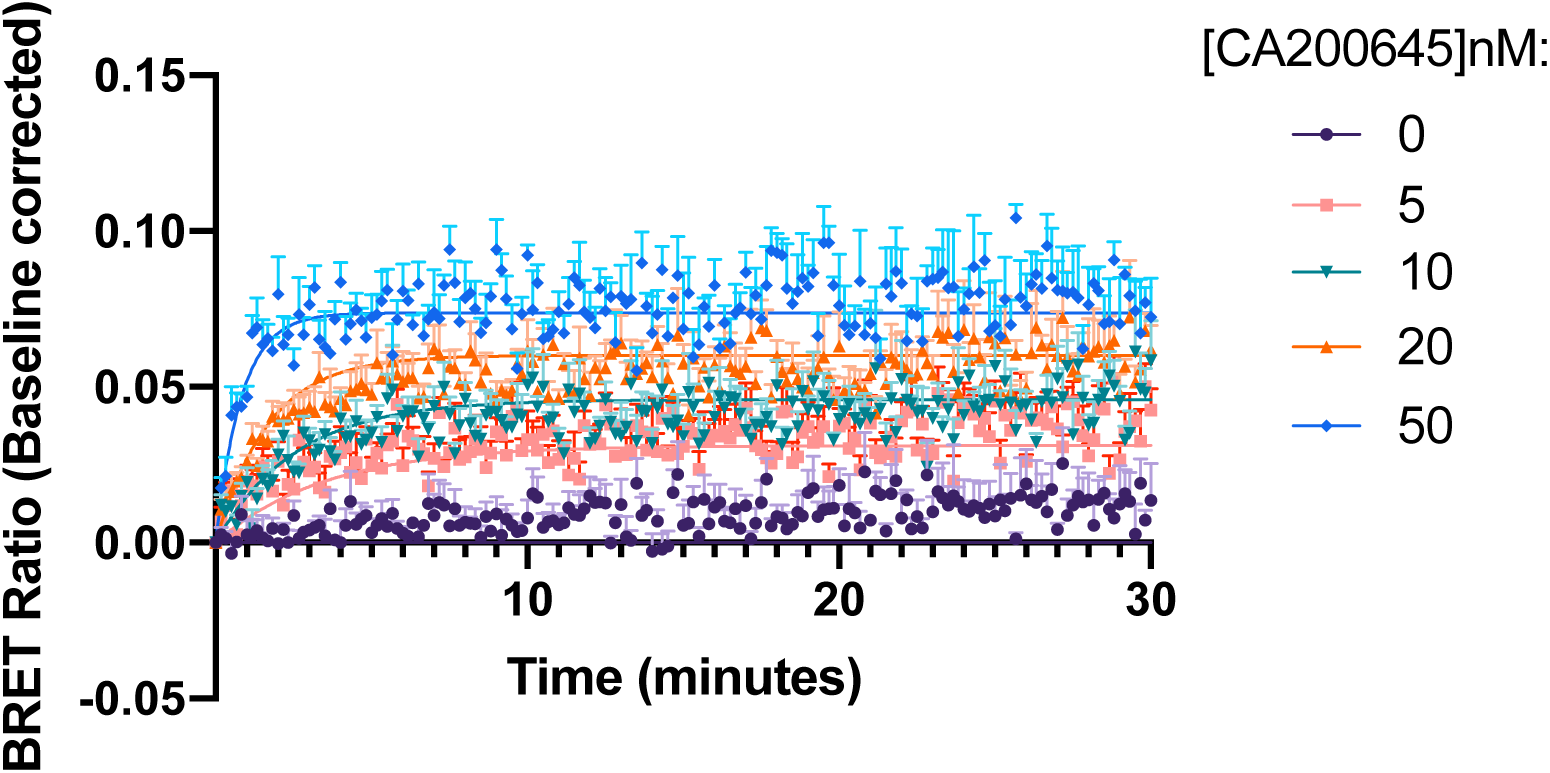
Kinetic measurements of CA200645 binding to Nluc-A_3_R. HEK 293 cells stably expressing Nluc-A_3_R where stimulated with the fluorescent ligand CA200645 at the indicated concentration. BRET between Nluc and the CA200645 was measured every 5 seconds for 30 minutes at room temperature. Determined kinetic parameters for CA200645 at Nluc-A_3_R were K_on_ = 2.86 ± 0.89 × 10^7^ M^-1^ and K_off_ = 0.4397 ± 0.014 min^-1^ with a resulting K_D_ of 17.92 ± 4.45 nM. Data were baseline corrected and shown here as representative of five independent experiments, conducted in duplicate.

**Supplementary Figure 13.**
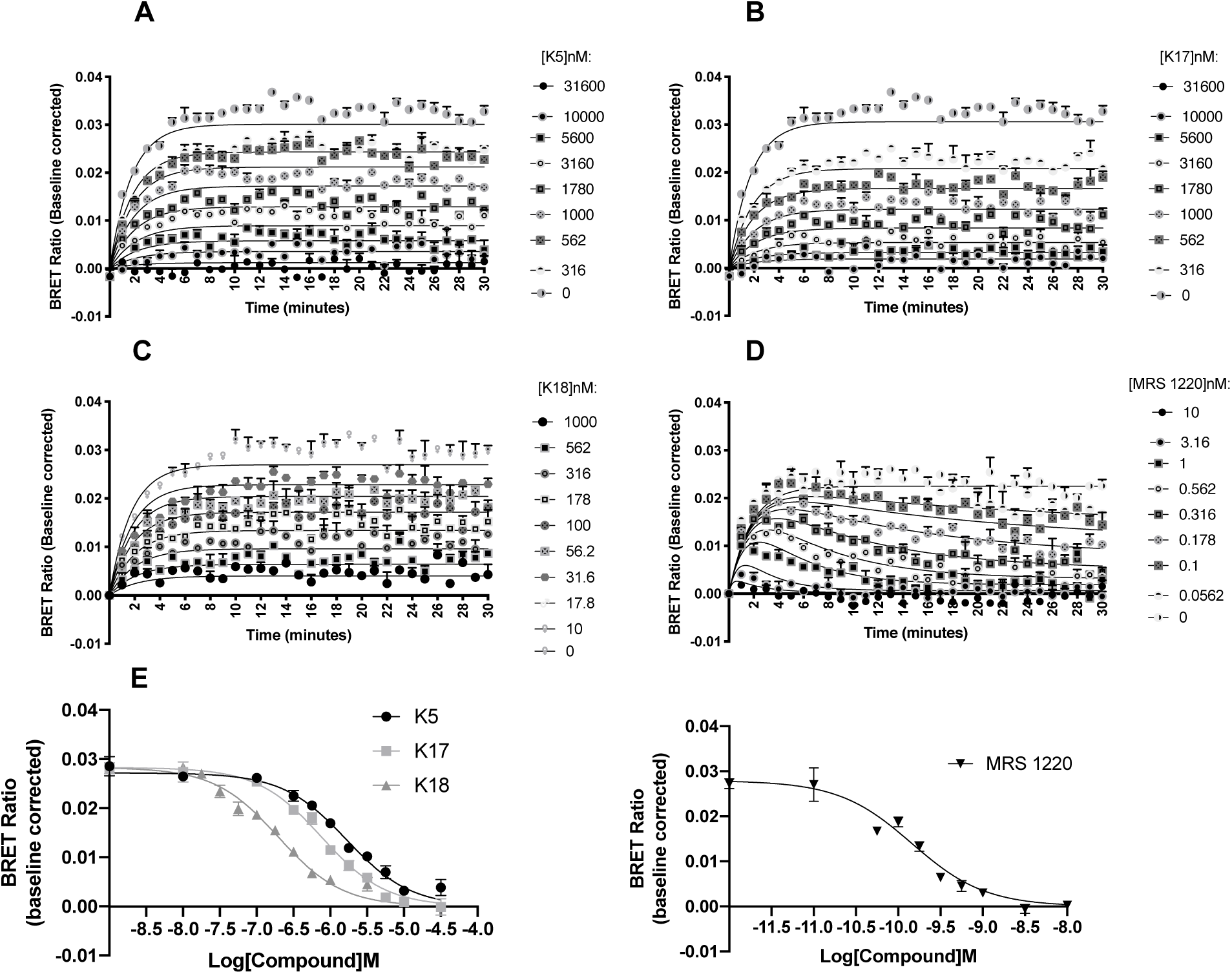
Inhibition of BRET between Nluc and CA200645 at the A_3_R by K5, K17, K18 and MRS 1220. HEK293 cells stably expressing Nluc-A_3_R were treated with 5 nM CA200645 and increasing concentrations of unlabelled compound (represented in nM) **A**) K5, **B**) K17, **C**) K18 or **D**) MRS 1220. For MRS 1220, this trace demonstrates a classic tracer ‘overshoot’, as has been previously described observed when the unlabelled compound has a slower off rate than the labelled CA200645 (K_off_ of 0.4397 ± 0.014 min^-1^ and 0.0248 ± 0.005 min^-1^, respectively) (Sykes *et al.*, 2019, Motulsky and Mahan, 1984; built into Prism). The data shown are representative of three independent experimental repeats (mean ± SEM) fitted with the appropriate model, as determined by statistical comparison between our new model (“Kinetics of competitive binding, rapid competitor dissociation”, derived in the Appendix I) (K5, K17 and K18) or the ‘kinetic of competitive binding’ model (built into Prism) for MRS 1220. (See Materials and Methods for fitting procedure and statistical comparison method.) **E**) The resulting concentration dependent decrease in BRET ratio at 10 minutes was taken to calculate pK_i_ through fitting the Cheng-Prusoff equation. Each data point represents mean ± SEM of five experiments performed in duplicate.

**Supplementary Figure 14.**
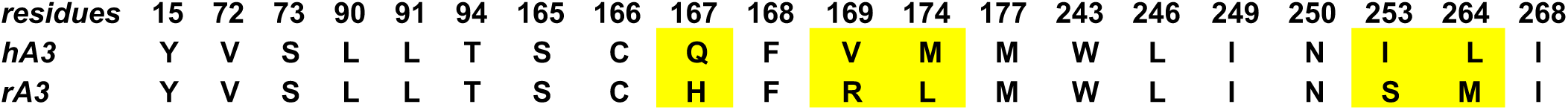
Comparison of the residues of the orthosteric binding area in human and rat A_3_Rs.

**Supplementary Figure 15.**
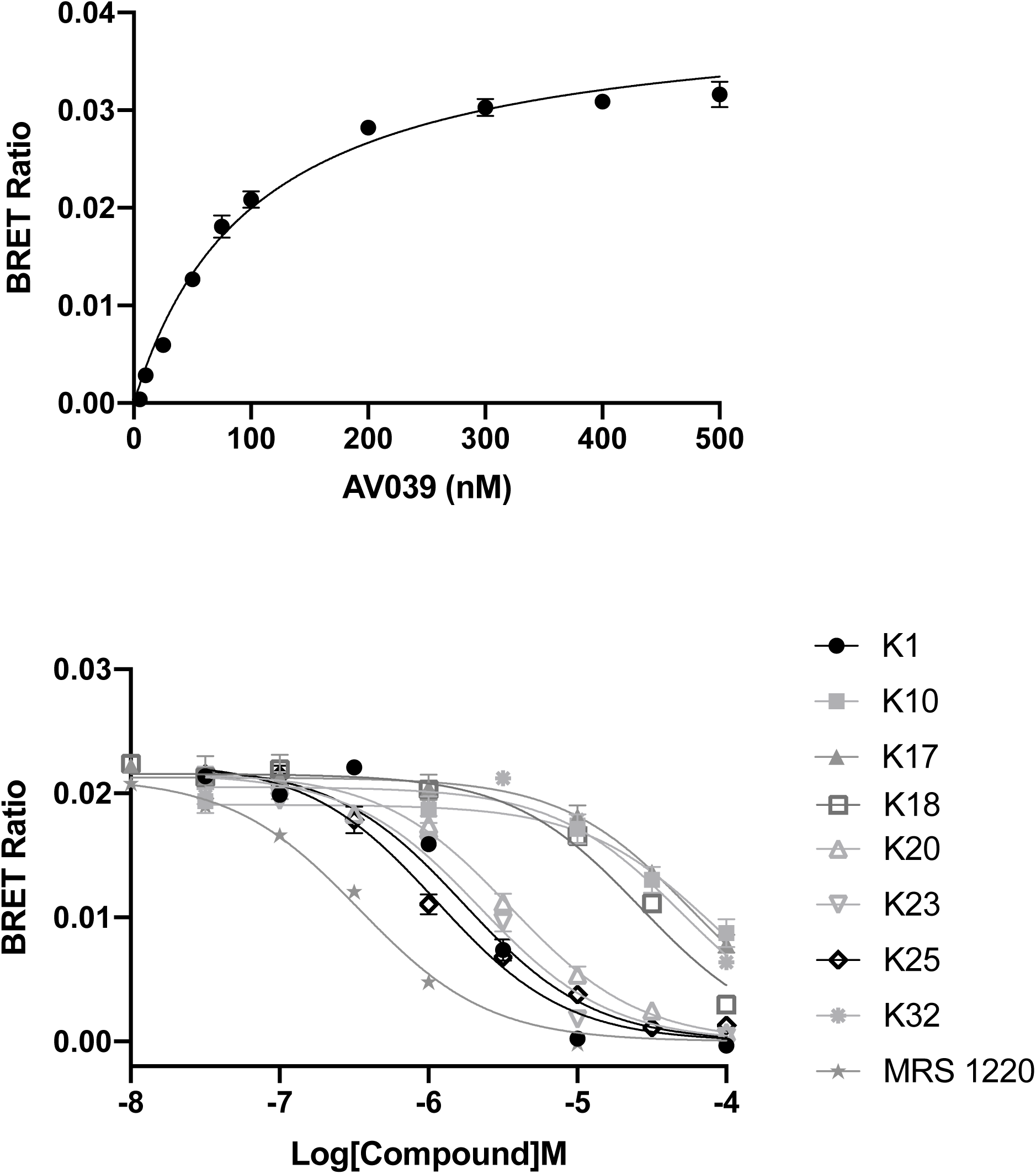
NanoBRET radioligand binding at the rat A_3_R. **A)** Saturation binding experiment with AV039 with a K_D_ determined as 102 ±7.59 nM through fitting the ‘One site – Specific binding’ model in Prism. Each data point represents mean ± SEM of *n = 5* experiments, performed in duplicate. **B)** Inhibition of BRET between Nluc and AV039 at the rat A_3_R by MRS 1220 and K compounds. HEK293 cells stably expressing Nluc-rat A_3_R were treated with 100 nM AV039 and increasing concentrations of unlabelled compound. The resulting concentration dependent decrease in BRET ratio at 5 minutes was taken to calculate pK_i_ through fitting the Cheng-Prusoff equation. Each data point represents mean ± SEM of *n* (*n* =5 for MRS 1220, K1, K20, K23 and K25, *n* =3 for K10, K17, K18 and K32) experiments, performed in duplicate.

**Supplementary Table 7.** Binding of compounds to the rat A_3_R. Equilibrium dissociation constant of MRS 1220 and K compounds as determined through NanoBRET ligand-binding (pK_i_)

| | $pK_i$ | $n$ |
| --- | --- | --- |
| **MRS 1220** | 6.74 ±0.04 | 5 |
| **K1** | 6.07 ±0.05 | 5 |
| **K10** | 4.19 ±0.09 | 3 |
| **K17** | 4.60 ±0.09 | 3 |
| **K18** | 4.60 ±0.04 | 3 |
| **K20** | 5.71 ±0.03 | 5 |
| **K23** | 5.93 ±0.04 | 5 |
| **K25** | 6.37 ±0.06 | 5 |
| **K32** | 4.05 ±0.10 | 3 |

**Supplementary Figure 16.**
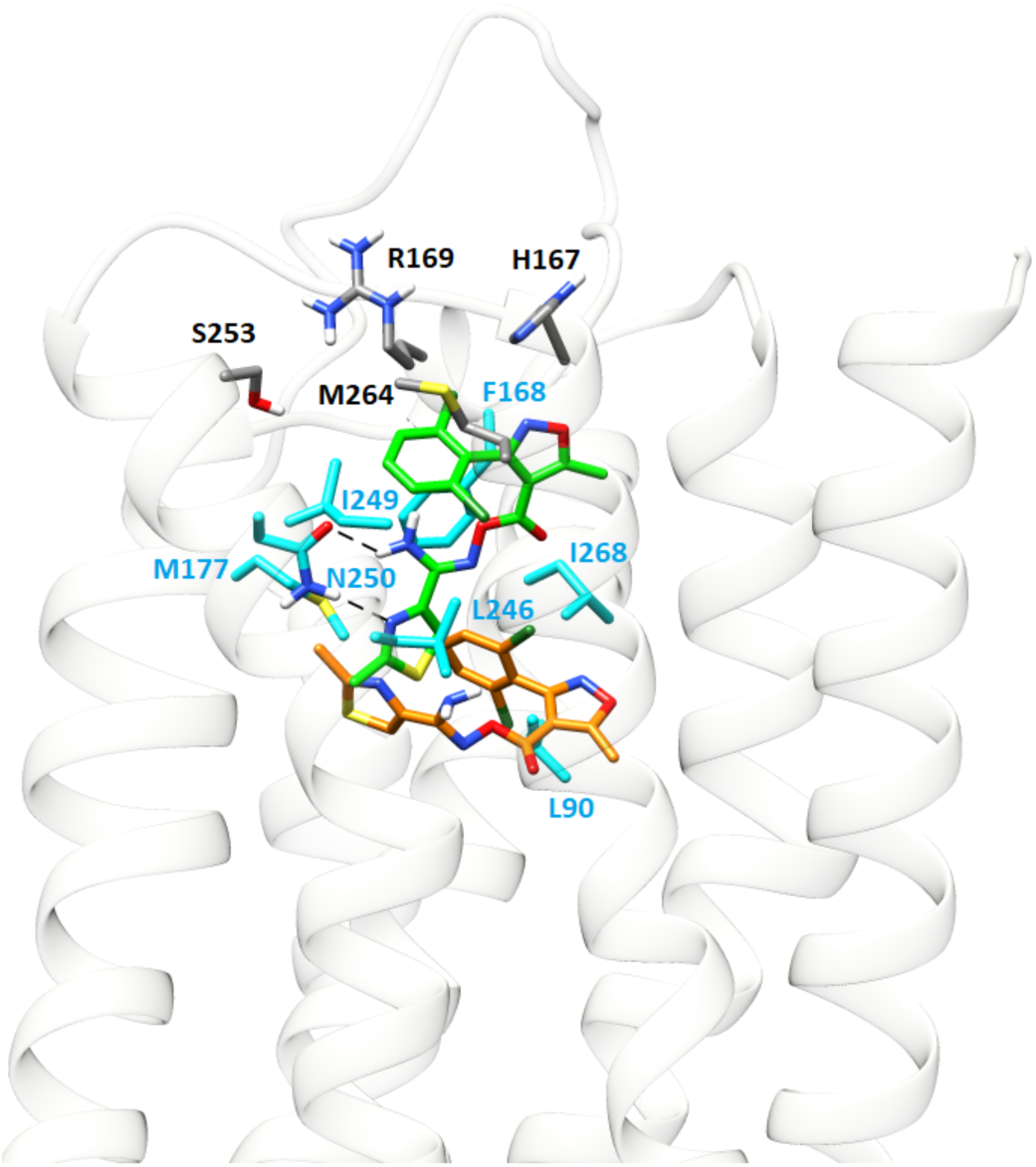
Rat A_3_R in complex with K18. Starting pose (carbons of the ligand in green), after 100 ns MD simulation (carbons of the ligand in orange). Light blue sticks show residues conserved with human A_3_R. M264^7.34^ most likely hampers K18 binding due to steric hindrance of the dichloro-phenyl group

**Supplementary Figure 17.**
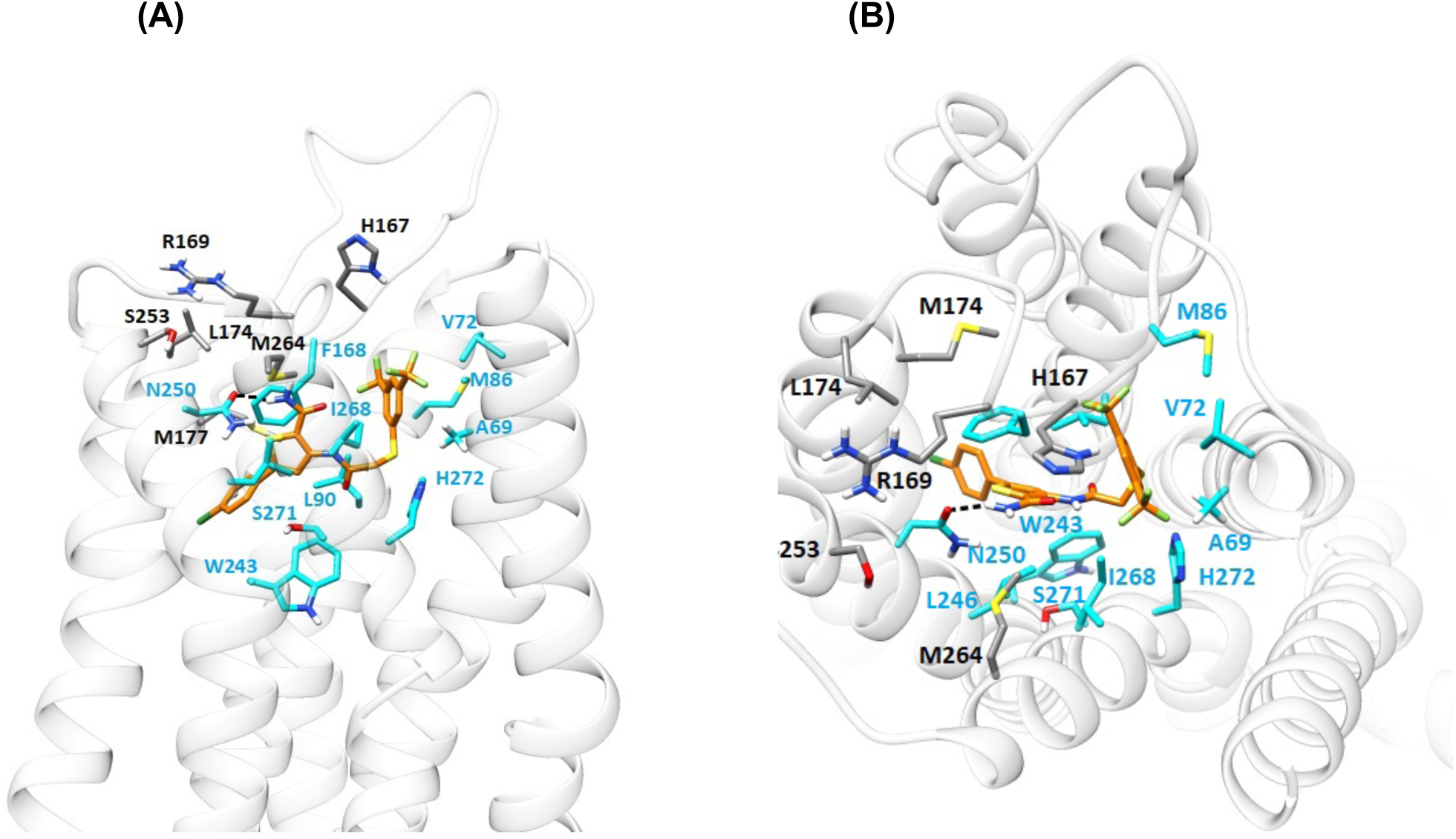
Average structure of rat A_3_R in complex with K25 from 100 ns MD simulations (carbons of the ligand are shown in orange sticks and light blue sticks show residues in contact with K25). K25 was docked into the orthosteric site of the rat A_3_R using the GoldScore scoring function and the highest scoring pose was inserted in a hydrated POPE bilayer. The complexes were subjected to MD simulations with Amber14ff and K25 adopts a potential binding pose within the orthosteric binding area (A) Side view. (B) Top view.

**Supplementary Figure 18.**
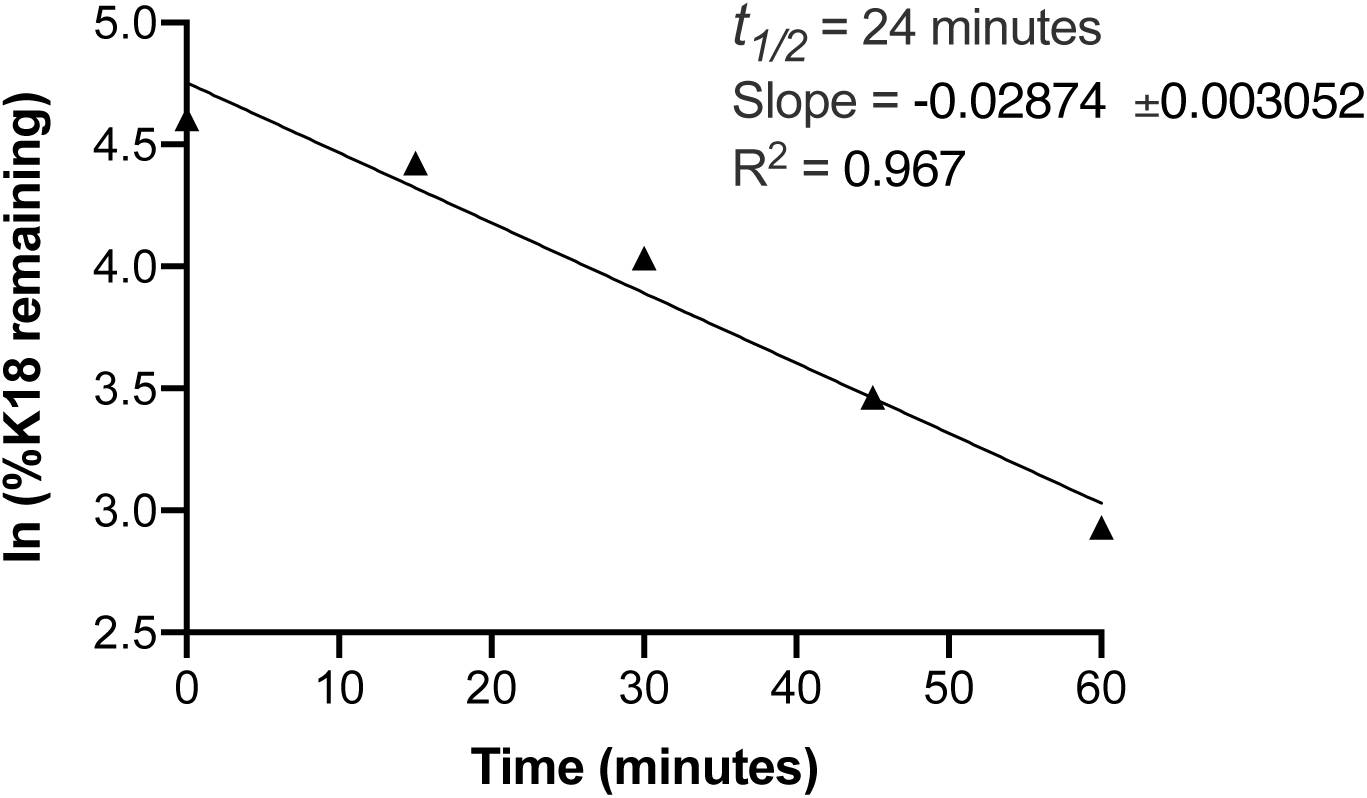
Intrinsic clearance of K18. The metabolic stability of K18 (0.1 μM) was studied using human liver microsomes (0.1 mg/mL) to derive the metabolic half-life (*t*_*1/2*_) from the slope (k). The *t*_*1/2*_ of K18 was determined as 24 minutes using the equation: *t*_*1/2*_ = - In(2)/k. Intrinsic clearance (CL_int_) was calculated as 287.2 μl/min/mg. Each data point represents the mean ± SEM of a single test conducted in duplicate.

## Appendix I: Adapting the kinetics of competitive binding equation for rapidly-dissociating unlabelled ligands

The Motulsky and Mahan equation is used to measure the association and dissociation rate constant of unlabelled compounds, in competition with labelled ligand for binding to a receptor (Motulsky and Mahan, 1984). When the unlabelled competitor dissociates rapidly, relative to the early time points of the assay, the fitted parameters can indicate a failure of the fit to provide realistic or sufficiently precise estimates of the model parameters. Here an equation is derived for rapidly-dissociating compounds that provides an estimate of the equilibrium binding affinity rather than the binding rate constants of the unlabelled competitor. It is assumed the competitor is at equilibrium throughout the time course of the binding assay. This is represented by the following scheme, where *K*_i_ is the equilibrium dissociation constant of the competitor:

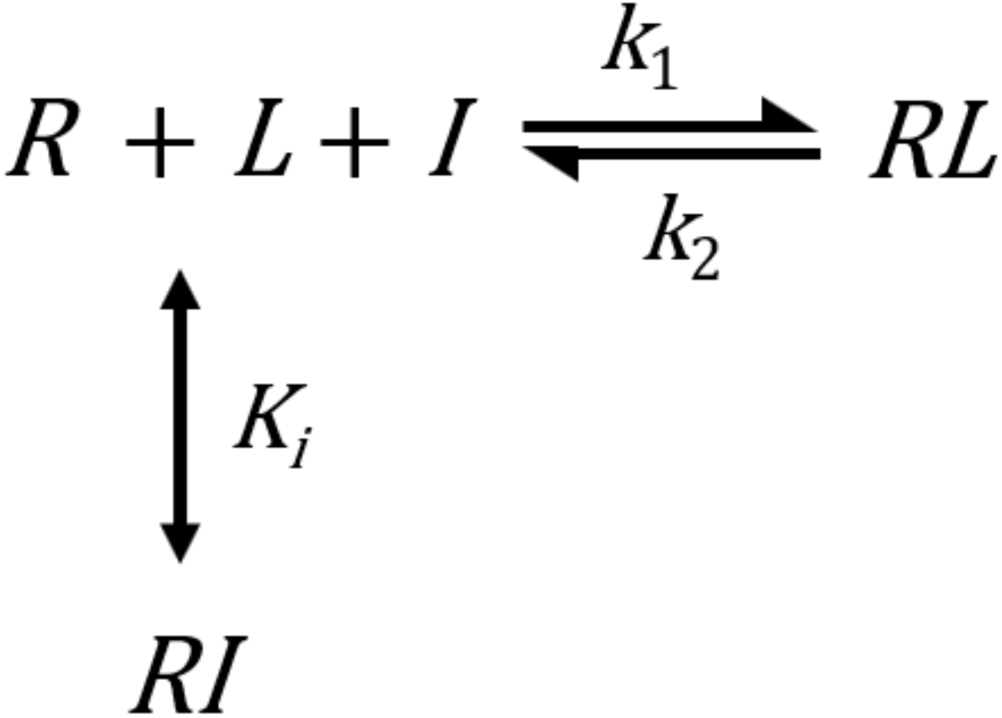

*R* is receptor, *L* is labelled ligand, *I* is unlabelled competitor, *k*_1_ is the association rate constant and *k*_2_ the dissociation rate constant of the labelled ligand. The assumptions underlying the original kinetics of competitive binding equation apply (Motulsky and Mahan, 1984): Only a small fraction (< 10%) of total tracer and compound concentration is bound by receptor (“Zone A”); receptor is exposed simultaneously to both ligands; receptor-ligand binding conforms to a single-site, single-step non-cooperative mass-action mechanism; and unlabelled ligand competitively inhibits tracer binding.

The goal is an equation that describes binding of labelled ligand to receptor over time in the presence of unlabelled competitor. We start with the differential equation for [*RL*]:

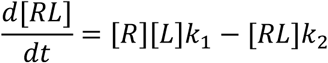

Next, *R* is substituted by employing the conservation of mass equation:

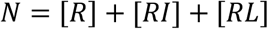

where *N* is the total concentration of receptor. Solving for [*R*] gives,

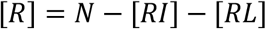

This expression is now substituted into the differential equation for [*RL*]:

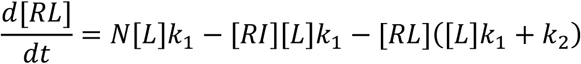

We now substitute [*RI*] with an expression in terms of [*RL*], as follows: Since *I* is at equilibrium with the receptor, the ratio of *R* to *RI* is constant, and can be represented by the following expression:

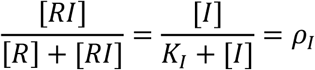

where the constant *ρ*_*I*_ is fractional occupancy by *I* of accessible receptors, i.e. those not bound by labelled ligand. Next, this equation is rearranged as follows:

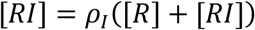

The bracketed term on the right-hand side, [*R*] + [*RI*] can be re-written in terms of [*RL*] and the constant *N*, using the conservation of mass equation for the receptor:

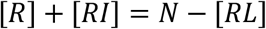

[*R*] + [*RI*] is now substituted with *N* − [*RL*]:

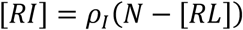

This is the desired expression: [*RI*] is expressed in terms of [*RL*]. This expression is now substituted into the differential equation for [*RL*], which gives, after rearranging,

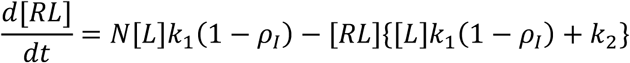

This equation is now integrated to obtain the [*RL*] *vs t* equation that can be used to fit experimental data:

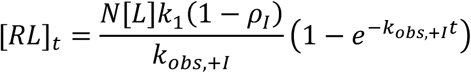

where

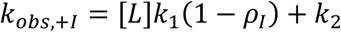

and

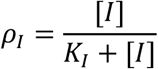

## Appendix II: Supporting information for computational biochemistry

### Preparation of the structures

Structures of the compounds K5, K17, K18 or MRS 1220 were prepared using Maestro (Version 10.5; Schrodinger, Inc.: New York, NY, 2015) and minimized as in a previous paper (Lagarias *et al.*, 2018). The inactive state homologue of A_3_R WT was taken from Adenosiland web-service (Floris *et al.*, 2012). The BLAST algorithm estimated the human A_1_R (Glukhova *et al.*, 2017; PDB ID 5UEN) as the most appropriate template for human A_3_R model having the most similar sequence. The rat A_3_R was generated using the protein structure homology model server SwissModel (Waterhouse *et al.*, 2018). We also applied MODELLER 9.18 (Šali and T. L. Blundell, 1993, Eswar *et al.*, 2003) which selected ten PDB structures with the highest sequential similarity as templates for homology modeling of rat A_3_R. Twenty homology models were generated and the model with the lowest DOPE (Discrete Optimized Protein Energy) value was selected. The two resulting rat A_3_R models created by Swiss Model and MODELLER 9.18 were compared to each other using the Protein Structure Alignment Tool of Desmond Maestro 2018-1 and were found to be very similar (Desmond Molecular Dynamics System, version 3.0; D.E. Shaw Res. New York, 2011; Maest. Interoperability Tools, 3.1; Schrodinger Res. New York, 2012).

The protein models were optimized as previously published using the Protein Preparation Wizard implementation in Schrodinger suite (Protein Prep. Wizard 2015-2; Epik version 2.4, Schrödinger, LLC, New York, NY, 2015; Impact version 5.9, Schrödinger, LLC, New York, NY, 2015; Prime version 3.2, Schrödinger, LLC, New York, NY, 2015). The ZM241385-inactive A_2A_R protein complex from 3EML (Jaakola *et al.*, 2008) was superimposed to human or rat A_3_R WT and the A_2A_R protein was removed resulting in ZM241385-A_3_R complex which was used as a template for docking of K5, K17, K18, K25 or MRS 1220 using GoldScore and ChemPLP scoring functions (Jones *et al.*, 1997, Eldridge *et al.*, 1997) and the GOLD (Version 5.2, Cambridge Crystallogr. Data Cent. Cambridge, U.K., 2015) (Verdonk *et al.*, 2005) as previously described (Lagarias *et al.*, 2018). The top high-scoring poses for K5, K17, K18, K25 or MRS 1220 in complex with A_3_R using GoldScore were better and were kept. These complexes were embedded in POPE bilayers using the System Builder utility of Desmond (Desmond Molecular Dynamics System, version 3.0; D.E. Shaw Res. New York, 2011; Maest. Interoperability Tools, 3.1; Schrodinger Res. New York, 2012). Complex and lipid systems were solvated using the TIP3P water model (Jorgensen, 1983). Na^+^ and Cl^-^ ions were placed in the water phase to neutralize the systems and to reach the experimental salt concentration of 0.150 M NaCl. A 10 Å-from the solute atoms-buffered orthorhombic system with periodic boundary conditions was constructed for all complexes.

### MD simulations

Each ligand-A_3_R complex in the bilayer was processed by the LEaP module in AmberTools14 under the AMBER14 software package (Case *et al.*, 2014). Amber ff14SB force field parameters (Maier *et al*., 2015) was applied to the protein, lipid14 to the lipids (Dickson *et al.*, 2014), GAFF to the ligands (Wang et al., 2004) and TIP3P (Jorgensen, 1983) to the water molecules for the calculation of bonded, vdW parameters and electrostatic interactions. Atomic charges were computed according to the RESP procedure (Bayly *et al.*, 1993) using Gaussian03 (Frisch *et al.*, 2003) and *antechamber* of AmberTools14. MD simulations in explicit solvent were performed using PMEMD (Case *et al.*, 2014). MD simulation protocol consists of five stages: a) Minimization, b) Heating, c) Adjustment of density, d) Equilibration and e) Production. The systems were minimized by 2500 steps of steepest descent to remove bad contacts and 7500 steps of conjugated gradient minimization in the presence of a harmonic restraint with a force constant of 5 kcal mol^-1^ Å^-2^ on all atoms of protein and ligand and non-bonded cut-off of 8.0 Å. The next stage in MD simulation protocol is to allow the system to heat up from 0 K to 310 K. Langevin thermostat (dynamics) (Izaguirre *et al*, 2001) as implemented in Amber14 (Case *et al.*, 2014) was used for temperature control employing a Langevin collision frequency of 2.0 ps^-1^. The system in two consecutive steps to 310 K in the presence of a harmonic restraint with a force constant of 10 kcal mol^-1^ Å^-2^ on all membrane, protein, and ligand atoms. In the first step, systems were heated to 100 K in a NVT of 50 ps length where the adjustment of the density was realized using the Berendsen barostat (Berendsen *et al.*, 1984) with a 2 ps coupling time. In the second step, the temperature was raised to 310 K in a NPT*γ* (with *γ* = 10 dyn cm^-1^) simulation of 500 ps length. Subsequently, the systems were equilibrated without restraints in a NPTγ simulation of 1 ns length with *T* = 310 K and γ = 10 dyn cm^-1^. The equilibration phase was followed by production simulation for 100 ns with system-specific lengths using the same protocol as in the final equilibration step. In the NPTγ simulations semiisotropic pressure scaling to *p* = 1 bar was applied using a pressure relaxation time of 1.0 ps. For the treatment of long-range electrostatic interactions, the Particle-mesh Ewald summation method (Darden *et al.*, 1993, Essmann *et al.*, 1995) was used, and short-range non-bonding interactions were truncated with an 8 Å cutoff. Bonds involving hydrogen atoms were constrained by the SHAKE algorithm (Ryckaert *et al.*, 1977), and a time step of 2 fs was used for the integration of the equations of motion. Snapshots recorded every 20 ps during the production Properties and dynamics of the protein and ligand systems as well as of the membrane were analyzed with the *ptraj* and *cpptraj* modules of AmberTools12 (Case *et al.*, 2014).

